# Revisiting inconsistency in large pharmacogenomic studies

**DOI:** 10.1101/026153

**Authors:** Zhaleh Safikhani, Mark Freeman, Petr Smirnov, Nehme El-Hachem, Adrian She, Rene Quevedo, Anna Goldenberg, Nicolai Juul Birkbak, Christos Hatzis, Leming Shi, Andrew H Beck, Hugo JWL Aerts, John Quackenbush, Benjamin Haibe-Kains

**Affiliations:** Princess Margaret Cancer Centre, University Health Network, Toronto, Ontario, Canada; Department of Medical Biophysics, University of Toronto, Toronto, Ontario, Canada; Institut de recherches cliniques de Montréal, Montreal, Quebec, Canada; Hospital for Sick Children, Toronto, Ontario, Canada; Department of Computer Science, University of Toronto, Toronto, Ontario, Canada; University College London, London, United Kingdom; Section of Medical Oncology, Yale University School of Medicine, New Haven, Connecticut; USA; Yale Cancer Center, Yale University, New Haven, Connecticut, USA; Fudan University, Shanghai City, China; University of Arkansas for Medical Sciences, Little Rock, Arkansas, USA; Department of Pathology, Beth Israel Deaconess Medical Center and Harvard Medical School, Boston, Massachusetts. USA; Department of Cancer Biology, Dana-Farber Cancer Institute, Boston, Massachusetts. USA; Department of Radiation Oncology and Radiology, Dana-Farber Cancer Institute, Brigham and Women’s Hospital, Harvard Medical School, Boston, Massachusetts, USA; Department of Biostatistics and Computational Biology and Center for Cancer Computational Biology, Boston, Massachusetts, USA; Department of Cancer Biology, Dana-Farber Cancer Institute, Boston, Massachusetts, USA

## Abstract

**Background:** In 2012, two large pharmacogenomic studies, the Genomics of Drug Sensitivity in Cancer (GDSC) and Cancer Cell Line Encyclopedia (CCLE), were published, each reported gene expression data and measures of drug response for a large number of drugs and hundreds of cell lines. In 2013, we published a comparative analysis that reported gene expression profiles for the 471 cell lines profiled in both studies and dose response measurements for the 15 drugs characterized in the common cell lines by both studies. While we found good concordance in gene expression profiles, there was substantial inconsistency in the drug responses reported by the GDSC and CCLE projects. Our paper was widely discussed and we received extensive feedback on the comparisons that we performed. This feedback, along with the release of new data, prompted us to revisit our initial analysis. Here we present a new analysis using these expanded data in which we address the most significant suggestions for improvements on our published analysis: that drugs with different response characteristics should have been treated differently, that targeted therapies and broad cytotoxic drugs should have been treated differently in assessing consistency, that consistency of both molecular profiles and drug sensitivity measurements should both be compared across cell lines to accurately assess differences in the studies, that we missed some biomarkers that are consistent between studies, and that the software analysis tools we provided with our analysis should have been easier to run, particularly as the GDSC and CCLE released additional data.

**Methods:** For each drug, we used published sensitivity data from the GDSC and CCLE to separately estimate drug dose-response curves. We then used two statistics, the area between drug dose-response curves (ABC) and the Matthews correlation coefficient (MCC), to robustly estimate the consistency of continuous and discrete drug sensitivity measures, respectively. We also used recently released RNA-seq data together with previously published gene expression microarray data to assess inter-platform reproducibility of cell line gene expression profiles.

**Results:** This re-analysis supports our previous finding that gene expression data are significantly more consistent than drug sensitivity measurements. The use of new statistics to assess data consistency allowed us to identify two broad effect drugs — 17-AAG and PD-0332901 — and three targeted drugs — PLX4720, nilotinib and crizotinib — with moderate to good consistency in drug sensitivity data between GDSC and CCLE. Not enough sensitive cell lines were screened in both studies to robustly assess consistency for three other targeted drugs, PHA-665752, erlotinib, and sorafenib. Concurring with our published results, we found evidence of inconsistencies in pharmacological phenotypes for the remaining eight drugs. Further, to discover “consistency” between studies required the use of multiple statistics and the selection of specific measures on a case-by-case basis.

**Conclusion:** Our results reaffirm our initial findings of an inconsistency in drug sensitivity measures for eight of fifteen drugs screened both in GDSC and CCLE, irrespective of which statistical metric was used to assess correlation. Taken together, our findings suggest that the phenotypic data on drug response in the GDSC and CCLE continue to present challenges for robust biomarker discovery. This re-analysis provides additional support for the argument that experimental standardization and validation of pharmacogenomic response will be necessary to advance the broad use of large pharmacogenomic screens.

## SUMMARY BOX

In 2013 we reported inconsistency in the drug sensitivity phenotypes measured by the Genomics of Drug Sensitivity in Cancer (GDSC) and the Cancer Cell Lines Encyclopedia (CCLE) studies. Here we revisit that analysis and address a number of potential concerns raised about our initial methodology:

- ***Different drugs should be compared based on the observed pattern of response.*** To address this concern, we considered drugs falling into three classes: (1) drugs with no observed activity in any of the cell lines; (2) drugs with sensitivity observed for only a small subset of cell lines; and (3) drugs producing a response in a large number of cell lines. For each class, we assessed the correlation in drug response between studies using a variety of metrics, selecting the metric that performed best in each individual comparison. While no metric identified any substantial consistency for the first class (sorafenib, erlotinib, and PHA-665752), judicious choice of metric found high consistency for three of eight highly targeted therapies in the second class (nilotinib, crizotinib, and PLX4720), but no metric found better than moderate correlation for two of four broad effect drugs in the third class (PD-0332901 and 17-AAG).
- ***Measure of consistency for targeted drugs***. Beyond considering drug response profiles, targeted drugs should be treated differently when assessing consistency. We used six different statistics to test consistency, using both continuous and discretized drug sensitivity data. We confirmed that Spearman rank correlation, used in our 2013 study, does not detect consistency for the three highly targeted therapies profiled by GDSC and CCLE. Other statistics, such as Somers’ Dxy or Matthews correlation coefficient, yielded moderate to high consistency for specific drugs, but there was no single metric that found good consistency for each of the targeted drugs.
- ***Consistency of molecular profiles across cell lines***. In our initial published analysis, we reported correlations based on comparing drug response “across cell lines” while gene expression levels were compared “between cell lines.” It has been suggested it would be more appropriate to compute correlations “across cell lines” for both molecular and pharmacological data. Here we report a number of statistical measures of consistency for both gene expression and drug response compared across cell lines and confirm our initial finding that gene expression is significantly more consistent than the reported drug phenotypes.
- ***Some published biomarkers are reproducible between studies***. In our initial comparative study we found that the majority of known biomarkers predictive of drugs response are reproducible across studies. We extended the list of known biomarkers and found that seven out of eleven are significant in GDSC and CCLE. While one can find such anecdotal examples, they do not lead to a general process for discovering a new biomarker in one study that can be applied to another study.
- ***Research reproducibility***. The code we provided with our original paper was incompatible with updated releases of the GDSC and CCLE datasets. We developed *PharmacoGx*, which is a flexible, open-source software package based on the statistical language R, and used it to derive the results reported here.

## INTRODUCTION

The goal of precision medicine is identification of the best therapy for each patient and their own unique manifestation of a disease. This is particularly important in oncology where multiple cytotoxic and targeted drugs are available, but their therapeutic benefits are often insufficient or limited to a subset of cancer patients. Large-scale pharmacogenomics studies in which drug and drug candidates are screened against panels of molecularly characterized cancer cell lines, have been proposed as a means for identifying drugs effective against specific cancers and for developing predictive genomic biomarkers of drug response. The Genomics of Drug Sensitivity in Cancer project (GDSC, referred to as the Cancer Genome Project [CGP] in our initial study) ^1^, and the Cancer Cell Line Encyclopedia (CCLE) ^2^ have each reported results of such screens, providing data on drug sensitivities and molecular profiles for collections of representative cancer cell lines.

Presented with these two large studies, our hope was that we could use the data to identify new gene expression biomarkers of drug response in one study that would predict response in the second. To our surprise, we were unable to find such biomarkers for many drugs, even when we limited our analysis to the drugs and cell lines screened in common by the GDSC and CCLE. There have since been a number of published studies that have reported difficulties in building and validating biomarkers of response using these two datasets ^3–6^.

To understand the cause of this failure, we compared the gene expression profiles and the drug response data reported by the GDSC and CCLE ^7,8^. We found that, although the gene expression data showed reasonable consistency between the two studies, the drug sensitivity measurements were surprisingly inconsistent. This inconsistency can be clearly seen by plotting drug response reported for each of fifteen drugs provided in both GDSC and CCLE for the 471 cell lines assayed by both studies ^7,8^. Since the publication of our comparative analysis, we received a great deal of constructive feedback from the scientific community regarding multiple aspects of the analysis we reported, including suggestions for analytical methods that might uncover greater consistency between the studies. We were also fortunate that both GDSC and CCLE have released new drug sensitivity and gene expression data, allowing us not only to revisit our initial analysis, but also to extend it using these new data.

To begin, we investigated alternative statistics to assess the inter-study consistency for drugs exhibiting different patterns of response across the collection of cell lines common to both studies. We then considered statistical methods for highly targeted drugs expected to be sensitive only in a subset of cell lines. We compared consistency estimates between continuous and discrete gene expression and drug sensitivity data, and importantly, assessed how potential discordance may affect the discovery of molecular features (biomarkers) predictive of drug response. We also revisited our analysis of consistency in gene expression levels between studies and evaluated “known biomarkers” of response expected to be predictive in these studies.

This extensive reanalysis found that by selecting specific statistical measures on a case-by-case basis, one can identify moderate to good consistency for two broad effect and three highly targeted therapies. However, overall our results support our initial observations that drug sensitivity data in GDSC and CCLE are inconsistent for the majority of the drugs, even when considering metrics yielding the highest consistency for individual drugs. Our present analysis adds further evidence supporting the need for robust and standardized experimental pipelines to assure generation of comparable, biologically relevant measures of drug response as well as unbiased statistical and machine learning methods to better predict response. Failure to do so will continue to limit the potential for use of large-scale pharmacogenomic screens in reliable drug development and precision medicine applications.

## RESULTS

The overall analysis design of our study is represented in Figure 1.

**Figure 1:**
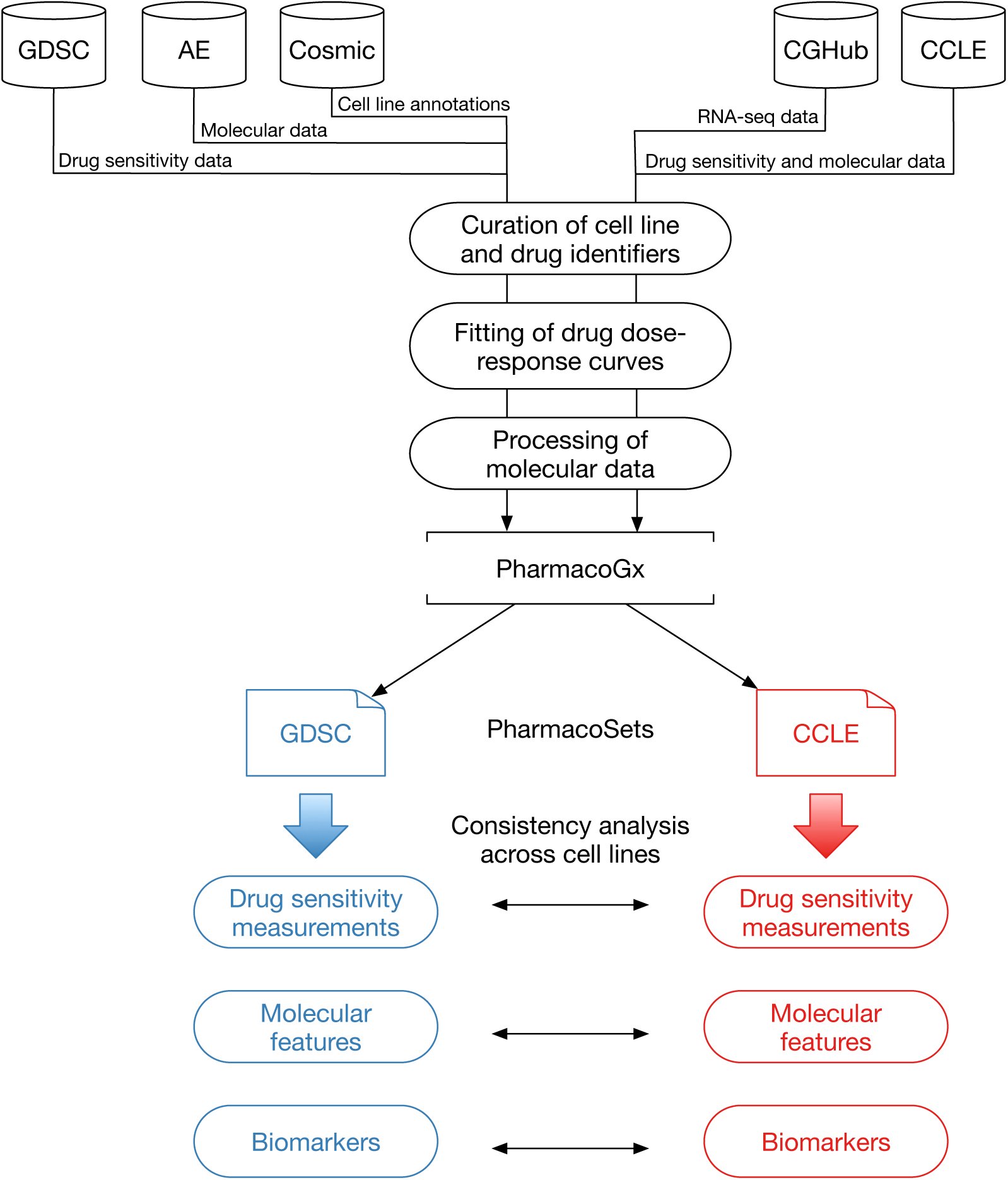
Analysis design. GDSC: Genomics of Drug Sensitivity in Cancer; AE: ArrayExpress; Cosmic: Catalogue of Somatic Mutations in Cancer; CGHub: Cancer Genomics Hub; CCLE: Cancer Cell Line Encyclopedia.

### Intersection between GDSC and CCLE

To identify the largest set of cell lines and drugs profiled by both GDSC and CCLE, we used the *PharmacoGx* computational platform ^9^ that is able to store, analyze, and compare curated pharmacogenomic datasets. We created new datasets for the new releases of the GDSC (June 2014 and July 2015 for drug sensitivity and gene expression data, respectively) and CCLE (February 2015) projects. The improved curation of new data using *PharmacoGx* identified 15 drugs in common between GDSC and CCLE as well 698 cell lines, originating from 23 tissue types (Supplementary Figure 1). This is the same number of shared drugs but the updated datasets contains a larger number of common cell lines than the 471 reported in our previous analysis ^7^.

### Comparing single nucleotide polymorphism (SNP) fingerprints

To check the accuracy of cell line name matching, we compared single nucleotide polymorphism (SNP) fingerprints using data released in both studies. We first controlled for the quality of the SNP arrays and excluded eleven of 1,396 profiles due to low quality (see Methods). We then compared SNP fingerprints of cell lines with identical name using > 80% as threshold for concordance. Consistent with the results reported by the CCLE ^2^, the vast majority of cell lines had highly concordant fingerprints (462 out of 470 cell lines with SNP profiles available in both GDSC and CCLE; Supplementary File 1). Using 80% genomic identity as a cutoff ^2,10^, we found eight cell lines with same identifier but different SNP identity (Figure 2); these were removed from our subsequent analyses to avoid discrepancies due to the use of possibly mislabeled or contaminated cell lines.

**Figure 2:**
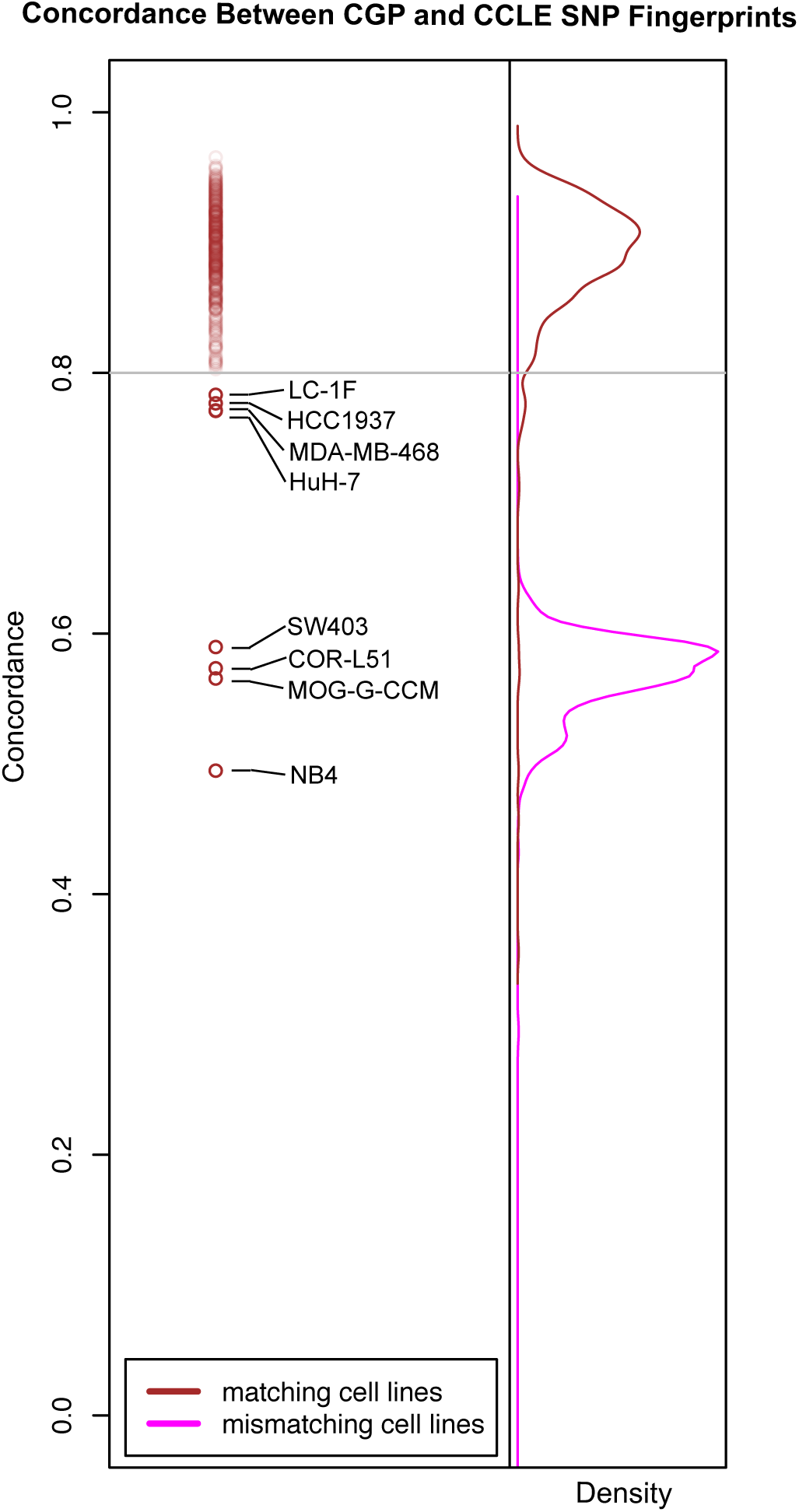
SNP fingerprinting between cancer cell lines screened in GDSC and CCLE.

### Estimation and filtering of drug dose-response curves

We used the recent release of drug dose-response data from GDSC and CCLE to fit dose-response curves and assess their quality. An important factor influencing the fitting of drug dose-response curves is the range of concentration used for each cell line/drug combination. In CCLE, all dose-response curves were measured at eight concentrations: 2.5□10^-3^, 8□10^-3^, 2.5□10^-2^, 8□10^-2^, 2.5□10^-1^, 8□10^-1^, 2.5, and 8 μM. However, in GDSC response was measured at a different set of concentrations for each drug. The minimum concentrations for different drugs range from 3.125□10^-5^ to 15.625 μM. In each case, the concentrations tested by GDSC form a geometric sequence of nine terms with a common ratio of two between successive concentrations. Thus, the maximum concentration tested for each drug is 256 times the minimum concentration for that drug and ranges from 8□10^-3^ to 4000 μM.

To properly fit drug dose-response curves, one must make multiple assumptions regarding the cell viability measurements generated by the pharmacological platform used in a given study. For instance, one assumes that viability ranges between 0% and 100% after data normalization and that consecutive viability measurements remain stable or decrease monotonically reflecting response to the drug being tested. Quality controls were implemented to flag dose-response curves that strongly violate these assumptions (Supplementary Methods). We identified 2315 (2.9%) and 123 (1%) dose-response curves that failed to pass in GDSC and CCLE, respectively, as exemplified in Figure 3 (all noisy curves are provided in Supplementary File 2). We excluded these cases to avoid erroneous curve fitting.

**Figure 3:**
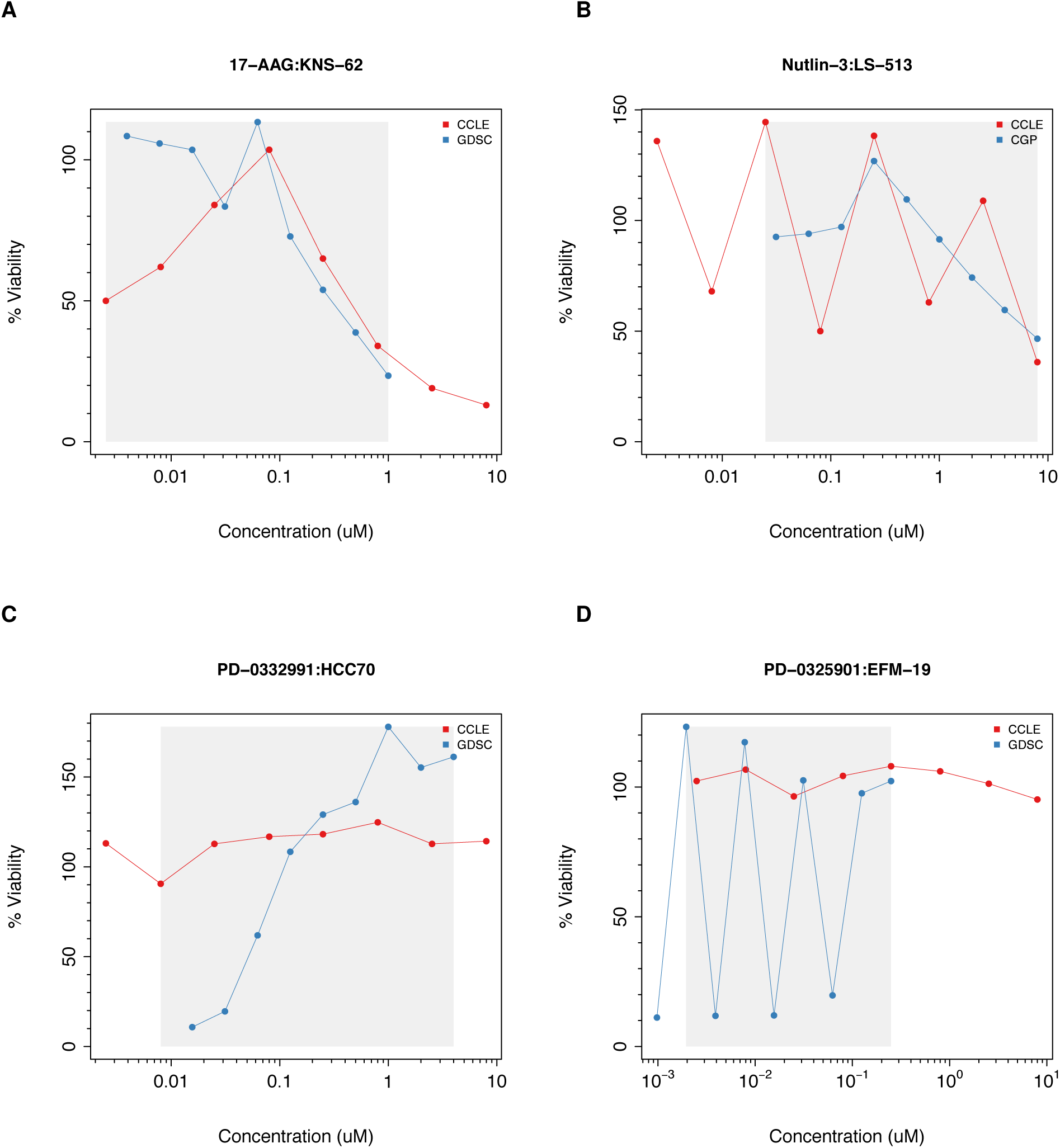
Examples of noisy drug dose-response curves identified during the filtering process in GDSC and CCLE. The grey area represents the common concentration range between studies.

We used least squares optimization to fit a three-parameter sigmoid model (Methods) for the drug dose-response curves in GDSC and CCLE (Supplementary File 3). For each fitted curve, we computed the most widely used drug activity metrics, that are the area under the curve (AUC) and the drug concentration required to inhibit 50% of cell viability (IC_50_).

### Consistency of drug sensitivity data

We began by computing the area between the two drug dose-response curves (ABC) to assess consistency of cell viability data for each cell line combination screened in both GDSC and CCLE using the common concentration range. ABC measures the difference between two drug-dose response curves by estimating the absolute area between these curves, which ranges from 0% (perfect consistency) to 100% (perfect inconsistency). The ABC statistic identified highly consistent (Figure 4A,B) and highly inconsistent (Figure 4C,D) dose-response curves between GDSC and CCLE. The mean of the ABC estimates for all drug-cell line combinations was 10% (Supplementary Figure 2A), with PD-0332991 yielding the highest discrepancies (Supplementary Figure 2B).

**Figure 4:**
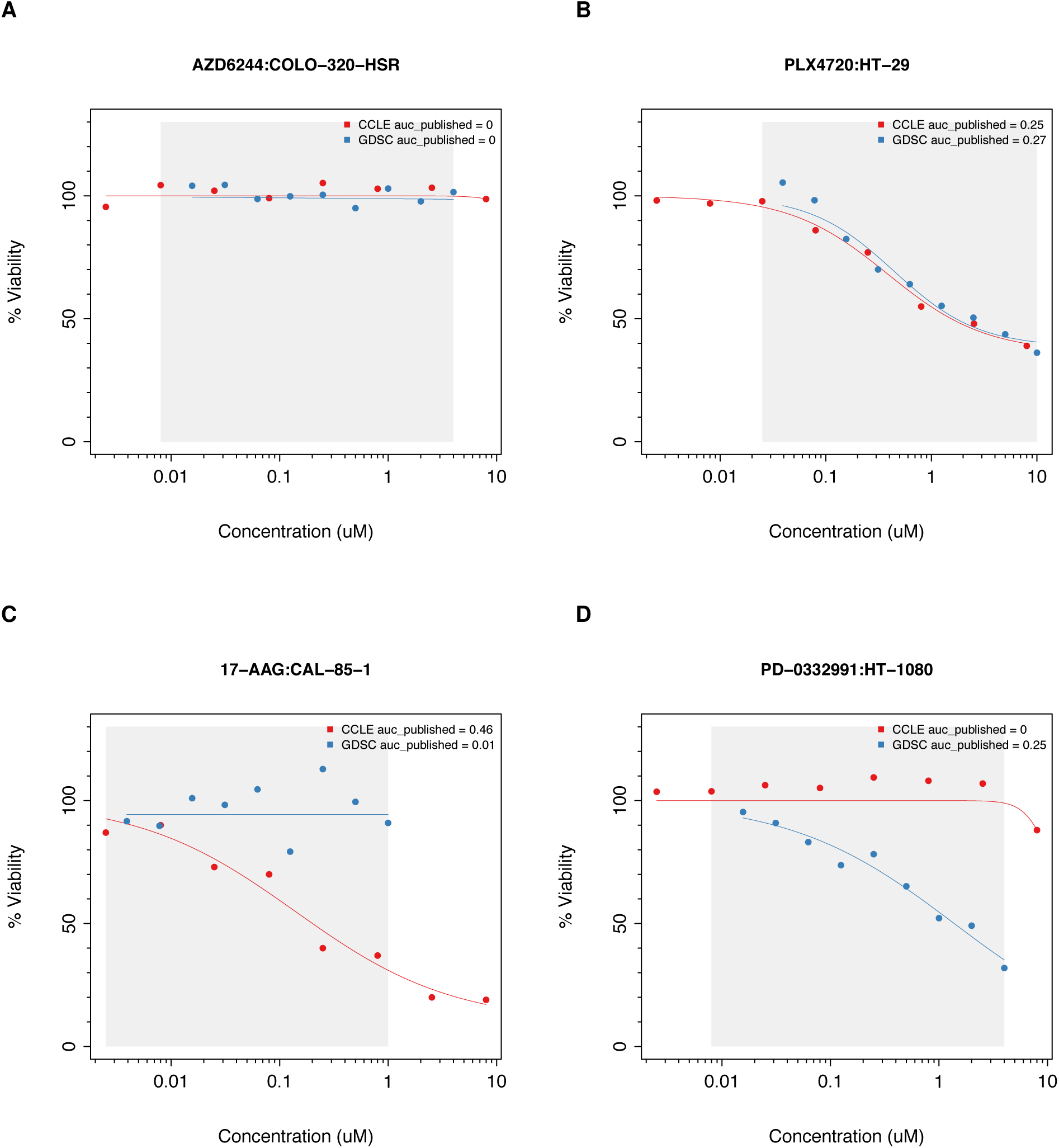
Examples of (A,B) consistent and (C,D) inconsistent drug dose-response curves in GDSC and CCLE. The grey area represents the common concentration range between studies.

We compared biological replicates in GDSC, which were performed independently at the Massachusetts General Hospital (MGH) and the Wellcome Trust Sanger Institute (WTSI). These experiments are comprised of 577 cell lines treated with AZD6482, a PI3Kβ inhibitor screened in GDSC (Supplementary File 4). We computed the ABC of these biological replicates and observed both highly consistent and inconsistent cases (Supplementary Figure 3). We then computed the median ABC values for each pair of drugs in GDSC and used these as a distance metric for complete linkage hierarchical clustering. We found that the MGH-and WTSI-administered AZD6482 experiments clustered together, suggesting that the differences between dose-response curves of biological replicates were smaller than the differences observed between different drugs (Supplementary Figure 4A). We performed the same clustering analysis by computing the ABC-based distance between all the drugs in GDSC and CCLE and observed that only three out of the fifteen common drugs clustered tightly (17-AAG, lapatinib, and PHA-665752; Supplementary Figure 4B). Despite the small number of cell lines exhibiting sensitivity to PHA-665752 and lapatinib, these drugs closely clustered between GDSC and CCLE; however this was not the case for other highly targeted therapies, such as AZD0530, nilotinib, crizotinib and TAE684 Supplementary Figure 4B).

Although the ABC values provide a measure of the degree of consistency between studies, it is the AUC and IC_50_ estimates, and their correlation with molecular features (such as mutational status and gene expression) that are commonly used to assess drug response. Therefore we revisited our comparative analysis of the drug sensitivity data using the expanded data now available and the standardized methods implemented in our *PharmacoGx* platform. Using the same three-parameter sigmoid model to fit drug dose-response curves in GDSC and CCLE (see Methods), we recomputed AUC and IC_50_ values and observed very high correlation between published and recomputed drug sensitivity values for each study individually (Spearman `ρ` > 0.93; Supplementary Figure 5).

**Figure 5:**
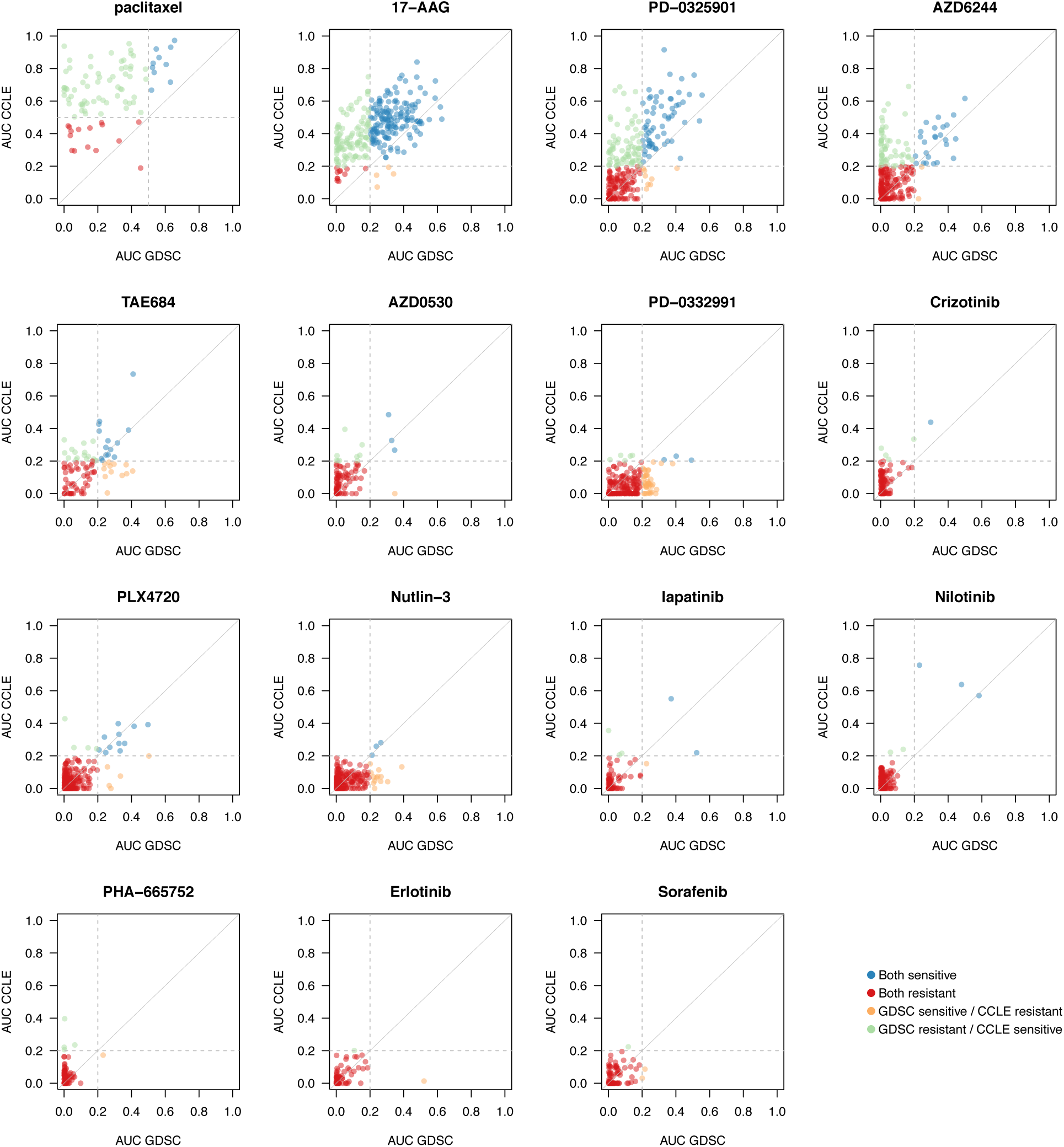
Comparison of AUC values as published in GDSC and CCLE. For cytotoxic drugs (paclitaxel), cell lines with AUC *<* 0.4 were considered as resistant, while for targeted therapies cell lines with AUC < 0.2 were considered resistant (grey dashed lines). In case of perfect consistency, all points would lie on the grey diagonal.

It has been suggested that some of the observed inconsistencies between the GDSC and CCLE may be due to the nature of targeted therapies, which are expected to have selective activity against some cell lines ^11^. This is not an unreasonable assumption as the measured response in resistant cell lines may represent random technical noise that one should not expect to be correlated between experiments. We therefore decided to clearly discriminate between highly targeted drugs with narrow growth inhibition effects and drugs with broader effects. We used the full GDSC and CCLE datasets to compare the variation of the drug sensitivity data of known targeted and cytotoxic therapies as classified in the original studies (Supplementary Figure 6). We observed that drugs can be classified in these two categories based on median absolute deviation (MAD) of the estimated AUC values (Youden’s optimal cutoff ^12^ of AUC MAD > 0.13 for cytotoxic drugs). We then used this cutoff on the common drug-cell line combinations in GDSC and CCLE to define three classes of drugs (Supplementary Figure 7):

**Figure 6:**
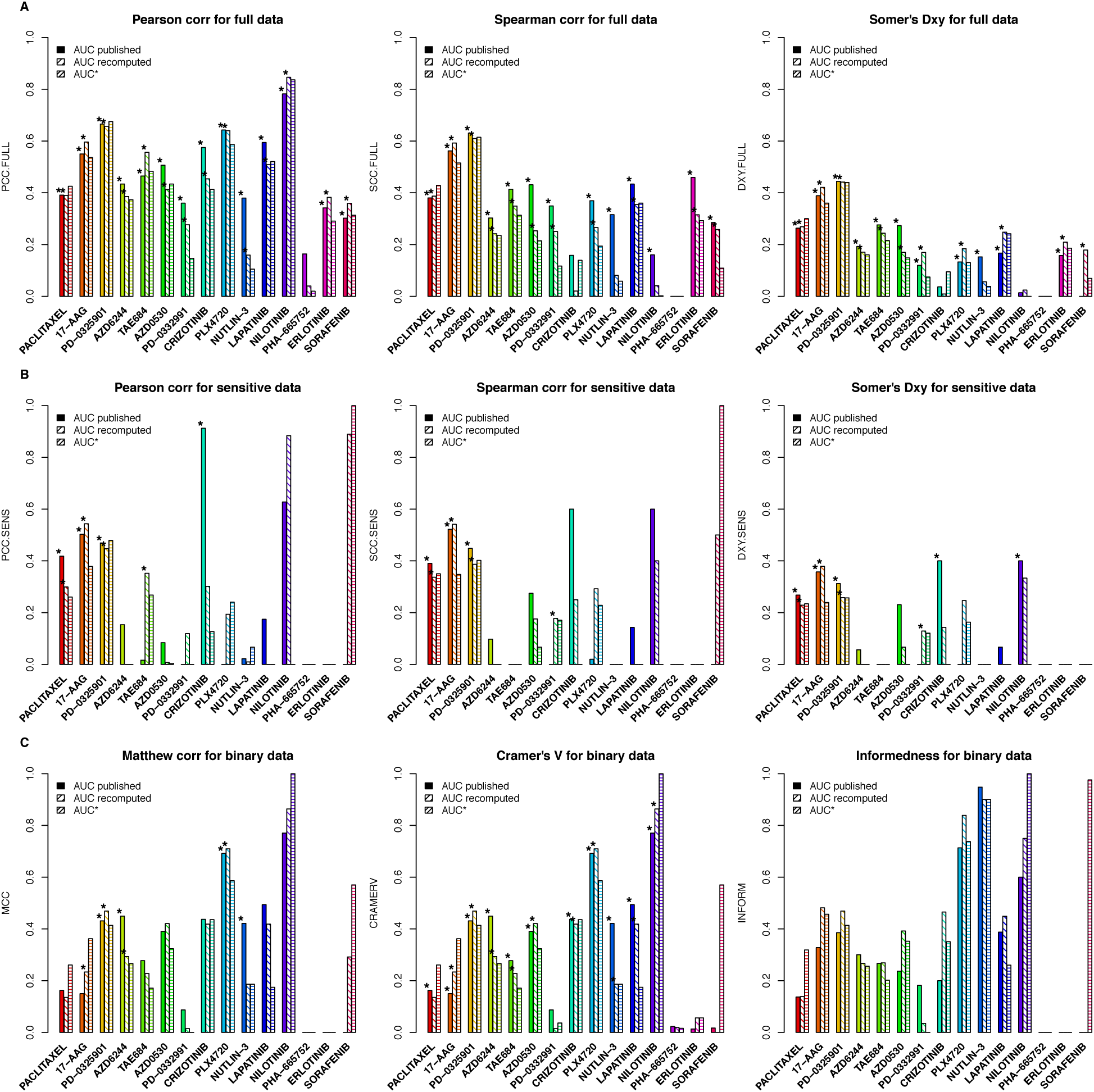
Consistency of AUC values as published and recomputed within *PharmacoGx*. AUC* is the AUC value calculated by considering only the common concentration range between GDSC and CCLE.

**Figure 7:**
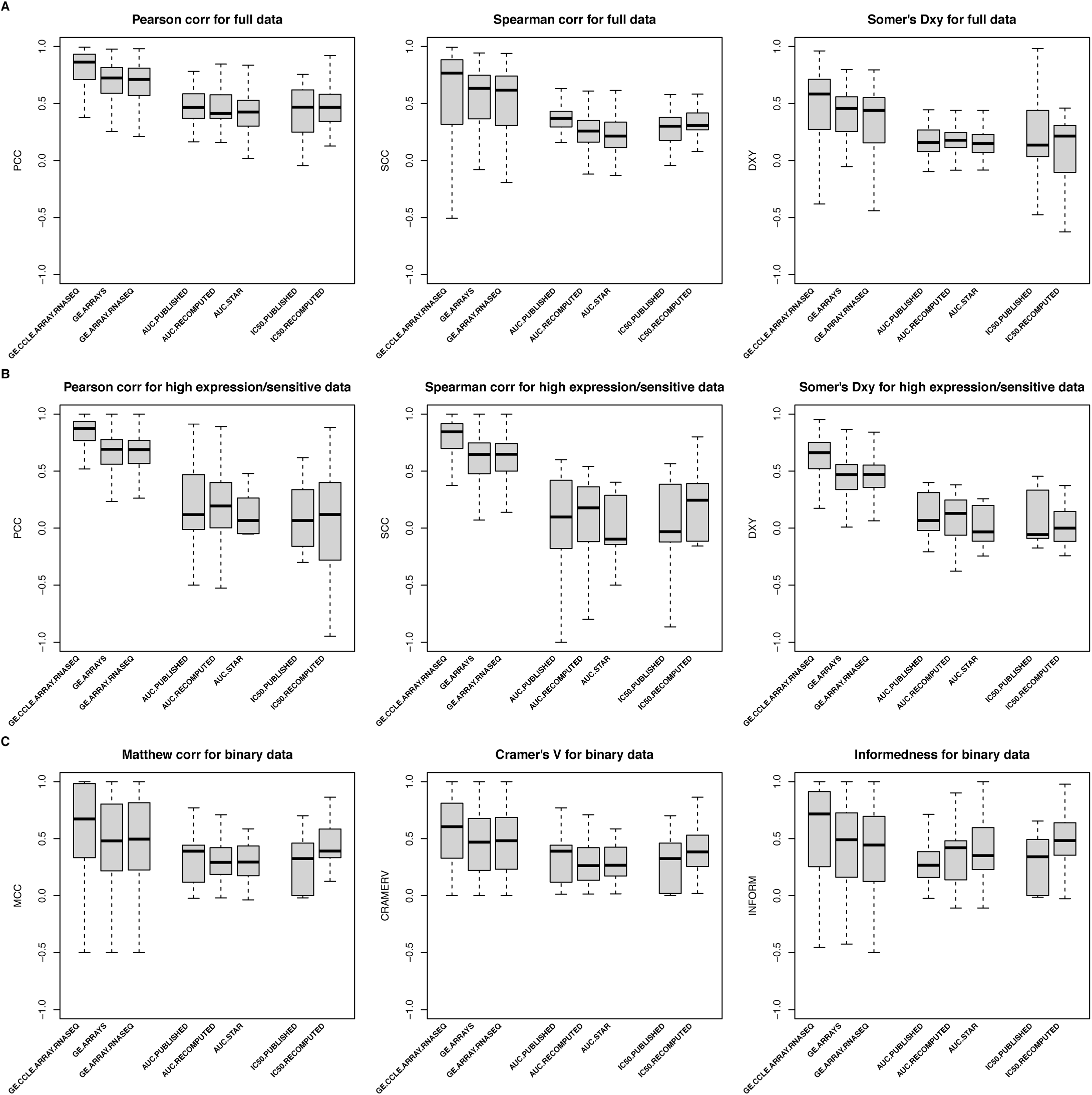
Consistency of gene expression and drug sensitivity data between GDSC and CCLE using multiple consistency measures.

- **No effect**: Drugs with minimal observed activity (typically active in less than 5 “non-resistant” cell lines with AUC > 0.2 or IC_50_ < 1 μM in either study). This class includes sorafenib, erlotinib and PHA-665752.
- **Narrow effect**: Highly targeted drugs with activity observed for only a small subset of cell lines (AUC MAD ≤ 0.13). This group includes nilotinib, lapatinib, nutlin-3, PLX4720, crizotinib, PD-0332991, AZD0530, and TAE684.
- **Broad effect:** Drugs producing a response in a large number of cell lines (AUC MAD > 0.13). This includes AZD6244, PD-0325901, 17-AAG and paclitaxel.

We then compared the AUC (Figure 5, Supplementary Figures 8 and 9 for published AUC, recomputed AUC and AUC computed based on the common concentration range, respectively) and IC_50_ (Supplementary Figures 10 and 11) values and calculated the consistency of drug sensitivity data between studies using all common cases and only those that the data suggested were sensitive in at least one study (Figures 6 and Supplementary Figure 12 for AUC and IC_50_, respectively, and Supplementary Tables 1-5). Given that no single metric can capture all forms of consistency, we extended our previous study by using the Pearson correlation ^13^, Spearman ^14^, and Somers’ Dxy ^15^ rank correlation coefficients to quantify the consistency of continuous drug sensitivity measurements across studies (see Methods).

**Table 1:**
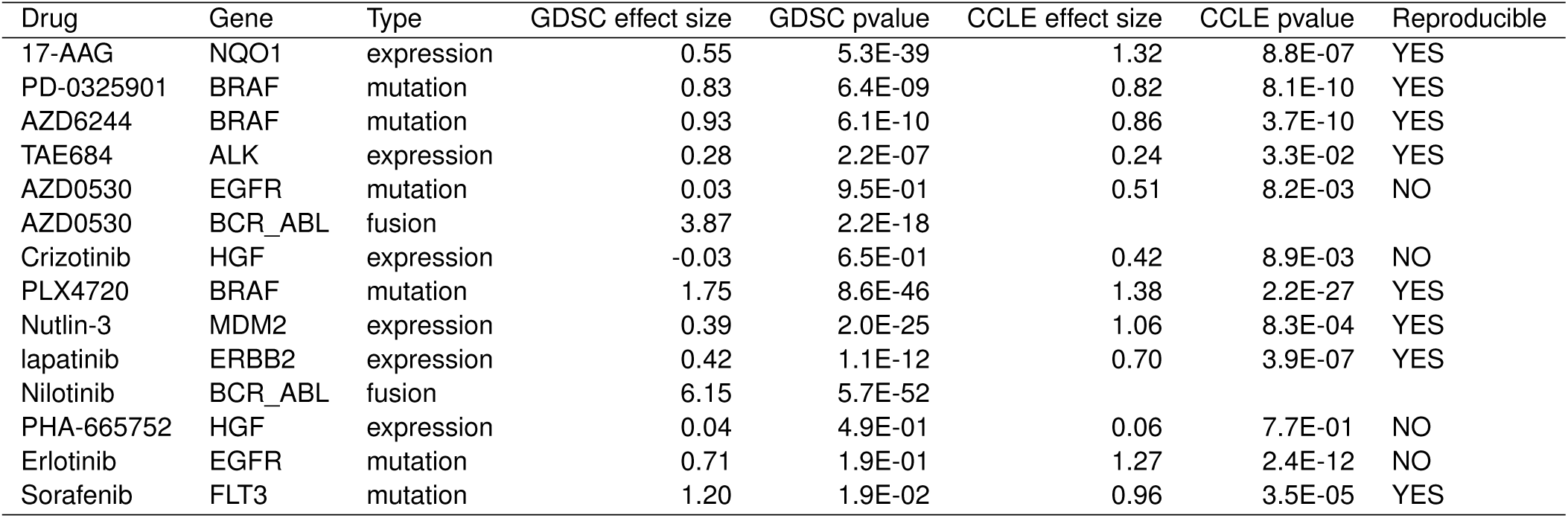
List of known gene-drug associations with their effect size and significance in GDSC and CCLE.

| Drug | Gene | Type | GDSC effect size | GDSC pvalue | CCLE effect size | CCLE pvalue | Reproducible |
| --- | --- | --- | --- | --- | --- | --- | --- |
| 17-AAG | NQO1 | expression | 0.55 | 5.3E-39 | 1.32 | 8.8E-07 | YES |
| PD-0325901 | BRAF | mutation | 0.83 | 6.4E-09 | 0.82 | 8.1E-10 | YES |
| AZD6244 | BRAF | mutation | 0.93 | 6.1E-10 | 0.86 | 3.7E-10 | YES |
| TAE684 | ALK | expression | 0.28 | 2.2E-07 | 0.24 | 3.3E-02 | YES |
| AZD0530 | EGFR | mutation | 0.03 | 9.5E-01 | 0.51 | 8.2E-03 | NO |
| AZD0530 | BCR_ABL | fusion | 3.87 | 2.2E-18 |  |  |  |
| Crizotinib | HGF | expression | -0.03 | 6.5E-01 | 0.42 | 8.9E-03 | NO |
| PLX4720 | BRAF | mutation | 1.75 | 8.6E-46 | 1.38 | 2.2E-27 | YES |
| Nutlin-3 | MDM2 | expression | 0.39 | 2.0E-25 | 1.06 | 8.3E-04 | YES |
| lapatinib | ERBB2 | expression | 0.42 | 1.1E-12 | 0.70 | 3.9E-07 | YES |
| Nilotinib | BCR_ABL | fusion | 6.15 | 5.7E-52 |  |  |  |
| PHA-665752 | HGF | expression | 0.04 | 4.9E-01 | 0.06 | 7.7E-01 | NO |
| Erlotinib | EGFR | mutation | 0.71 | 1.9E-01 | 1.27 | 2.4E-12 | NO |
| Sorafenib | FLT3 | mutation | 1.20 | 1.9E-02 | 0.96 | 3.5E-05 | YES |

**Figure 8:**
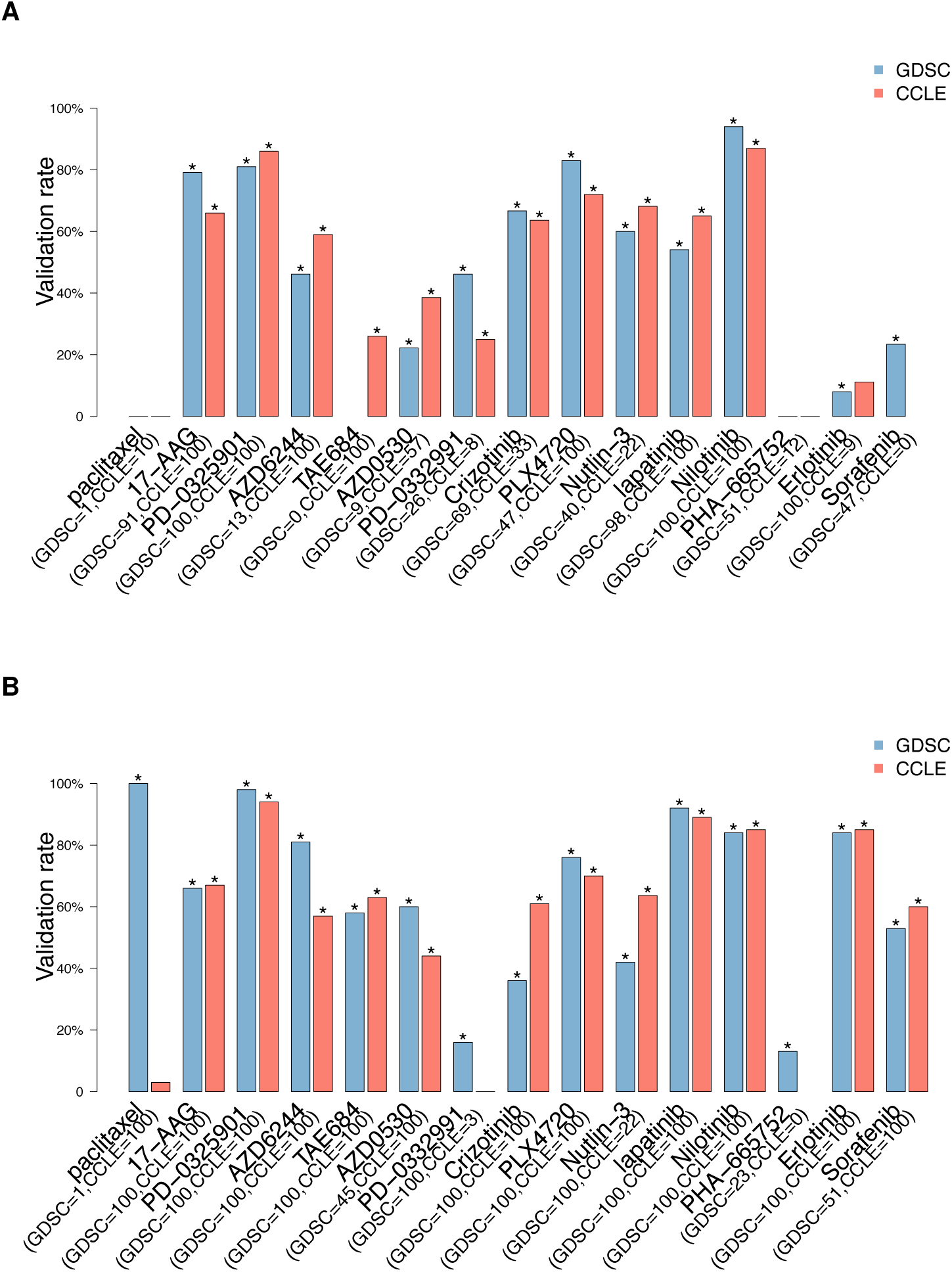
Proportion of gene-drug associations identified in a discovery set (top 100 gene-drug associations as ranked by p-values and FDR < 5%) and validated in an independent validation dataset. In blue and red are the gene-drug associations identified in GDSC and CCLE, respectively. Associations are identified using gene expression data as input and (A) continuous published AUC values as output in a linear model using only common cell lines or (B) all cell lines. The number of selected gene-drugs associations in each datasets is provided in parentheses. The symbol ’*’ represents the significance of the proportion of validated gene-drug associations, computed as the frequency of 1000 random subsets of markers of the same size having equal or greater validation rate compared to the observed rate.

As expected, no consistency was observed for drugs with “no effect” (Figure 6A). For AUC of drugs with narrow and broad effects, Somers’ Dxy was the most stringent, with consistency estimated to be < 0.4 except for two drugs (PD-0325901 and 17-AAG), which were also the two drugs identified as the most consistent using Spearman correlation (`ρ` ∼ 0.6; Figure 6A). However, these statistics did not capture potential consistency for the most highly targeted therapies, nilotinib, crizotinib, and PLX4720, for which the Pearson correlation coefficient gave the best evidence of concordance, as this statistics is strongly influenced by a small number of highly sensitive cell lines (Figure 5).

We then restricted our analysis to the cell lines identified as sensitive in at least one study and computed the same consistency measures (Figure 6B). To our surprise, eliminating the resistant cell lines resulted in decreased consistency for most drugs, which suggests a high level of inconsistency across sensitive cell lines, with the only exceptions of the highly targeted drugs nilotinib and crizotinib.

To test whether discretization of drug sensitivity data into binary calls (“resistant” vs. “sensitive”; see Methods) improves consistency across studies, we used three association statistics, the Matthews correlation coefficient ^16^, Cramer’s V ^17^, and the informedness ^18^ statistics (Figure 6C). These statistics are designed for use with imbalanced classes, which is particularly relevant in large pharmacogenomic datasets where, for targeted therapies, there are often many more resistant cell lines than sensitive ones. As expected, the highly targeted therapies, nilotinib and PLX4720 (and nutlin-3 using informedness), yielded high level of consistency, but this was not the case for the other targeted therapies. We also found that the drug sensitivity calls for drugs with broader inhibitory effects were also poorly correlated between studies (Figure 6C).

We performed the same analysis using IC_50_ values truncated to the maximum concentration used for each drug in each study separately. We observed similar patterns with nilotinib and crizotinib yielding moderate to high consistency across studies (Supplementary Figure 12). Note that Somers’ Dxy rank correlation is biased in the presence of many repeated values in the datasets being analyzed, which is the case for truncated IC_50_ — pairs of cell line with identical IC_50_ values in one dataset but not in the other will not be taken into account as evidence of inconsistency — which explains the artifactual perfect consistency it suggests for both nilotinib and crizotinib.

### Consistency of gene expression across cell lines

Discovering new biomarkers predictive of drug response requires both robust pharmacological data and molecular profiles. In our original study, we showed that the gene expression profiles for each cell line profiled by both GDSC and CCLE were highly consistent. However, we found that mutation profiles were only moderately consistent, a result that was later confirmed by Hudson et al. ^19^.

There have been questions as to whether the measures of consistency we reported for drug response should be compared to those we reported for gene expression. Specifically, we reported correlations based on comparing drug response “across cell lines,” meaning that we examined the correlation of response of each cell line to a particular drug reported by the GDSC with the response of the same cell line to the same drug reported by the CCLE. In contrast we reported correlation of gene expression levels “between cell lines,” meaning that we compared the expression of all genes within each cell line in the GDSC to the expression of all genes in the same cell line in the CCLE (see Supplementary Methods). It has been suggested that a more valid comparison would be to compare both drug response and gene expression across cell lines. We report the results of such an “across cell lines” analysis of gene expression here, computed using techniques analogous to those we used to compare drug response.

We began by comparing the distribution of gene expression measurements generated using the microarray Affymetrix HG-U219 platform in GDSC, the microarray Affymetrix HG-U133PLUS2 platform and the new Illumina RNA-seq data in CCLE (Supplementary Figure 13). We observed similar bimodal distributions, suggesting the presence of a natural cutoff to discriminate between lowly vs. highly expressed genes. We therefore fit a mixture of two gaussians and identified an expression cutoff for each platform separately (Supplementary Figure 13). We then compared the consistency of continuous and discretized gene expression values between (*i*) the microarray Affymetrix HG-U133PLUS2 and Illumina RNA-seq platforms within CCLE (intra-lab consistency); (*ii*) the microarray Affymetrix HG-U219 and HG-U133PLUS2 platforms used in GDSC and CCLE, respectively (microarray, inter-lab consistency); and (*iii*) the microarray Affymetrix HG-U219 and Illumina RNA-seq platforms used in GDSC and CCLE, respectively (inter-lab consistency). Supporting our previous observations, we found that gene expression measurements are significantly more consistent than drug sensitivity values when using all cell lines (Wilcoxon rank sum test p-value < 0.05; Figure 7A; Supplementary Figure 14A).

Similarly to the filtering we performed for drug sensitivity data, we subsequently restricted our analysis to the cell lines showing high expression of a given gene/cell line combination in at least one study. Again, gene expression measurements were significantly more consistent than drug sensitivity values in this case (Wilcoxon rank sum test p-value < 0.05; Figure 7B; Supplementary Figure 14B). When dichotomizing data into lowly/highly expressing cell lines and resistant/sensitive cell lines, the gene expression data were still more consistent (Figure 7C) although the difference was not always significant (Supplementary Figure 14C).

### Consistency of gene-drug associations

The primary goal of the GDSC and CCLE studies was to identify new genomic predictors of drug response for both targeted and cytotoxic therapies. We therefore evaluated whether the good consistency in drug sensitivity data observed for nilotinib, PLX4720 and crizotinib, and the moderate consistency observed for 17-AAG and PD-0332901 would translate in reproducible biomarkers. We estimated gene–drug associations by fitting, for each gene and drug, a linear regression model including microarray-based gene expression as predictor of drug sensitivity, adjusted for tissue source (see Methods). Given the high correlation between the published and recomputed AUC values in each study (Supplementary Figure 5) and their similar consistency (Figure 6), all gene-drug associations were computed using published AUC for clarity.

We first computed the strength and significance of each gene expression in both datasets separately. Similarly to our initial study ^7^, the strength of a given gene-drug association is provided by the standardized coefficient associated to the corresponding gene expression in the linear model and its significance is provided by the p-value of this coefficient (see Methods). We then identified gene-drug associations that were reproducible in both datasets (same sign and FDR < 5%) or that were dataset-specific (different sign or significant in only one dataset) using continuous (Supplementary Figures 15 and 16 for common and all cell lines, respectively) and discretized (Supplementary Figures 17 and 18 for common and all cell lines, respectively) published AUC values as drug sensitivity data. We assessed the overlap of gene-drug associations discovered in both datasets using the Jaccard index ^20^. All jaccard indices were low, with nilotinib yielded the largest overlap of gene-drug associations (32%), followed by PD-0325901 and erlotinib (almost 20%), while the other drugs yielded less than 15% overlap (Supplementary Figure 19). Our results further indicate that larger overlap exists for gene-drug associations identified using the continuous drug sensitivity data compared with associations using discretized drug sensitivity calls (Wilcoxon signed rank test p-value of 4x10^-2^ and 2x10^-3^ for the common set and the full set of cell lines, respectively). We therefore focused our analyses on the gene-drug associations identified using continuous published AUC values. The number (and identity) of gene-drug associations computed using continuous published AUC values are provided in Supplementary Tables 6 and 7 (Supplementary Files 5 and 6) for common and all cell lines, respectively.

Given that simply intersecting significant gene-drug associations identified in each dataset separately yielded poor reproducibility for all drugs, we sought to more closely mimic the biomarker discovery and validation process. We therefore used one dataset to discover significant gene-drug associations and test whether this subset of markers validated in an independent dataset. Using the discovery dataset, gene-drug associations are first ranked by nominal p-values and their FDR is computed. An association is selected if it is part of the top 100 markers and its FDR is less than 5%. This procedure ensure to control for both significance and number of selected biomarkers, which can vary with respect to the cell line panel used for the analysis (larger panels enable the identification of more significant biomarkers due to increased statistical power). A gene-drug association is validated in an independent dataset if its nominal p-value is less than 0.05 and its “direction”, that is whether the marker is associated with sensitivity or resistance, is identical to the one estimated during the discovery process.

We computed the proportions of validated gene-drug associations for each drug using gene expression data in GDSC as discovery set and CCLE as validation set, and vice-versa (Figure 8). Overall, we found that expression-based markers for PD-0325901 and nilotinib yielded a high validation rate (> 80%) with either dataset as discovery set using the common cell lines screened in GDSC and CCLE (Figure 8A). When using the entire cell line panels used in each study, two more drugs - lapatinib and erlotinib - yielded high validation rate (Figure 8B). 17-AAG, and PLX4720 yielded validation rate between 60% and 80%, while the other drugs yielded a validation rate around 50% or lower. For eight out of the fifteen drugs, using the entire panel of cell lines screened in each study (Figure 8B) improved the validation rate compared to limiting the analysis to common cell lines (Figure 8A). However validation rate decreased for five other drugs, suggesting that using large, but different panels of cell lines may increase statistical power but could also introduce biases in the biomarker discovery process.

We then investigated whether higher validation rates would be obtained by using more stringent significance threshold and relaxing the constraint on the number of significant associations in the discovery set (Supplementary Figures 20 and 21). Using common cell lines, we found that proportion of validated gene-drug association monotonically increases with FDR stringency for six drugs, with very high validation rate for the most stringent FDR cutoff (validation rate > 80% for FDR < 0.1%) for 17-AAG, PD-0325901, PLX4720 and nilotinib using either dataset as discovery set (Supplementary Figure 20). Using the entire panel of cell lines in each study actually improved validation rate for six drugs, AZD6244, TAE684, AZD0530, lapatinib — and erlotinib and sorafenib, for which insufficient number of sensitive cell lines were screened in both GDSC and CCLE (Supplementary Figure 21). However, validation rate decreased for 17-AAG, crizotinib and PLX4720, which suggest again that large, but different panels of cell lines might introduce selection bias for some drugs.

### Known biomarkers

As recently reported by Goodspeed et al ^11^, several known biomarkers for targeted therapies have been shown to be predictive in both GDSC and CCLE. In our initial comparative study we also found the following known gene-drug associations:

- BRAF mutations were significantly associated with sensitivity to MEK inhibitors (AZD6244 and PD-0325901) and BRAF^V600E^ inhibitor (PLX4720) with nominal p-values < 0.01; see Supplementary Files 10-13 of our initial study.
- ERBB2 expression was significantly associated with sensitivity to lapatinib with nominal p-value = 0.04 and 8.4□10^-15^ for GDSC and CCLE, respectively; see Supplementary Files 4 and 5 of our initial study.
- NQ01 expression was significantly associated with sensitivity to 17-AAG with nominal p-value = 2.4□10^-13^ and 6.2□10^-14^ for GDSC and CCLE, respectively; see Supplementary Files 4 and 5 of our initial study.
- MDM2 expression was significantly associated with sensitivity to Nutlin-3 with nominal p-value = 7.7□10^-18^ and 7□10^-8^ for GDSC and CCLE, respectively; see Supplementary Files 4 and 5 of our initial study.
- ALK expression was significantly associated with sensitivity to TAE684 with nominal p-value = 1.6□10^-9^ and 1.7□10^-9^ or GDSC and CCLE, respectively; see Supplementary Files 4 and 5 of our initial study.

We revisited our biomarker analysis using the new data released by GDSC and CCLE, and our *PharmacoGx* platform to test whether additional known biomarkers can be identified. In addition to the expression-based gene-drug association reported in Supplementary File 6, we recomputed all gene-drug associations based on mutations (Supplementary File 7) and gene fusions using the entire panel of cell lines in each study. We confirmed the reproducibility of the known associations reported in our initial study, but we were not able to find reproducible associations for EGFR mutations with response to AZD0530 and erlotinib, and HGF expression with response to crizotinib (Table 1). The reproducibility of the vast majority of these previously known associations attest to the relevance of the GDSC and CCLE datasets although our results demonstrated that the noise and inconsistency in drug sensitivity data renders discovery of new biomarkers difficult for the majority of the drugs.

## DISCUSSION

Our original motivation in analyzing the GDSC and CCLE data was to develop predictive gene expression biomarkers of drug response. When we applied a number of methods using one study to select gene expression features and to train a classifier, and then tested it by trying to predict reported drug response in the second study, our predictive models failed to validate for half of the drugs tested ^4^. Indeed, out of nine predictors yielding concordance index ^21^ ≥0.65 in cross-validation in the training set (GDSC), only four were validated in identical cell lines treated with the same drugs in the validation set (CCLE) ^4^.

As we explored the reasons for this failure, we first checked whether cell lines could have drifted and consequently exhibited different transcriptional profiles between GDSC and CCLE. We found that any genome-wide expression profile in one study would almost always identify “itself” (its purported biological replica) as being most similar among the cell lines in the other study. In a way this is not surprising. When gene expression studies were in their infancy, there were many reports that compared the results from studies and found that they were inconsistent and unreproducible in new studies — as demonstrated by the countless biomarkers that fail to reproduce beyond their initial publication. As a result, scientists involved in gene expression studies “circled the wagons” and developed both much more standardized laboratory protocols and “best practices” for reproducible analysis, including data normalization and batch corrections, that now mean that independent measurements from different laboratories are far more often consistent and so can be used for signature development and validation ^22,23^.

Unexpectedly, when we compared phenotypic measures of drug response that were released by the GDSC and CCLE projects, we found discrepancies in growth inhibition effects of multiple anticancer agents. What that means in practice is that, for some drugs, an expression-based biomarker of drug response learned from one study would not likely be predictive of the reported response in the other. And consequently neither of the studies might be useful in predicting response in patients as many had hoped when these large pharmacogenomic screens were published.

The feedback from the scientific community on our analysis, the availability of new data from the GDSC and CCLE, as well as improvements in the *PharmacoGx* software platform we developed to support this type of analyses ^9^, prompted us to revisit the question of consistency in these studies to see if we could find a principled way to identify correlated drug response phenotypes. By testing a variety of methods of classifying the data, and choosing the metric which gave the best consistency for each drug, we were able to find moderate to good consistency of sensitivity data for two broad effect and three highly targeted drugs. We also confirmed the overall lack of consistency between the studies for eight drugs, while there were not enough sensitive cell lines that had been screened by both GDSC and CCLE to properly assess consistency for the remaining three drugs. The summary box included with this paper briefly describes the most significant issues that people have raised in discussing our previous findings with us and summarizes what we have found in our reanalysis.

Some have suggested that one way to improve correlation would have have been to compare the studies and throw out the most discordant data as noise and then compare the remaining concordant data. While this would certainly find concordance in the remaining data, the approach is equivalent to fitting data to a desired result, which is bad practice and certainly could not be extended to other data sets or to the classification of patient tumors as responsive or nonresponsive to a particular therapy.

There is, however, merit in the suggestion that one would not expect to see correlation in noise. And noise is precisely what one would expect to see in drug response data from cell lines that are resistant to a particular drug or nonresponsive across the range of doses tested. As reported here, filtering the data in each study independently to classify cell lines in a binary fashion, and then comparing the binary classification between studies using a variety of metrics developed to handle the intricacies of this sort of response data, also failed to find simple correlations in the data, except for three of the highly targeted therapies, nilotinib, PLX4720 and crizotinib. What this ultimately means is that the most and the least sensitive cell lines would not appear to be the same when comparing the two studies.

There are many reasons for potential differences in measured phenotypes reported by the GDSC and CCLE, including substantial differences in doses used for each drug and in the methods used to both assay cell viability and to estimate drug response parameters. Ultimately what our analysis suggests is that not only is there a need to carefully and appropriately compare measurements, but also that there is a pressing need for standardization of both laboratory and computational methods for assaying drug response.

The primary goal of the GDSC and CCLE studies was to link molecular features of a large panel of cancer cell lines to their sensitivity to cytotoxic and targeted drugs. The reproducibility of most of the known gene-drug associations provides evidence that these large pharmacogenomic datasets are biologically relevant. When we investigated whether we could find significant gene-drug associations discovered in one dataset that validate in the other independent dataset, we observed over 75% validation rate for the most significant expression-based biomarkers for eight of fifteen drugs, which is a major improvement over our initial comparative study. However, this does not suggest that one can use these studies to find new, reproducible gene-drug associations for the rest of the drugs - excluding paclitaxel and PHA-655752 for which no significant biomarkers could be identified - as the majority of associations can be found in only one dataset but not in both.

This study has several potential limitations. First, while the raw drug sensitivity data are publicly available for GDSC, these data have not been released within the CCLE study. We could not fit the drug dose-response curves using the technical triplicates but rather relied on the published median sensitivity values. Second, we discretized drug sensitivity values by selecting a common threshold to discriminate between highly resistant (AUC ≤ 0.2 and IC_50_ ≥ 1 μM) and the rest of the cell lines for all the targeted agents. However, it is clear that such a threshold could be optimized for each drug, which might have an impact on the consistency of drug phenotypes and gene-drug associations based on binary sensitivity calls (note that the same applies for gene expression data as well). Unfortunately the size of the current drug sensitivity datasets is not sufficient to develop drug-specific thresholds for sensitivity values but the release of larger pharmacogenomic studies may allow us to address this issue in the near future. Lastly, the current set of mutations of mutations assessed in both study is small (64 mutations), which drastically limits the search for mutation-based and other genomic aberrations associated with drug response. Again, potential releases of whole-genome and whole-exome sequencing will enable to better explore the genomic space of biomarkers in cancer cell lines, and their reproducibility across studies.

## CONCLUSION

As is true of many scientists working in genomics and oncology, we were excited when the GDSC and CCLE released their initial data sets and were very hopeful that these projects would help to accelerate drug discovery and further the development of precision medicine in oncology. However, what we found initially, and what the reanalysis presented here further indicates, is that there are inconsistencies between the measured phenotypic response to drugs in these studies. Even in our reanalysis, where we used methods specific to individual drugs and the response characteristics of the cell lines tested, we were only able to find new biomarkers predictive of response for around half of the drugs screened in both studies. Consequently, it is challenging to use the data from these studies to develop general purpose classification rules for all drugs.

Our finding that gene expression measurements are significantly more consistent than drug sensitivity data, indicate that the main barrier to biomarker development using these data is the unreliability in the reported response phenotypes for many drugs. For studies such as these to realize their full potential, additional work must be done to develop robust and reproducible experimental and analytical protocols so that the same compound, tested on the same set of cell lines by different groups, yields consistent and comparable results. Barring this, a predictive biomarker of response developed from one study is unlikely to be able to reliably validated on another, and consequently, is unlikely to be useful in predicting patient response.

From having worked in large-scale genomic analyses, we recognize the challenges involved in planning and executing such studies and commend the GDSC and CCLE for their work and for making all the data available. However, we strongly encourage the GDSC, the CCLE, the pharmacogenomics and bioinformatics communities as a whole, to invest the necessary time and effort to standardize drug response assays in order to achieve greater consistency and to assure that measurements in cell lines are relevant for predicting response in patients. Ultimately, that effort will help to assure that mammoth undertakings in drug characterization can deliver on their promise to identify better therapies and biomarkers predictive of response.

## METHODS

Code and software are available upon request and will be made publicly available upon publication.

### The PharmacoGx platform

The lack of standardization of cell line and drug identifiers hinders comparison of molecular and pharmacological data between large-scale pharmacogenomic studies, such as the GDSC and CCLE. To address this issue we developed *PharmacoGx*, a computational platform enabling users to download and interrogate large pharmacogenomic datasets that were extensively curated to ensure maximum overlap and consistency ^9^. *PharmacoGx* provides (*i*) a new object class, called *PharmacoSet*, that acts as a container for the high-throughput pharmacological and molecular data generated in large pharmacogenomics studies (detailed structure provided in Supplementary Methods); and (*ii*) a set of parallelized functions to assess the reproducibility of pharmacological and molecular data and to identify molecular features associated with drug effects. The *PharmacoGx* package is open-source and publicly available on the Comprehensive R Archive Network (https://cran.rproject.org/web/packages/PharmacoGx/).

### The GDSC (formerly CGP) dataset

#### Drug sensitivity data

We used the data release 5 (June 2014) with 6,734 new IC_50_ values for a total of 79,903 drug dose-response curves for 139 different drugs tested on a panel of up to 672 unique cell lines.

#### Molecular profiles

Gene expression data were downloaded from ArrayExpress, accession number E-MTAB-3610. This new data were generated using Affymetrix HG-U219 microarray platform. We processed and normalized the CEL files using RMA ^24^ with BrainArray ^25^ chip description file based on Ensembl gene identifiers (version 19). This resulted in a matrix of normalized expression for 17,616 unique Ensembl gene ids.

Mutation and gene fusion calls were downloaded from the GDSC website (http://www.cancerrxgene.org/downloads/) and processed as in our initial study ^7^.

### The CCLE dataset

#### Drug sensitivity data

We used the drug sensitivity data available from the CCLE website (http://www.broadinstitute.org/ccle/home) and updated on February 2015 with a total number of 11670 dose-response curves for 24 drugs tested in a panel of up to 504 cell lines.

#### Molecular profiles

Gene expression data were downloaded from the CCLE website and CGHub ^26^ for the Affymetrix HG-U133PLUS2 and Illumina HiSeq 2500 platforms, respectively. Normalization of microarray data (1036 cell lines) was performed the same way than for GDSC. RNA-seq data (935 cell lines) were downloaded as BAM files previously aligned using TopHat ^27^ and the quantification of gene expression was performed using Cufflinks ^27^ based on Ensembl GrCh37 human reference genome.

Mutation data were retrieved from the CCLE website and processed as in our initial study ^7^.

#### Curation of drug and cell line identifiers

The lack of standardization for cell line names and drug identifiers represents a major barrier for performing comparative analyses of large pharmacogenomics studies, such as GDSC and CCLE. We therefore curated these datasets to maximize the overlap in cell lines and drugs by assigning a unique identifier to each cell line and drug. Entities with the same unique identifier were matched. Manual search was then applied to match any remaining cell lines or drugs which were not matched based on string similarity. The cell line curation was validated by ensuring that the cell lines with matched name had a similar SNP fingerprint (see below). The drug curation was validated by examining the extended fingerprint of each of their SMILES strings ^28^ and ensuring that the Tanimoto similarity ^29^ between two drugs called as the same, as determined by this fingerprint, was above 0.95.

#### Cell line identity using SNP fingerprinting

To assess the identity of cell lines from GDSC and CCLE, data of low quality was first excluded from our analysis panel (detailed procedure described in Supplementary Methods). Of the 973 CEL files from GDSC, only 66 fell below the 0.4 threshold (6.88%) for contrast QC scores, indicating issues in resolving base calls. Additionally, five of the 1,190 CEL files from CCLE had an absolute difference between contrast QC scores for Nsp and Sty fragments greater than 2, thus indicating some issues with the efficacy of one enzyme set during sample preparation. CEL files with contrast QC scores indicative of some sort of issue with the assay that would affect the genotype call rate or birdseed accuracy were removed and genotype calling was conducted on the remaining CEL files using Birdseed version 2. The resulting files were then filtered to keep only the 1006 SNP fingerprints that originated from CEL files that had a common cell line annotation between GDSC and CCLE (503 CEL files from each). Finally, pairwise concordances of all SNP fingerprints were generated according to the method outlined by Hong et al. ^30^.

#### Drug dose-response curves

To identify artefactual drug dose-response curves due to experimental or normalization issues, we developed simple quality controls (QC; details in Supplementary Methods). Briefly, we checked whether normalized viability measurements range between 0% and 100% and that consecutive measurements remain stable or decrease monotonically reflecting response to the drug being tested. The drug dose-response curves which did not pass these simple QC were flagged and removed from subsequent analyses as the curve fitting would have yielded erroneous results.

All dose-response curves were fitted to the equation

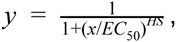

where *y* = 0 denotes death of all infected cells, *y* = *y*(0) = 1 denotes no effect of the drug dose, *EC*_50_ is the concentration at which viability is reduced to half of the viability observed in the presence of an arbitrarily large concentration of drug, and *HS* is a parameter describing the cooperativity of binding. *HS* < 1 denotes negative binding cooperativity, *HS* = 1 denotes noncooperative binding, and *HS* > 1 denotes positive binding cooperativity. The parameters of the curves were fitted using the least squares optimization framework. Comparison of our dose-response curve model with those used in the GDSC and CCLE publications is provided in Supplementary Methods.

### Discretization of pharmacogenomic data

#### Drug sensitivity data

To discretize the drug sensitivity data, we used AUC ≤ 0.2 (IC_50_ ≥ 1 μM) and AUC ≤ 0.4 (IC_50_ ≥ 10 μM) to identify the “resistant” cell lines for targeted and cytotoxic drugs, respectively, while the rest of the cell lines are classified as “sensitive. These reasonable, although somewhat arbitrary, cutoffs enabled to explore the potential of such binary drug sensitivity calls as new drug phenotypic measures to find consistency in drug sensitivity data and gene-drug associations.

#### Gene expression data

To discretize the drug sensitivity data into lowly vs. highly expressed genes, we fit a mixture of 2 gaussians of unequal variance using the full distribution of expression values of the 17,401 genes in common between GDSC and CCLE datasets. We defined the expression threshold as the expression value for which the posterior probability of belonging to the left tail of the highly expression distribution is 10%.

#### Mutation data

Similarly to the GDSC and CCLE publications, we transformed the original mutation data into binary values that represent the absence (0) or presence (1) of any missense mutations in a given gene in a given cell line.

#### Gene-drug associations

We assessed the association, across cell lines, between a molecular feature and response to a given drug, referred to as gene-drug association, using a linear regression model adjusted for tissue source:

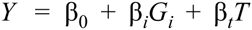

where *Y* denotes the drug sensitivity variable, *G*_*i*_ and *T* denote the expression of gene *i* and the tissue source respectively, and *β*s are the regression coefficients. The strength of gene-drug association is quantified by *β*_*i*_, above and beyond the relationship between drug sensitivity and tissue source. The variables *Y* and *G* are scaled (standard deviation equals to 1) to estimate standardized coefficients from the linear model. Significance of the gene-drug association is estimated by the statistical significance of *β*_*i*_ (two-sided t test). When applicable, p-values were corrected for multiple testing using the false discovery rate (FDR) approach ^31^.

As we recognized that continuous drug sensitivity is not normally distributed, which violates one of the assumption of the linear regression model described above, we also assessed the consistency of gene-drug association using discretized (binary) drug sensitivity calls as the response variable in a logistic regression model adjusted for tissue source, similarly to the linear regression model.

### Measure of consistency

#### Area between curves (ABC)

To quantify the difference between two dose-response curves, we computed the area between curves (ABC). ABC is calculated by taking the unsigned area between the two curves over the intersection of the concentration range tested in the two experiments of interest, and normalizing that area by the length of the intersection interval. In the present study, we compared the curves fitted for the same drug-cell line combinations tested both in GDSC and CCLE. Further details are provided in Supplementary Methods.

#### Pearson correlation coefficient (PCC)

PCC is a measure of the linear correlation between two variables, giving a value between +1 and −1 inclusive, where 1 represents total positive correlation, 0 represents no correlation, and −1 represents total negative correlation ^13^. PCC is sensitive to the presence of outliers, like a few sensitive cell lines in the case of drug sensitivity data measured for highly targeted therapies or genes rarely expressed.

#### Spearman rank correlation coefficient (SCC)

SCC is a nonparametric measure of statistical dependence between two variables and is is defined as the Pearson correlation coefficient between the ranked variables ^14^. It assesses how well the relationship between two variables can be described using a monotonic function. If there are no repeated data values, a perfect Spearman correlation of +1 or −1 occurs when each of the variables is a perfect monotone function of the other. Contrary to PCC, SCC can capture non linear relationship between variables but is insensitive to outliers, which is frequent for drug sensitivity data measured for highly targeted therapies or genes rarely expressed.

#### Somers’ Dxy rank correlation (DXY)

DXY is a non-parametric measure of association equivalent to (*C* - 0.5) * 2 where *C* represents the concordance index ^21^ that is the probability that two variables will rank a random pair of samples the same way ^15^.

#### Matthews correlation coefficient (MCC)

MCC is used in machine learning as a measure of the quality of binary classifications ^16^. It takes into account true and false positives and negatives and is generally regarded as a balanced measure which can be used even if the classes are of very different sizes. MCC is in essence a correlation coefficient between two binary classifications; it returns a value between −1 (perfect opposite classification) and +1 (identical classifications), with 0 representing association no better than random chance.

#### Cramer’s V (CRAMERV)

CRAMERV is a measure of association between two nominal variables, based on Pearson’s chi-squared statistic, giving a value between 0 (no association) and +1 (perfect association) ^17^. In the case of 2×2 contingency table, such as binary drug sensitivity or gene expression measurements, CRAMERV is equivalent to the Phi coefficient.

#### Informedness (INFORM)

For a 2x2 contingency table comparing two binary classifications, INFORM can be defined as Specificity + Sensitivity -1, which is equivalent to true positive rate - false positive rate ^18^. The magnitude of INFORM gives the probability of an informed decision between the two classes, where INFORM > 0 represents appropriate use of information, 0 represents chance-level decision, < 0 represents perverse use of information.

## FIGURES/TABLES SUPPLEMENTARY INFORMATION

Revisiting inconsistency in large pharmacogenomic studies

## 3 Supplementary Tables

**Supplementary Table 1:** Consistency of AUC values as published. Values are missing when less than 3 observations were available in a given category (resistant or sensitive). PCC: Pearson Correlation Coefficient; SCC: Spearman correlation coefficient; DXY: Somer’s Dxy index; MCC: Mathew correlation coefficient; CRAMERV: Cramer’s V measure of association; INFORM: Informedness measure of association. FULL: Consistency computed using all the common drug-cell line combinations; SENS: Consistency computed using the drug-cell line combinations measured as sensitive in at least one study.

|  | PCC.FULL | SCC.FULL | DXY.FULL | PCC.SENS | SCC.SENS | DXY.SENS | MCC | CRAMERV | INFORM |
| --- | --- | --- | --- | --- | --- | --- | --- | --- | --- |
| paclitaxel | 0.39 | 0.38 | 0.26 | 0.42 | 0.39 | 0.27 | 0.16 | 0.16 | 0.14 |
| 17-AAG | 0.55 | 0.56 | 0.39 | 0.50 | 0.52 | 0.36 | 0.15 | 0.15 | 0.33 |
| PD-0325901 | 0.67 | 0.63 | 0.44 | 0.47 | 0.45 | 0.31 | 0.43 | 0.43 | 0.39 |
| AZD6244 | 0.43 | 0.30 | 0.19 | 0.15 | 0.10 | 0.06 | 0.45 | 0.45 | 0.30 |
| TAE684 | 0.46 | 0.41 | 0.28 | 0.02 | -0.26 | -0.21 | 0.28 | 0.28 | 0.27 |
| AZD0530 | 0.51 | 0.43 | 0.27 | 0.08 | 0.27 | 0.23 | 0.39 | 0.39 | 0.24 |
| PD-0332991 | 0.36 | 0.35 | 0.12 | -0.06 | -0.16 | -0.13 | 0.09 | 0.09 | 0.18 |
| Crizotinib | 0.58 | 0.16 | 0.04 | 0.91 | 0.60 | 0.40 | 0.44 | 0.44 | 0.20 |
| PLX4720 | 0.64 | 0.37 | 0.13 | -0.01 | 0.02 | -0.02 | 0.69 | 0.69 | 0.71 |
| Nutlin-3 | 0.38 | 0.32 | 0.15 | 0.02 | -0.02 | 0.00 | 0.42 | 0.42 | 0.95 |
| lapatinib | 0.59 | 0.43 | 0.17 | 0.17 | 0.14 | 0.07 | 0.49 | 0.49 | 0.39 |
| Nilotinib | 0.78 | 0.16 | 0.01 | 0.63 | 0.60 | 0.40 | 0.77 | 0.77 | 0.60 |
| PHA-665752 | 0.16 | -0.02 | -0.10 | -0.50 | -0.20 | -0.20 | -0.02 | 0.02 | -0.01 |
| Erlotinib | 0.34 | 0.46 | 0.16 |  | -1.00 |  | -0.01 | 0.01 | -0.01 |
| Sorafenib | 0.30 | 0.28 | -0.00 | -0.91 | -0.50 |  | -0.02 | 0.02 | -0.02 |

**Supplementary Table 2:**
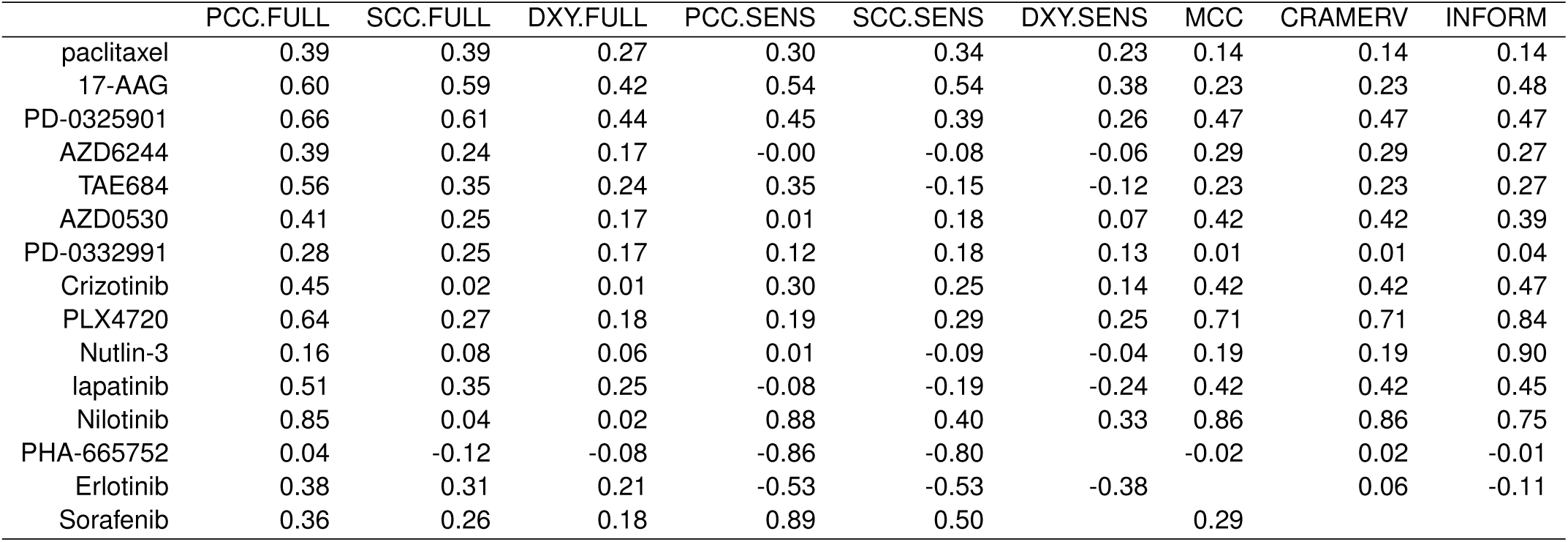
Consistency of AUC values as recomputed within *PharmacoGx*. Values are missing when less than 3 observations were available in a given category (resistant or sensitive). PCC: Pearson Correlation Coefficient; SCC: Spearman correlation coefficient; DXY: Somer’s Dxy index; MCC: Mathew correlation coefficietn; CRAMERV: Cramer’s V measure of association; INFORM: Informedness measure of association. FULL: Consistency computed using all the common drug-cell line combinations; SENS: Consistency computed using the drug-cell line combinations measured as sensitive in at least one study.

|  | PCC.FULL | SCC.FULL | DXY.FULL | PCC.SENS | SCC.SENS | DXY.SENS | MCC | CRAMERV | INFORM |
| --- | --- | --- | --- | --- | --- | --- | --- | --- | --- |
| paclitaxel | 0.39 | 0.39 | 0.27 | 0.30 | 0.34 | 0.23 | 0.14 | 0.14 | 0.14 |
| 17-AAG | 0.60 | 0.59 | 0.42 | 0.54 | 0.54 | 0.38 | 0.23 | 0.23 | 0.48 |
| PD-0325901 | 0.66 | 0.61 | 0.44 | 0.45 | 0.39 | 0.26 | 0.47 | 0.47 | 0.47 |
| AZD6244 | 0.39 | 0.24 | 0.17 | -0.00 | -0.08 | -0.06 | 0.29 | 0.29 | 0.27 |
| TAE684 | 0.56 | 0.35 | 0.24 | 0.35 | -0.15 | -0.12 | 0.23 | 0.23 | 0.27 |
| AZD0530 | 0.41 | 0.25 | 0.17 | 0.01 | 0.18 | 0.07 | 0.42 | 0.42 | 0.39 |
| PD-0332991 | 0.28 | 0.25 | 0.17 | 0.12 | 0.18 | 0.13 | 0.01 | 0.01 | 0.04 |
| Crizotinib | 0.45 | 0.02 | 0.01 | 0.30 | 0.25 | 0.14 | 0.42 | 0.42 | 0.47 |
| PLX4720 | 0.64 | 0.27 | 0.18 | 0.19 | 0.29 | 0.25 | 0.71 | 0.71 | 0.84 |
| Nutlin-3 | 0.16 | 0.08 | 0.06 | 0.01 | -0.09 | -0.04 | 0.19 | 0.19 | 0.90 |
| lapatinib | 0.51 | 0.35 | 0.25 | -0.08 | -0.19 | -0.24 | 0.42 | 0.42 | 0.45 |
| Nilotinib | 0.85 | 0.04 | 0.02 | 0.88 | 0.40 | 0.33 | 0.86 | 0.86 | 0.75 |
| PHA-665752 | 0.04 | -0.12 | -0.08 | -0.86 | -0.80 |  | -0.02 | 0.02 | -0.01 |
| Erlotinib | 0.38 | 0.31 | 0.21 | -0.53 | -0.53 | -0.38 |  | 0.06 | -0.11 |
| Sorafenib | 0.36 | 0.26 | 0.18 | 0.89 | 0.50 |  | 0.29 |  |  |

**Supplementary Table 3:** Consistency of AUC* (STAR) values as recomputed within *PharmacoGx*. Values are missing when less than 3 observation where available in a given category (resistant or sensitive). PCC: Pearson Correlation Coefficient; SCC: Spearman correlation coefficient; DXY: Somer’s Dxy index; MCC: Mathew correlation coefficient; CRAMERV: Cramer’s V measure of association; INFORM: Informedness measure of association. FULL: Consistency computed using all the common drug-cell line combinations; SENS: Consistency computed using the drug-cell line combinations measured as sensitive in at least one study.

|  | PCC.FULL | SCC.FULL | DXY.FULL | PCC.SENS | SCC.SENS | DXY.SENS | MCC | CRAMERV | INFORM |
| --- | --- | --- | --- | --- | --- | --- | --- | --- | --- |
| paclitaxel | 0.43 | 0.43 | 0.30 | 0.26 | 0.35 | 0.23 | 0.26 | 0.26 | 0.32 |
| 17-AAG | 0.54 | 0.52 | 0.36 | 0.38 | 0.35 | 0.24 | 0.36 | 0.36 | 0.46 |
| PD-0325901 | 0.68 | 0.61 | 0.44 | 0.48 | 0.40 | 0.26 | 0.41 | 0.41 | 0.41 |
| AZD6244 | 0.37 | 0.24 | 0.16 | -0.04 | -0.13 | -0.09 | 0.27 | 0.27 | 0.26 |
| TAE684 | 0.48 | 0.31 | 0.22 | 0.27 | -0.15 | -0.14 | 0.17 | 0.17 | 0.20 |
| AZD0530 | 0.43 | 0.21 | 0.15 | 0.00 | 0.07 | 0.00 | 0.32 | 0.32 | 0.35 |
| PD-0332991 | 0.15 | 0.12 | 0.07 | -0.04 | 0.17 | 0.12 | -0.04 | 0.04 | -0.10 |
| Crizotinib | 0.41 | 0.14 | 0.09 | 0.13 | -0.12 | -0.07 | 0.44 | 0.44 | 0.35 |
| PLX4720 | 0.59 | 0.19 | 0.13 | 0.24 | 0.23 | 0.16 | 0.59 | 0.59 | 0.74 |
| Nutlin-3 | 0.10 | 0.06 | 0.04 | 0.07 | -0.10 | -0.08 | 0.19 | 0.19 | 0.90 |
| lapatinib | 0.52 | 0.36 | 0.24 | -0.05 | -0.13 | -0.17 | 0.17 | 0.17 | 0.26 |
| Nilotinib | 0.84 | 0.00 | -0.01 | -0.92 | -0.50 |  | 1.00 | 1.00 | 1.00 |
| PHA-665752 | 0.02 | -0.13 | -0.08 | -0.99 | -0.50 |  | -0.02 | 0.02 | -0.01 |
| Erlotinib | 0.29 | 0.29 | 0.19 | -0.61 | -0.31 | -0.24 |  | 0.06 | -0.11 |
| Sorafenib | 0.31 | 0.11 | 0.07 | 1.00 | 1.00 |  | 0.57 | 0.57 | 0.98 |

**Supplementary Table 4:** Consistency of IC_50_ values as published. Values are missing when less than 3 observations were available in a given category (resistant or sensitive). PCC: Pearson Correlation coefficient; SCC: Spearman correlation coefficient; DXY: Somer’s Dxy index; MCC: Mathew correlation coefficient; CRAMERV: Cramer’s V measure of association; INFORM: Informedness measure of association. FULL: Consistency computed using all the common drug-cell line combinations; SENS: Consistency computed using the drug-cell line combinations measured as sensitive in at least one study.

|  | PCC.FULL | SCC.FULL | DXY.FULL | PCC.SENS | SCC.SENS | DXY.SENS | MCC | CRAMERV | INFORM |
| --- | --- | --- | --- | --- | --- | --- | --- | --- | --- |
| paclitaxel | 0.01 | 0.10 | 0.03 | -0.15 | -0.03 | -0.06 | 0.00 | 0.00 | 0.00 |
| 17-AAG | 0.50 | 0.58 | 0.44 | 0.42 | 0.50 | 0.36 | 0.36 | 0.36 | 0.51 |
| PD-0325901 | 0.62 | 0.58 | 0.14 | 0.41 | 0.37 | 0.14 | 0.45 | 0.45 | 0.50 |
| AZD6244 | 0.47 | 0.38 | 0.10 | -0.18 | -0.12 | -0.12 | 0.37 | 0.37 | 0.28 |
| TAE684 | 0.47 | 0.29 | 0.22 | 0.22 | -0.12 | -0.10 | 0.29 | 0.29 | 0.34 |
| AZD0530 | 0.52 | 0.34 | 0.05 | -0.30 | -0.09 | -0.07 | 0.26 | 0.26 | 0.30 |
| PD-0332991 | 0.36 | 0.11 | -0.61 | 0.04 | 0.22 | -0.17 | 0.21 | 0.21 | 0.42 |
| Crizotinib | 0.65 | 0.37 | 0.83 | 0.23 | 0.40 | 0.33 | 0.66 | 0.66 | 0.66 |
| PLX4720 | 0.64 | 0.49 | 0.26 | -0.16 | -0.12 | -0.08 | 0.46 | 0.46 | 0.39 |
| Nutlin-3 | 0.41 | 0.38 | -0.48 |  | -1.00 |  | -0.00 | 0.00 | -0.00 |
| lapatinib | 0.62 | 0.29 | 0.44 | 0.34 | 0.56 | 0.33 | 0.56 | 0.56 | 0.49 |
| Nilotinib | 0.76 | 0.30 | 0.98 | 0.62 | 0.40 | 0.45 | 0.70 | 0.70 | 0.50 |
| PHA-665752 | -0.03 | -0.04 |  |  | -1.00 |  | -0.01 | 0.01 | -0.01 |
| Erlotinib | -0.05 | 0.04 | -0.46 | -1.00 | -0.87 |  | -0.02 | 0.02 | -0.01 |
| Sorafenib | 0.14 | 0.24 | 0.13 | 0.07 | 0.26 |  |  |  | 0.00 |

**Supplementary Table 5:**
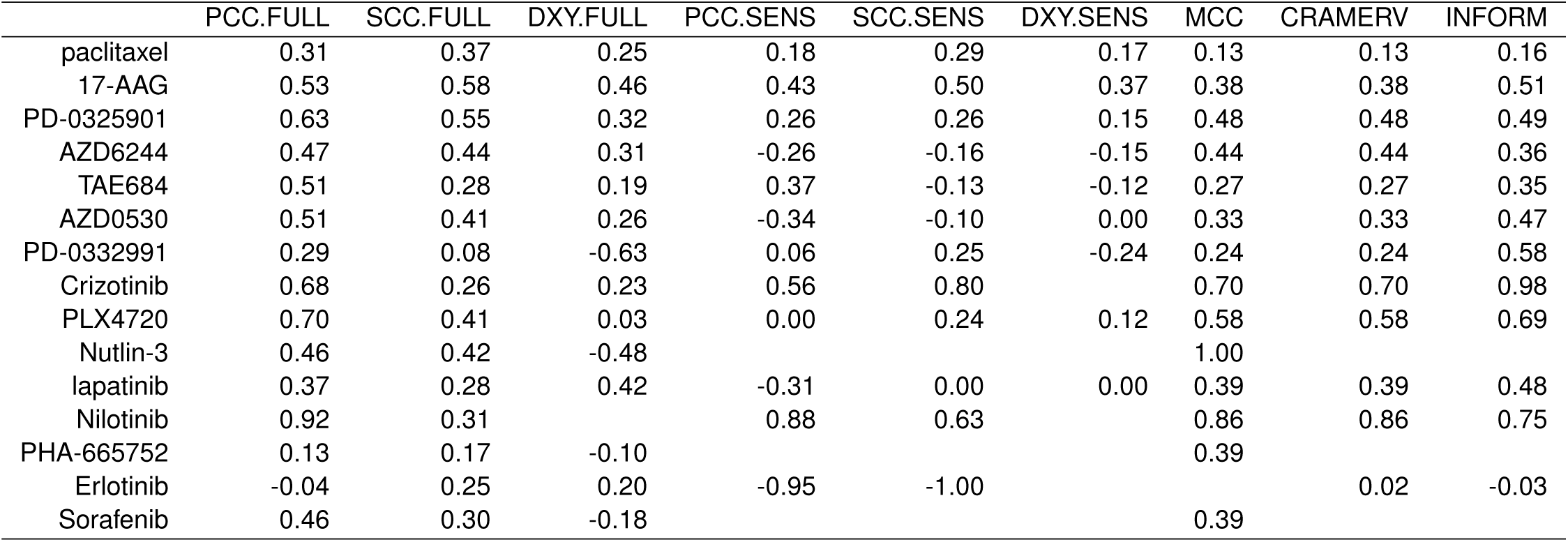
Consistency of IC_50_ values as recomputed within *PharmacoGx*. Values are missing when less than 3 observations were available in a given category (resistant or sensitive). PCC: Pearson Correlation Coefficient; SCC: Spearman correlation coefficient; DXY: Somer’s Dxy index; MCC: Mathew correlation coefficient; CRAMERV: Cramer’s V measure of association; INFORM: Informedness measure of association. FULL: Consistency computed using all the common drug-cell line combinations; SENS: Consistency computed using the drug-cell line combinations measured as sensitive in at least one study.

**Supplementary Table 6:** Table reporting the total number of expression-based gene-drug associations identified using continuous published AUC and only the cell lines in common between GDSC and CCLE. The proportion of associations that are dataset-specific or reproducible across GDSC and CCLE are provided in the last three columns. The column ’% Both’ reports the overlap of gene-drug associations between the two studies, as computed using the Jaccard index.

|  | # GDSC | # CCLE | % GDSC | % CCLE | % Both |
| --- | --- | --- | --- | --- | --- |
| paclitaxel | 1 | 10 | 9 | 91 | 0 |
| 17-AAG | 62 | 221 | 20 | 71 | 9 |
| PD-0325901 | 481 | 881 | 29 | 53 | 18 |
| AZD6244 | 11 | 409 | 3 | 97 | 0 |
| TAE684 | 0 | 150 | 0 | 100 | 0 |
| AZD0530 | 8 | 56 | 12 | 86 | 2 |
| PD-0332991 | 26 | 8 | 76 | 24 | 0 |
| Crizotinib | 59 | 23 | 64 | 25 | 11 |
| PLX4720 | 24 | 174 | 11 | 79 | 10 |
| Nutlin-3 | 33 | 15 | 60 | 27 | 13 |
| lapatinib | 76 | 128 | 34 | 57 | 10 |
| Nilotinib | 289 | 311 | 33 | 36 | 32 |
| PHA-665752 | 51 | 12 | 81 | 19 | 0 |
| Erlotinib | 171 | 9 | 95 | 5 | 0 |
| Sorafenib | 47 | 0 | 100 | 0 | 0 |

**Supplementary Table 7:**
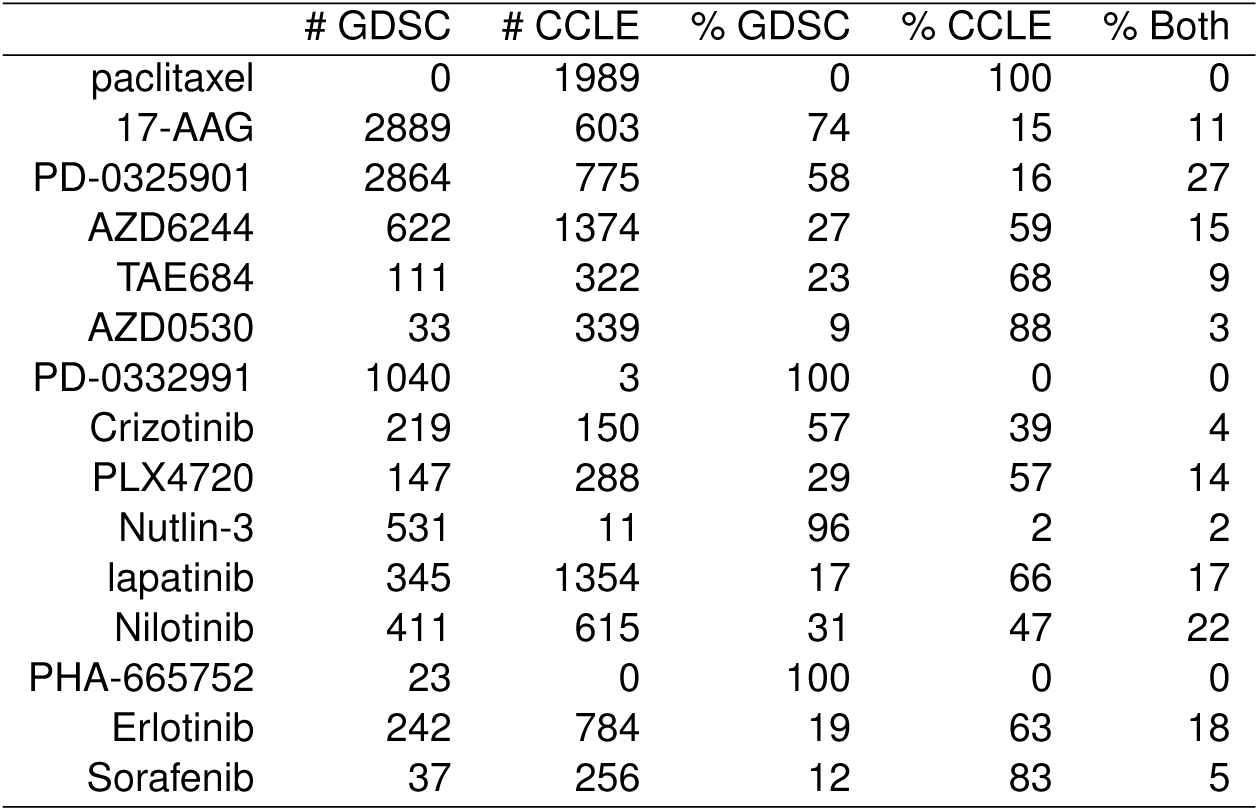
Table reporting the total number of expression-based gene-drug associations identified using continuous published AUC and all cell lines in GDSC and CCLE. The proportion of associations that are dataset-specific or reproducible across GDSC and CCLE are provided in the last three columns. The column ’% Both’ reports the overlap of gene-drug associations between the two studies, as computed using the Jaccard index

|  | # GDSC | # CCLE | % GDSC | % CCLE | % Both |
| --- | --- | --- | --- | --- | --- |
| paclitaxel | 0 | 1989 | 0 | 100 | 0 |
| 17-AAG | 2889 | 603 | 74 | 15 | 11 |
| PD-0325901 | 2864 | 775 | 58 | 16 | 27 |
| AZD6244 | 622 | 1374 | 27 | 59 | 15 |
| TAE684 | 111 | 322 | 23 | 68 | 9 |
| AZD0530 | 33 | 339 | 9 | 88 | 3 |
| PD-0332991 | 1040 | 3 | 100 | 0 | 0 |
| Crizotinib | 219 | 150 | 57 | 39 | 4 |
| PLX4720 | 147 | 288 | 29 | 57 | 14 |
| Nutlin-3 | 531 | 11 | 96 | 2 | 2 |
| lapatinib | 345 | 1354 | 17 | 66 | 17 |
| Nilotinib | 411 | 615 | 31 | 47 | 22 |
| PHA-665752 | 23 | 0 | 100 | 0 | 0 |
| Erlotinib | 242 | 784 | 19 | 63 | 18 |
| Sorafenib | 37 | 256 | 12 | 83 | 5 |

## Supplementary Figures

**Supplementary Figure 1:**
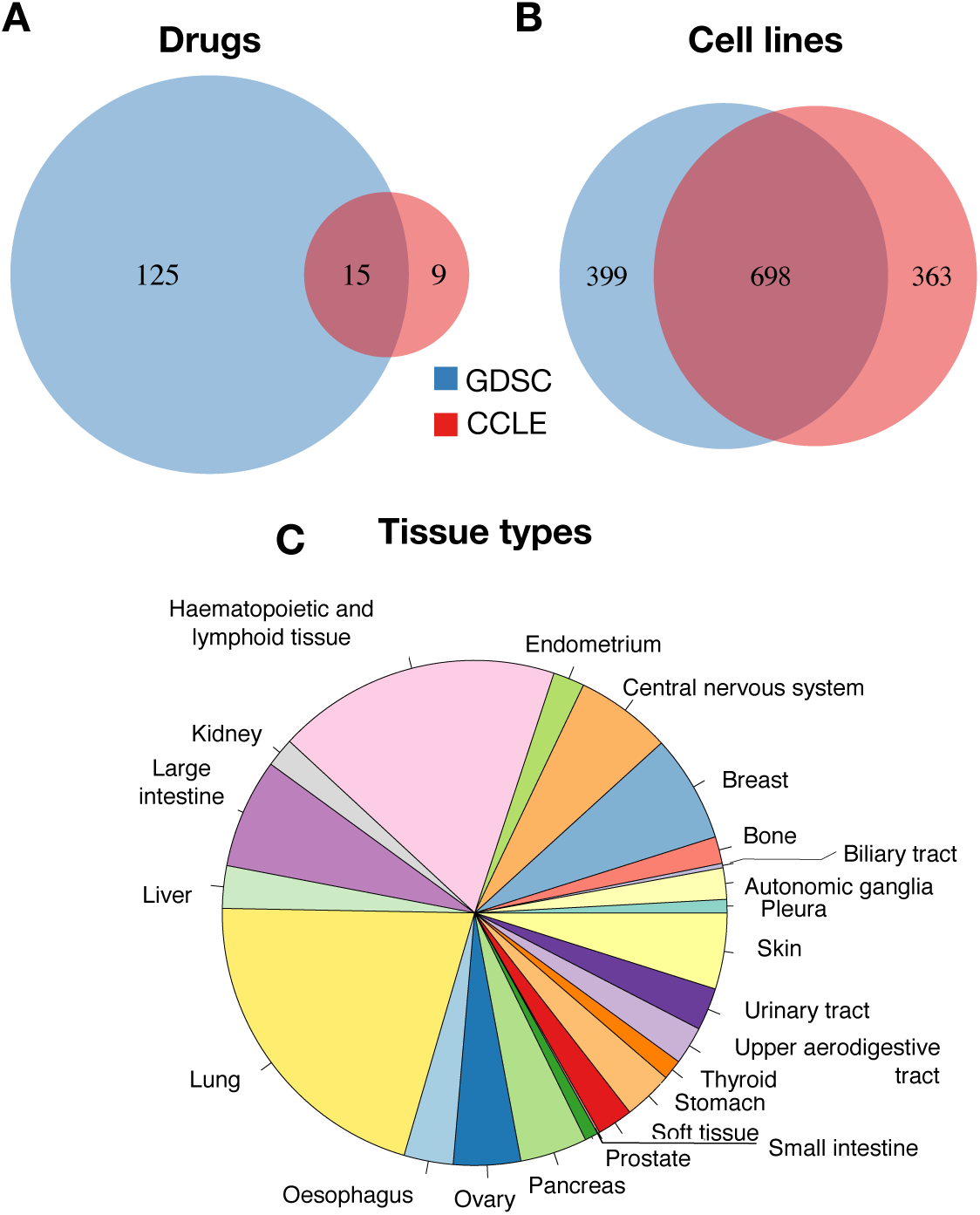
Intersection between GDSC and CCLE. Overlap of (A) drugs, (B) cell lines and (C) tissue types.

**Supplementary Figure 2:**
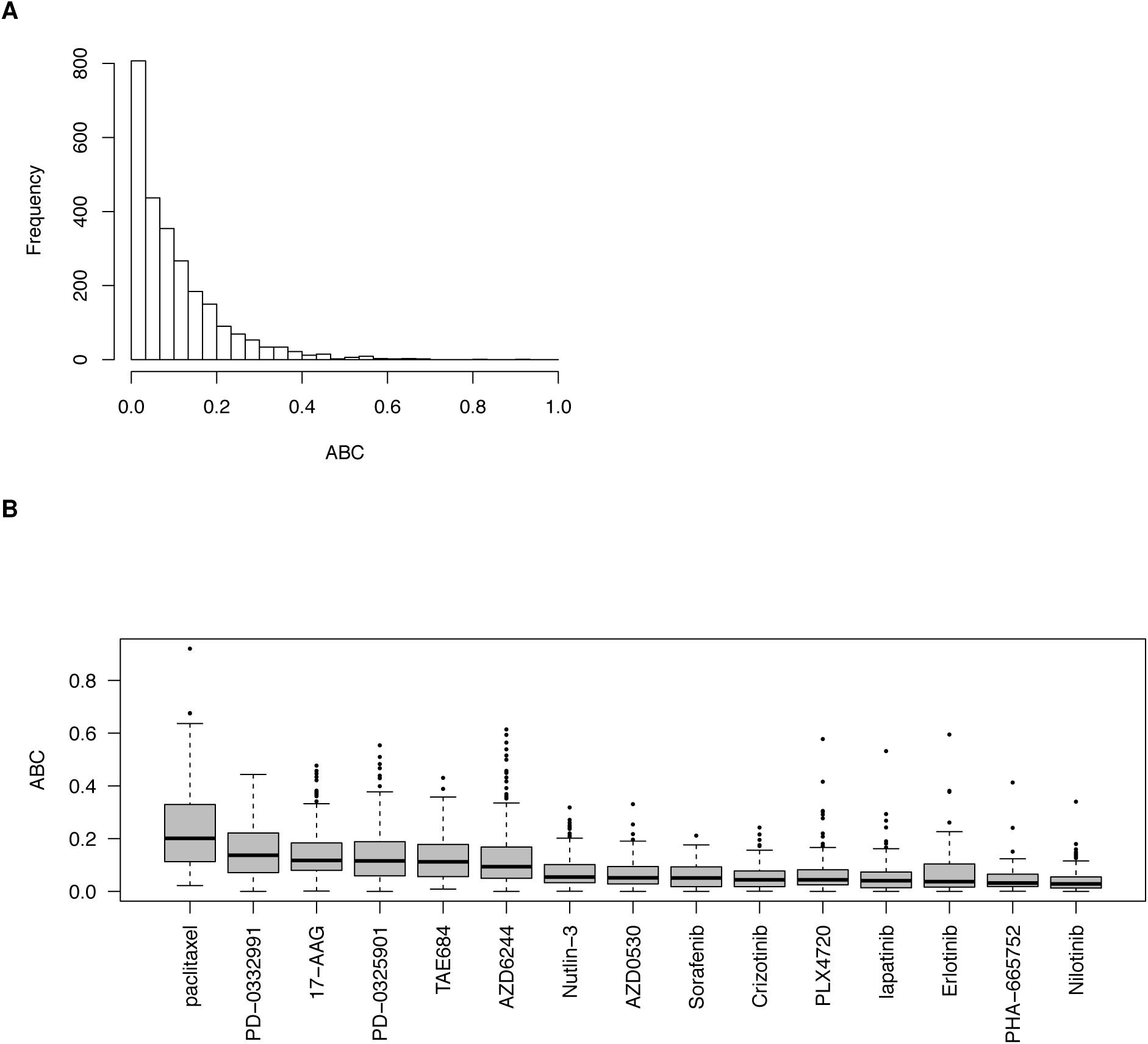
(A) Histogram of ABC estimates for all common drug dose-response curves between GDSC and CCLE. (B) Boxes represent the median and inter quartile range of ABC for drug-cell line combinations screened in GDSC and CCLE.

**Supplementary Figure 3:**
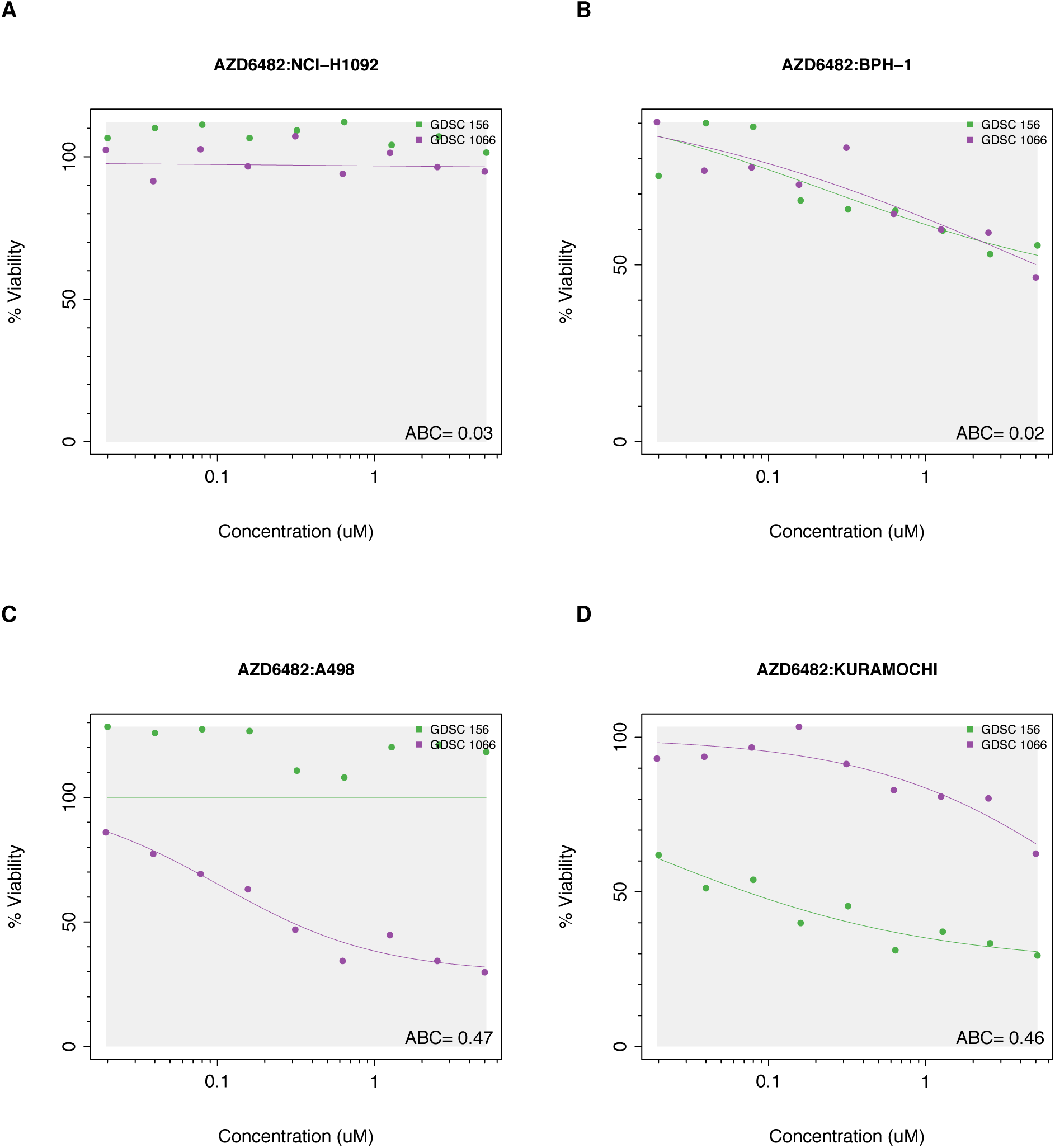
Examples of (A,B) consistent and (C,D) inconsistent replicated experiments screening AZD6482 in GDSC. The grey area represents the common concentration range between studies.

**Supplementary Figure 4:**
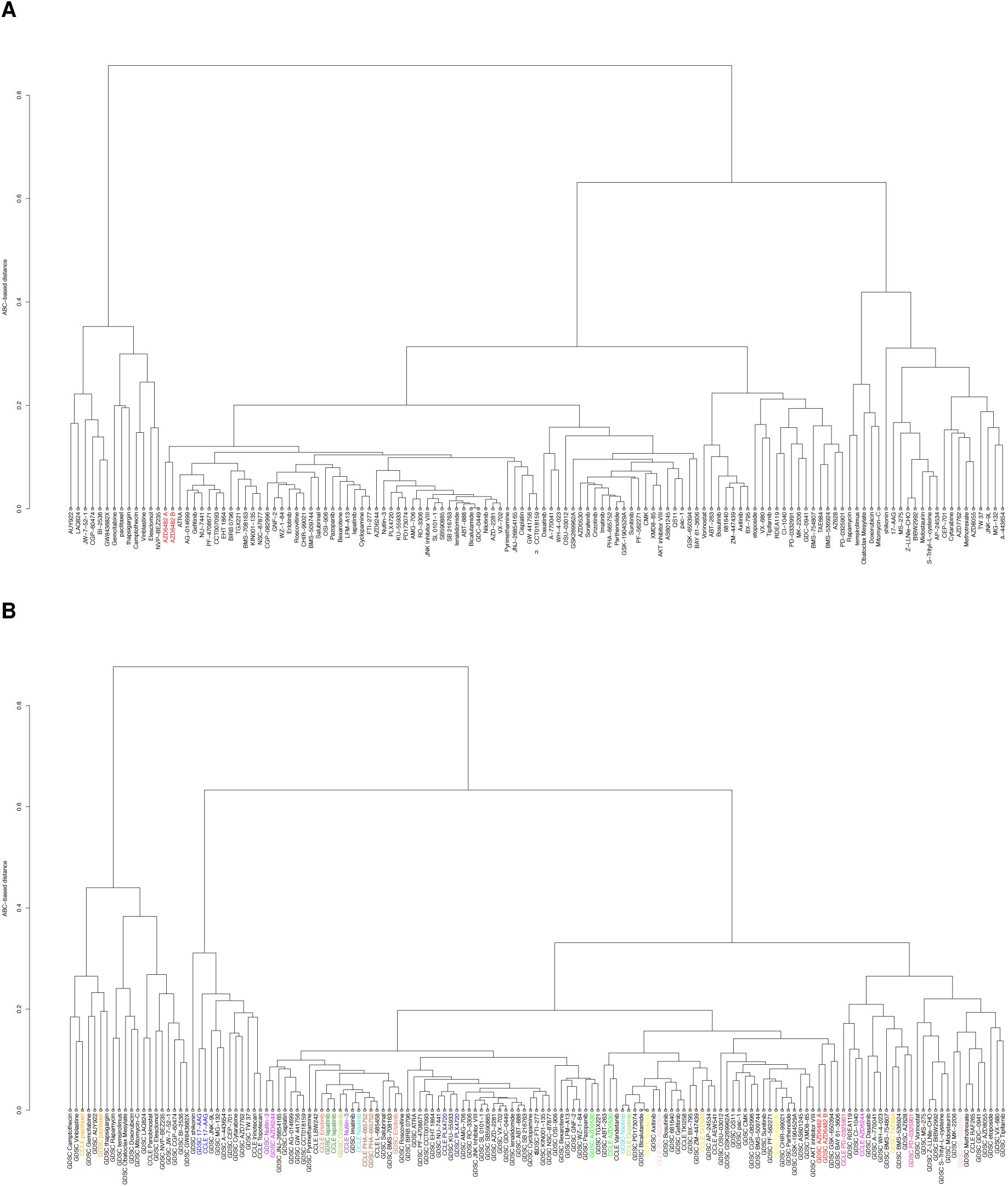
(A) Dendrogram of the clustering of all drugs in GDSC based on their mean ABC values. (B) Dendogram of the clustering of all drugs in CCLE and GDSC based on their mean ABC values, overlapped drugs are shown with the same colour.

**Supplementary Figure 5:**
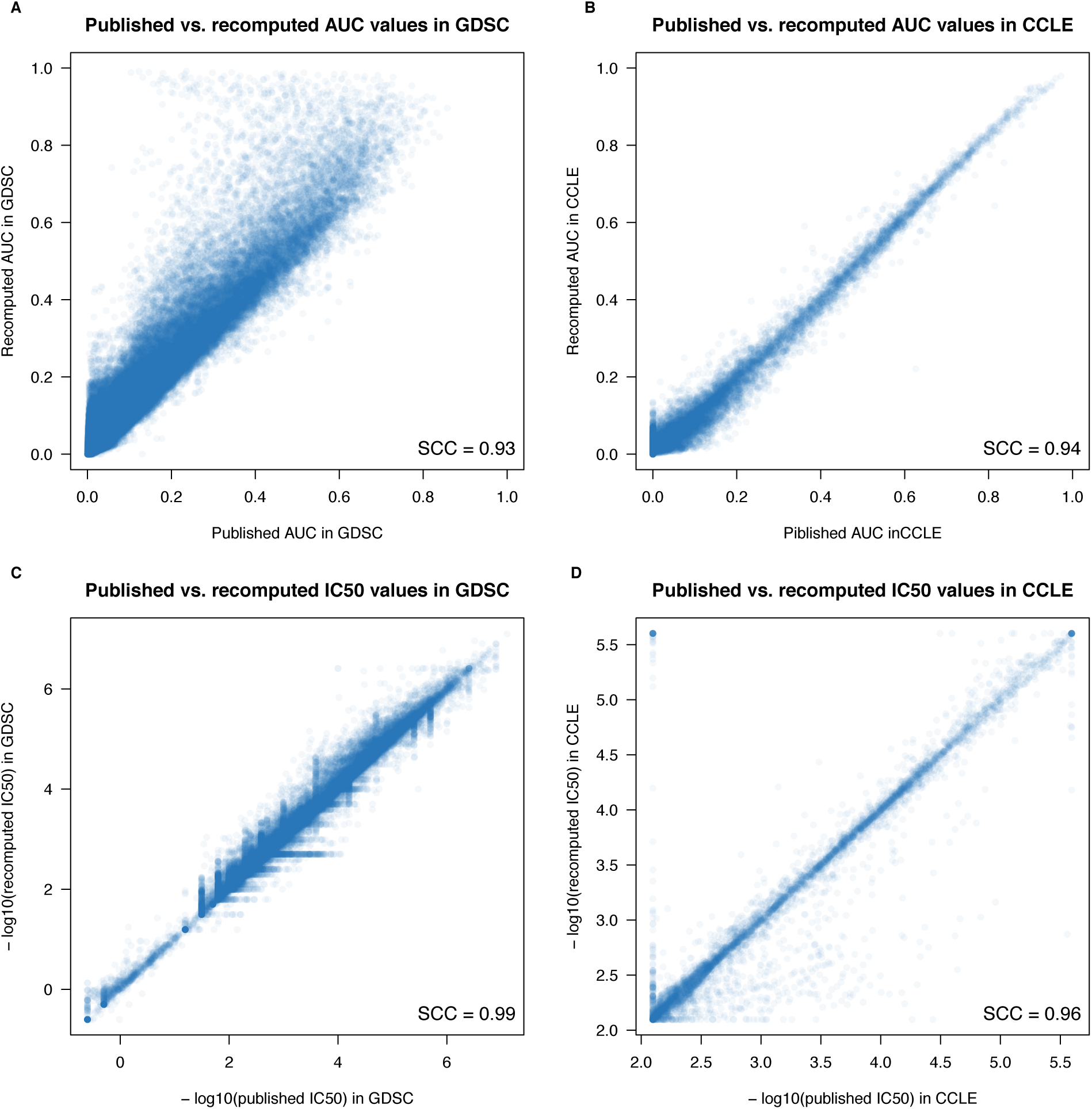
Comparison between published and recomputed drug sensitivity values between GDSC and CCLE. (A) AUC in GDSC; (B) AUC in CCLE; (C) IC_50_ in GDSC; (D) IC_50_ in CCLE. SCC stands for Spearman correlation coefficient.

**Supplementary Figure 6:**
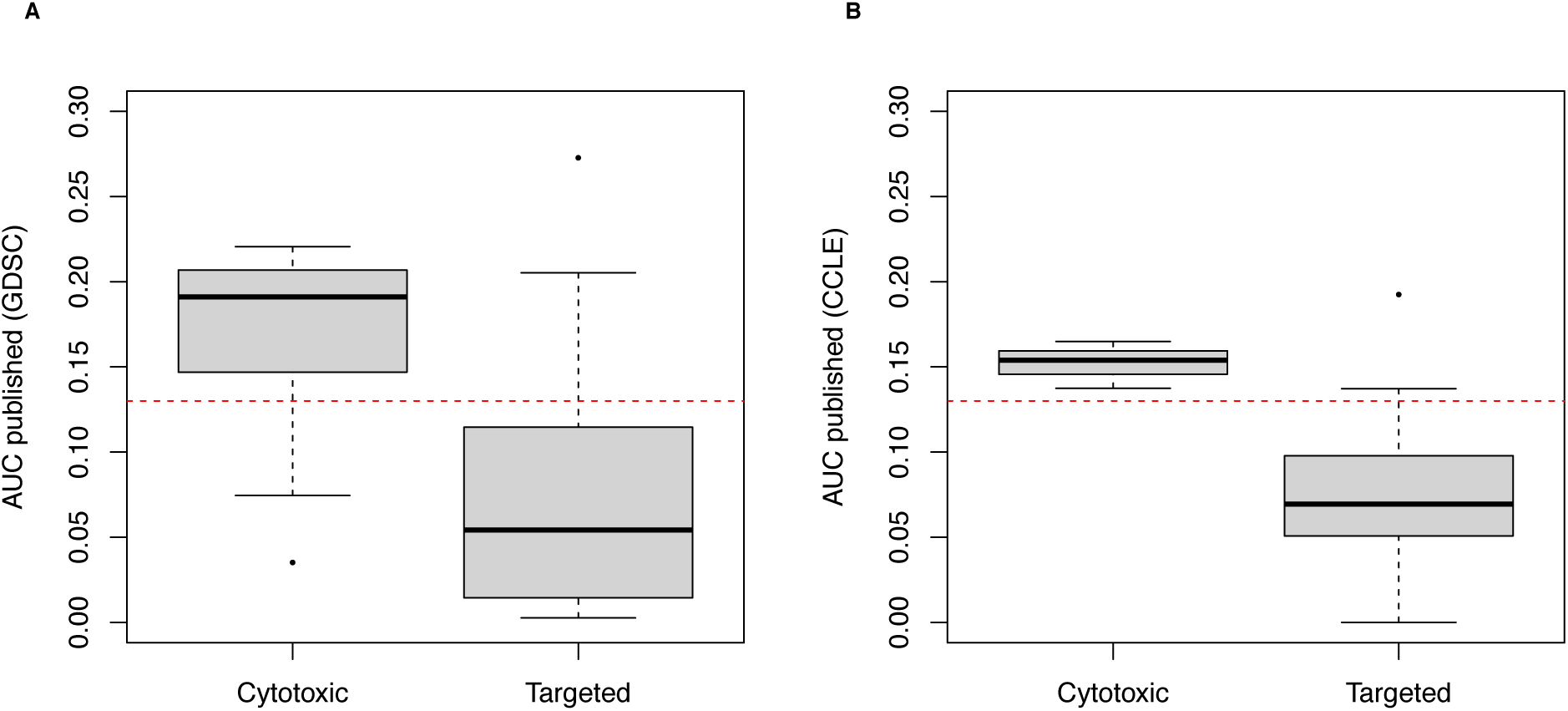
Comparison of median absolute deviation (MAD) of published AUC values between cytotoxic and targeted drugs using all cell lines in (A) GDSC and (B) CCLE.

**Supplementary Figure 7:**
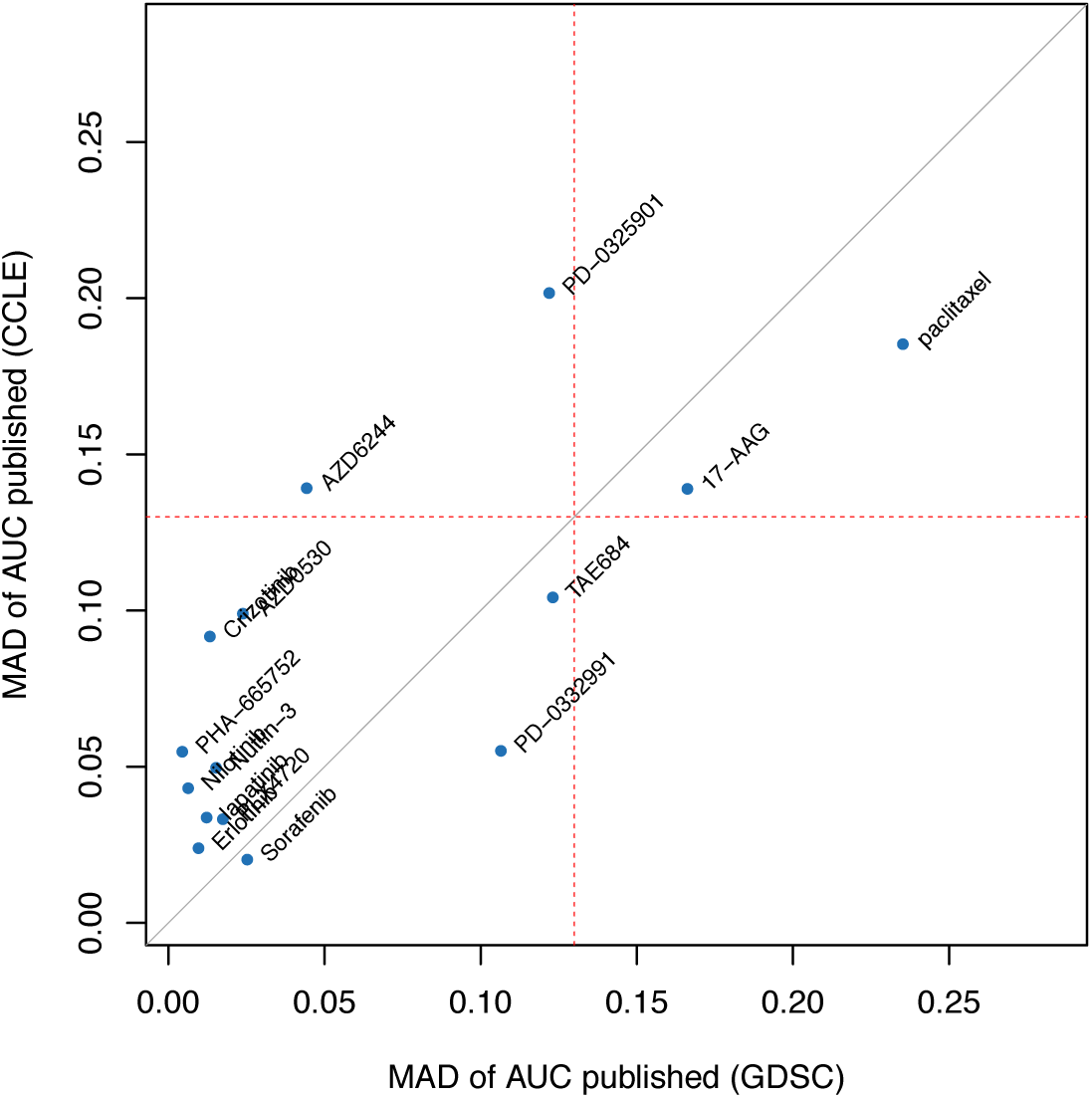
Comparison of median absolute deviation (MAD) of published AUC values between drugs using common cell lines in (A) GDSC and (B) CCLE.

**Supplementary Figure 8:**
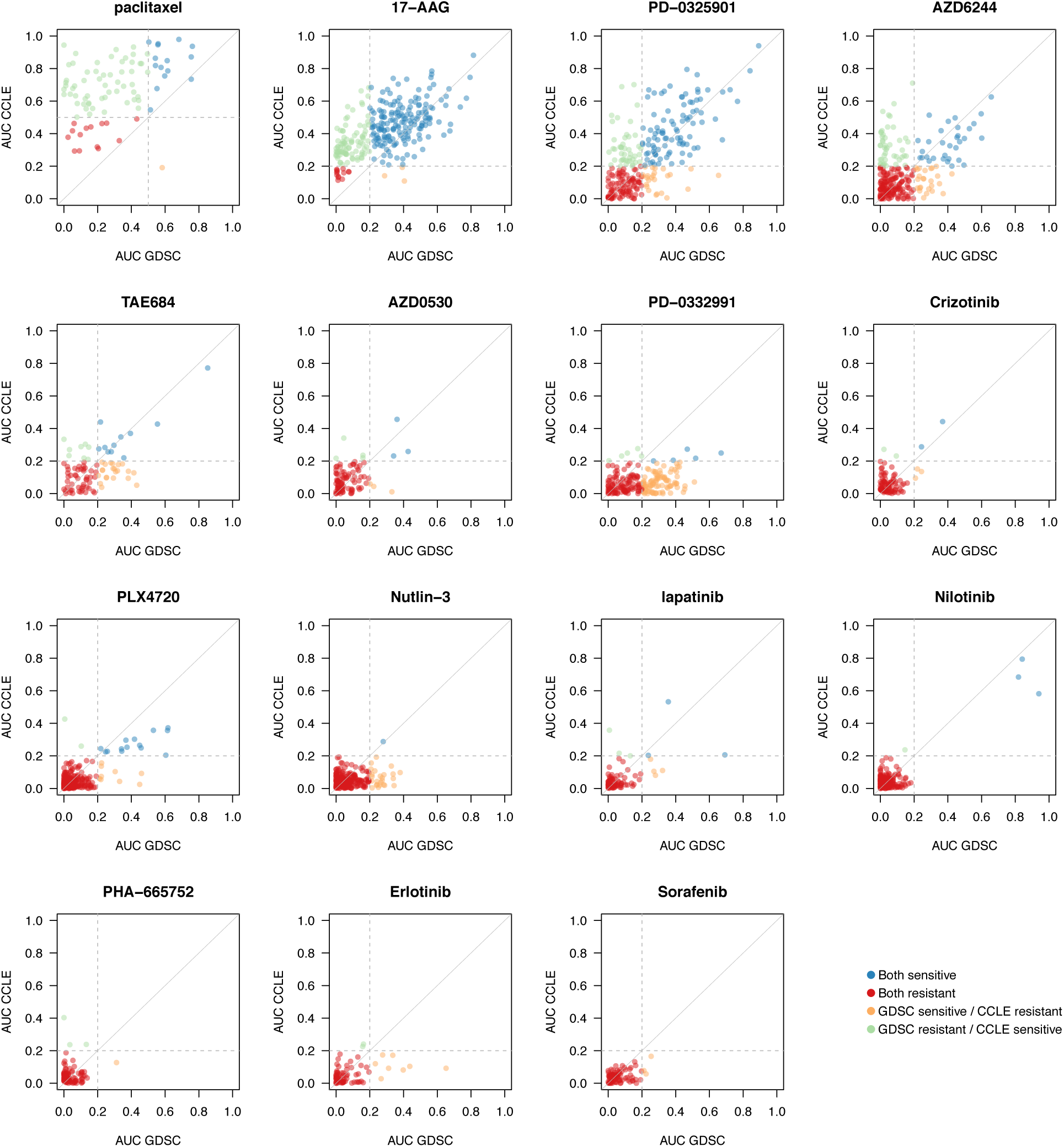
Comparison of AUC values between GDSC and CCLE, as recomputed within *Pharma-coGx*. For cytotoxic drugs (paclitaxel), cell lines with AUC < 0.4 were considered as resistant, while for targeted therapies cell lines with AUC < 0.2 were considered resistant (grey dashed lines). In case of perfect consistency, all points would lie on the grey diagonal.

**Supplementary Figure 9:**
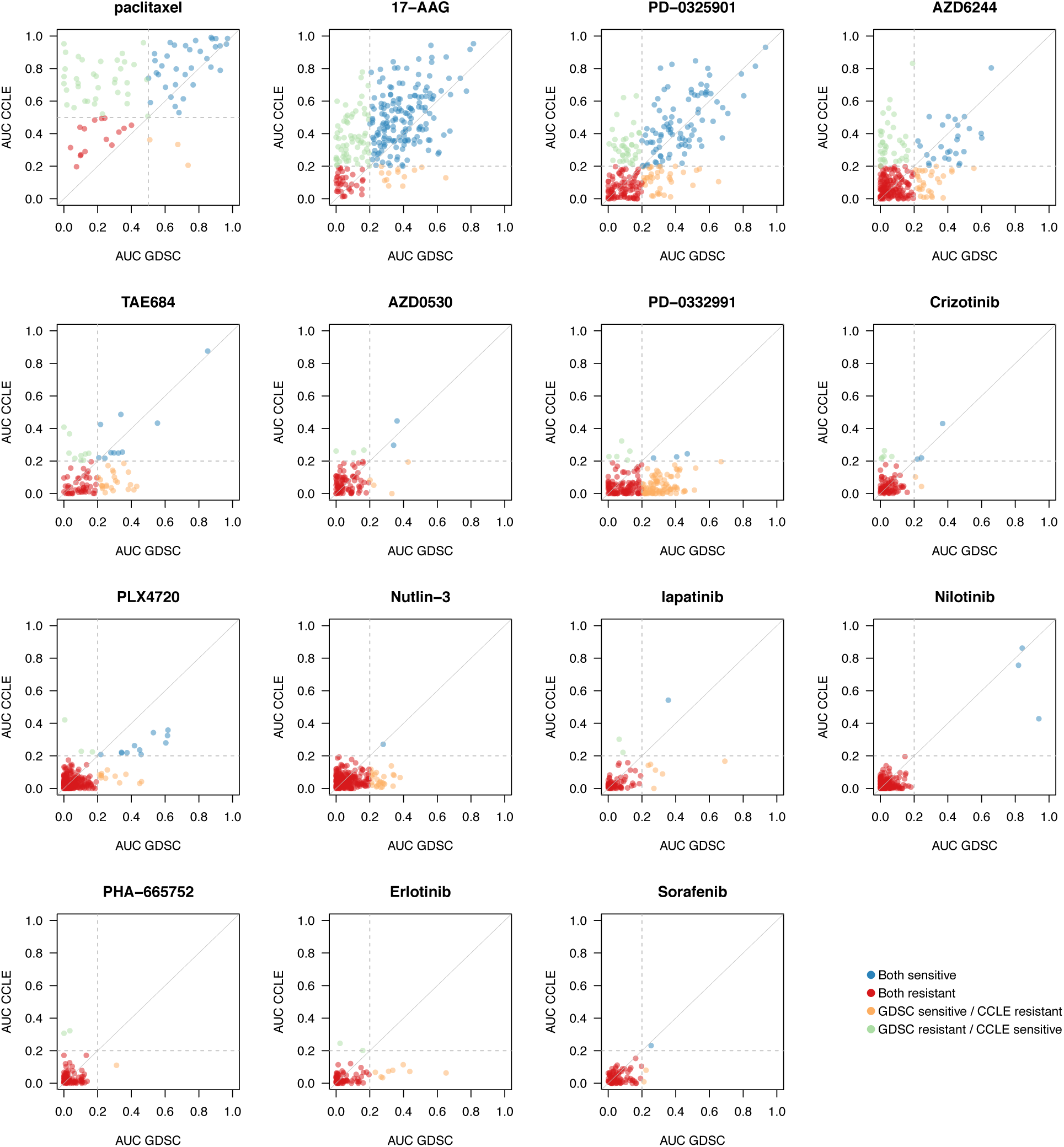
Comparison of AUC* values between GDSC and CCLE, as recomputed within *Pharma-coGx*. For cytotoxic drugs (paclitaxel), cell lines with AUC* < 0.4 were considered as resistant, while for targeted therapies cell lines with AUC* < 0.2 were considered resistant (grey dashed lines). In case of perfect consistency, all points would lie on the grey diagonal.

**Supplementary Figure 10:**
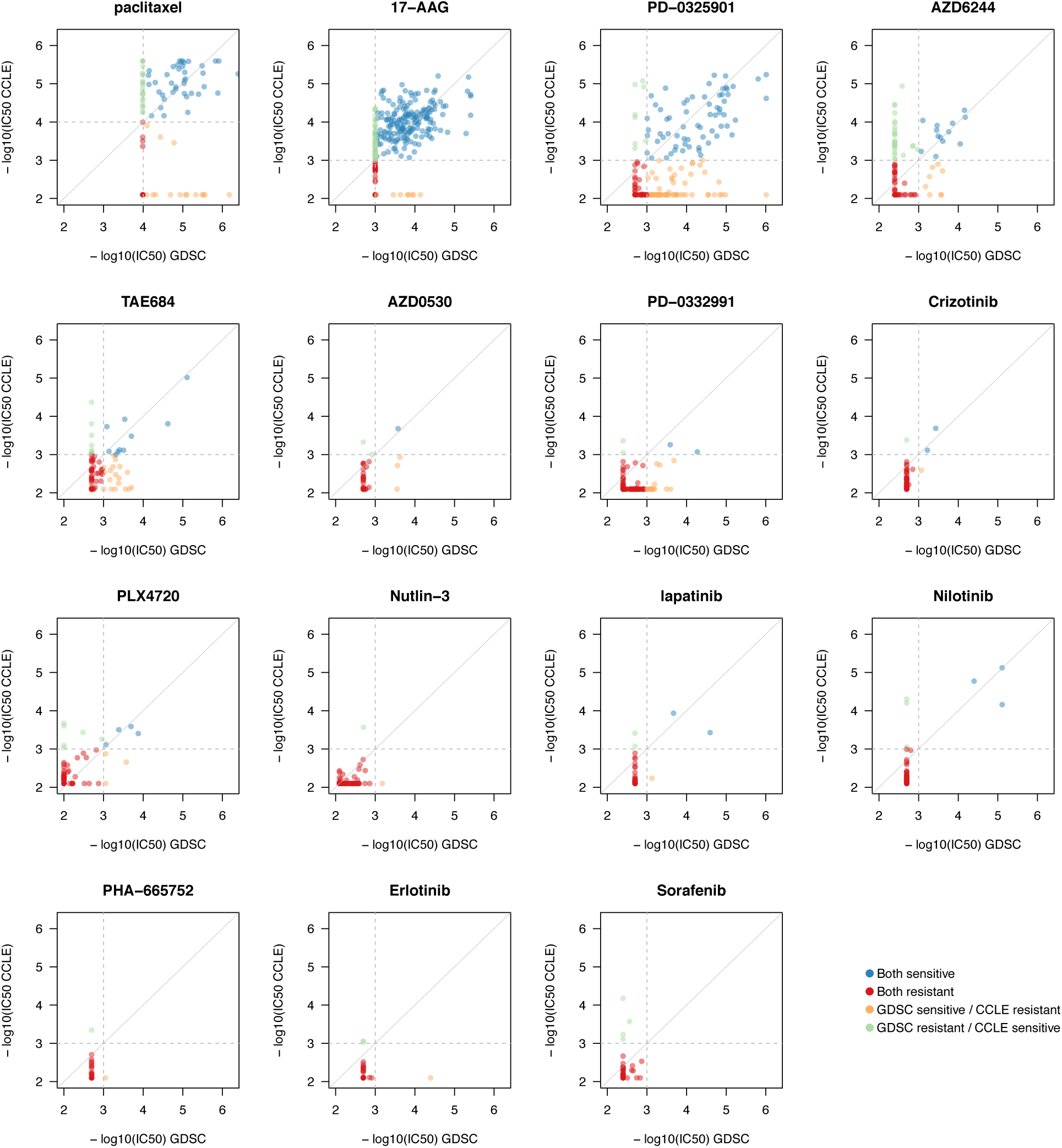
Consistency of IC_50_ values between GDSC and CCLE, as published. For cytotoxic drugs (paclitaxel), cell lines with IC_50_ ≤ 10*μ*M were considered as resistant, while for targeted therapies cell lines with IC_50_ ≤ 1*μ*M were considered resistant (grey dashed lines). In case of perfect consistency, all points would lie on the grey diagonal.

**Supplementary Figure 11:**
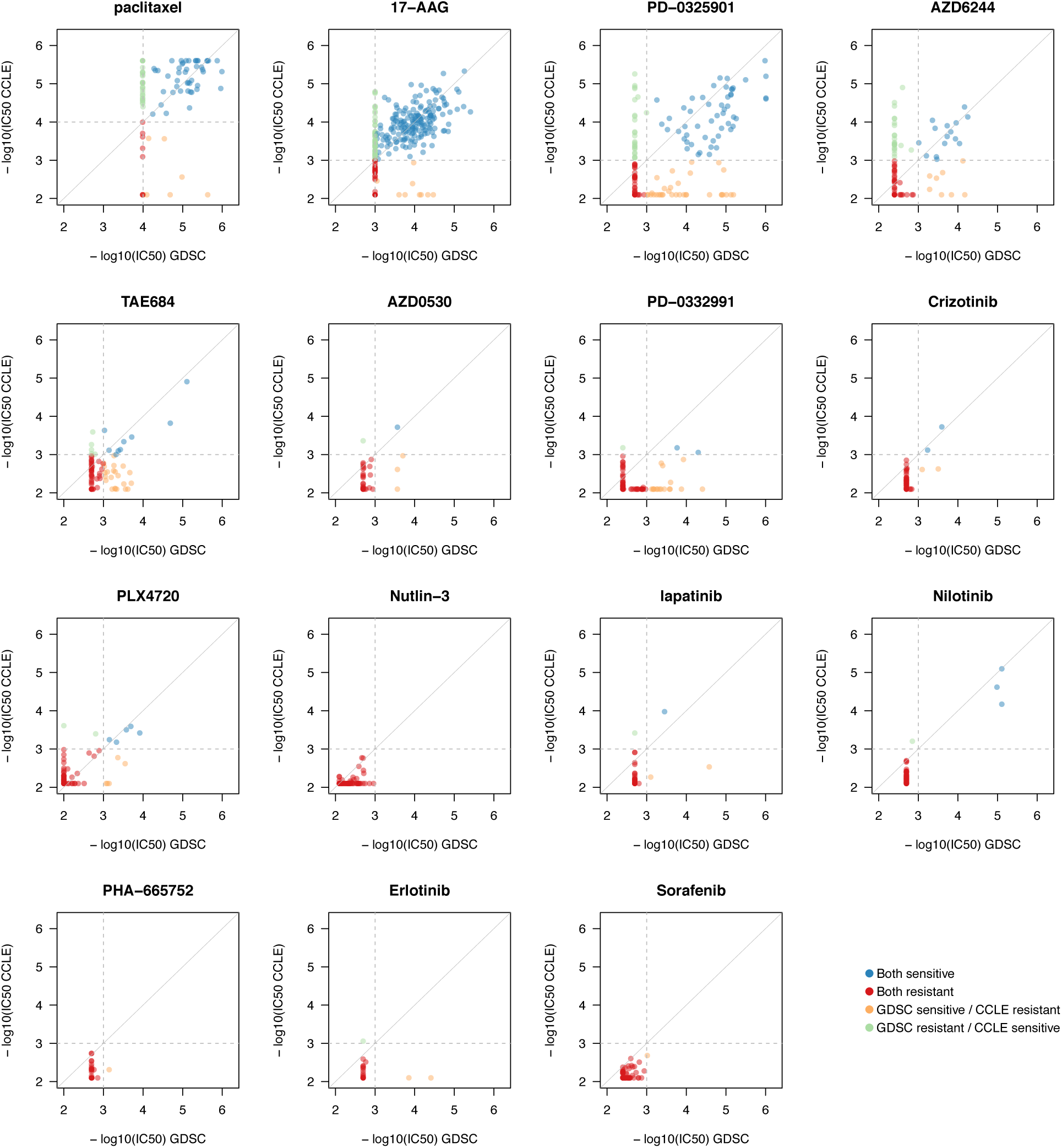
Consistency of IC_50_ values between GDSC and CCLE, as recomputed within *Phar-macoGx*. For cytotoxic drugs (paclitaxel), cell lines with IC_50_ ≤ 10*μ*M were considered as resistant, while for targeted therapies cell lines with IC_50_ ≤ 1*μ*M were considered resistant (grey dashed lines). In case of perfect consistency, all points would lie on the grey diagonal.

**Supplementary Figure 12:**
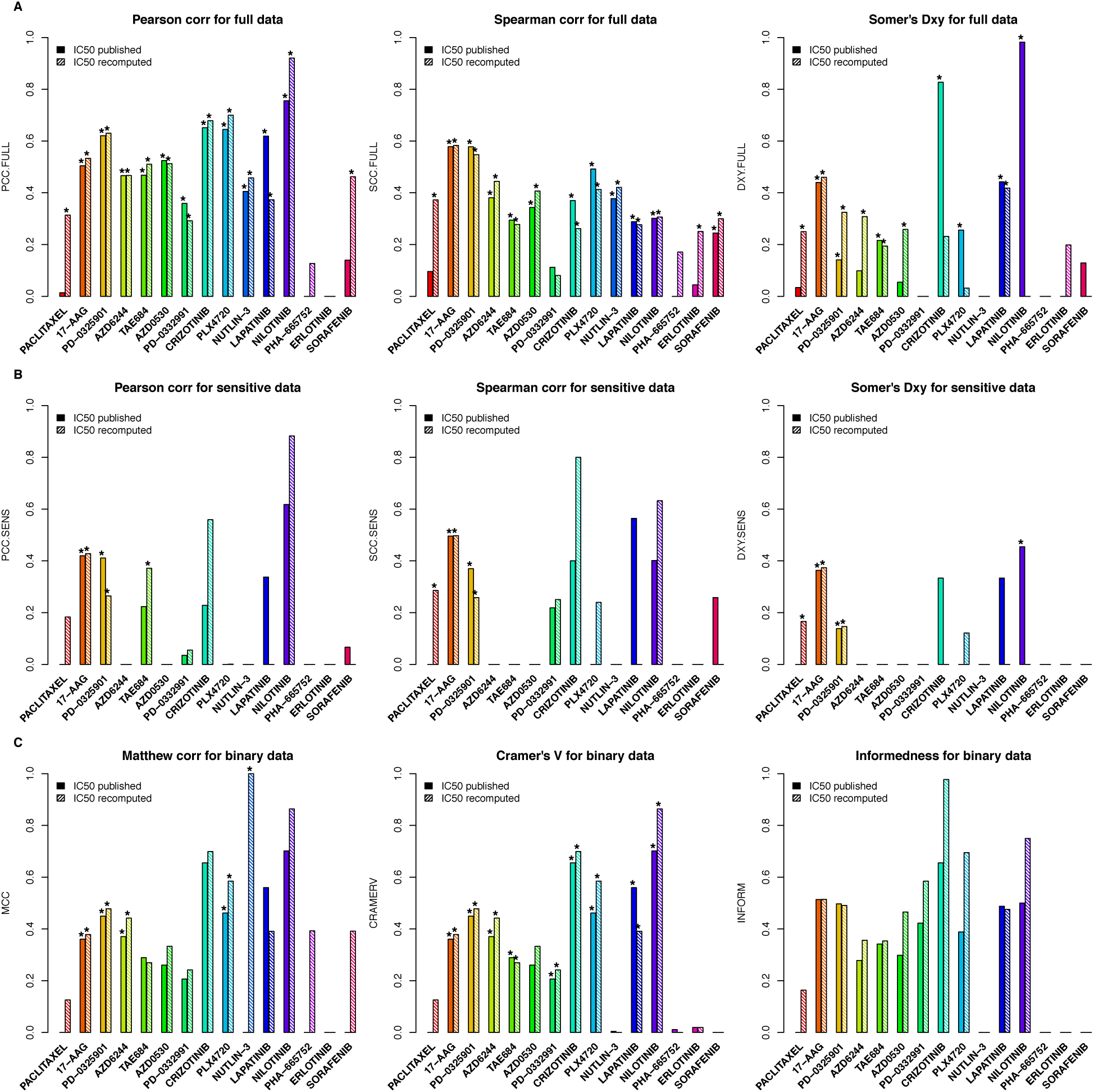
Consistency of IC_50_ values between GDSC and CCLE, as published and recomputed within *PharmacoGx*.

**Supplementary Figure 13:**
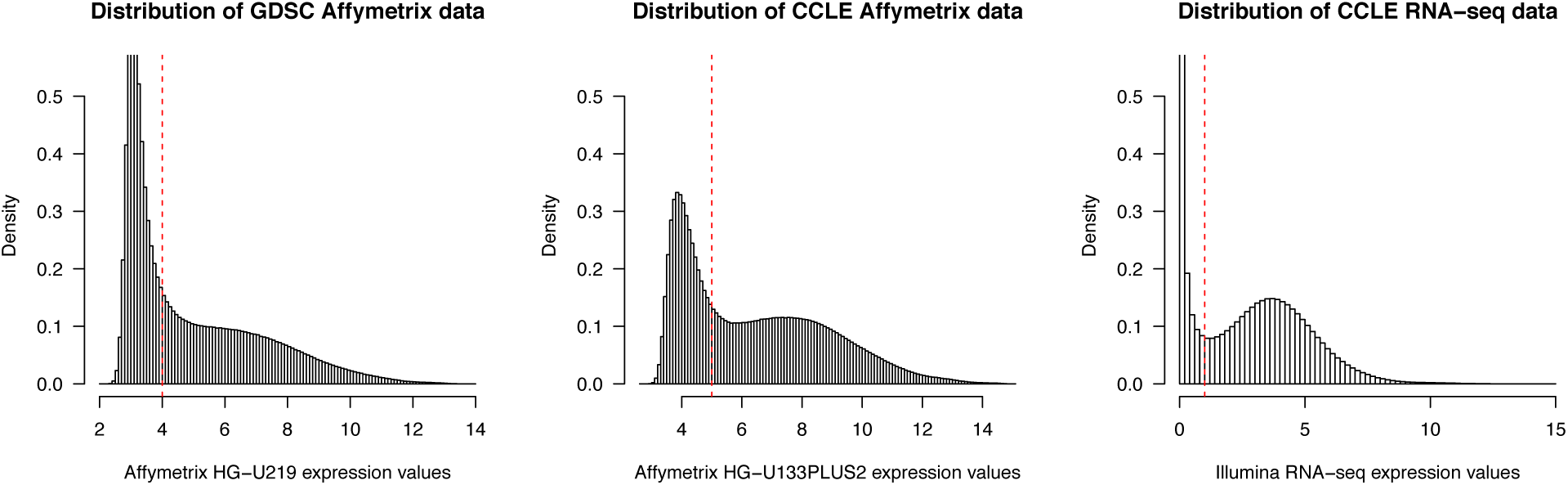
Distribution of gene expression values and corresponding cutoffs for the microarray Affymetrix HG-U219 platform in GDSC (cutoff = 4), the microarray Affymetrix HG-U133PLUS2 platform in CCLE (cutoff = 5) and the new Illumina RNA-seq data in CCLE (cutoff = 1).

**Supplementary Figure 14:**
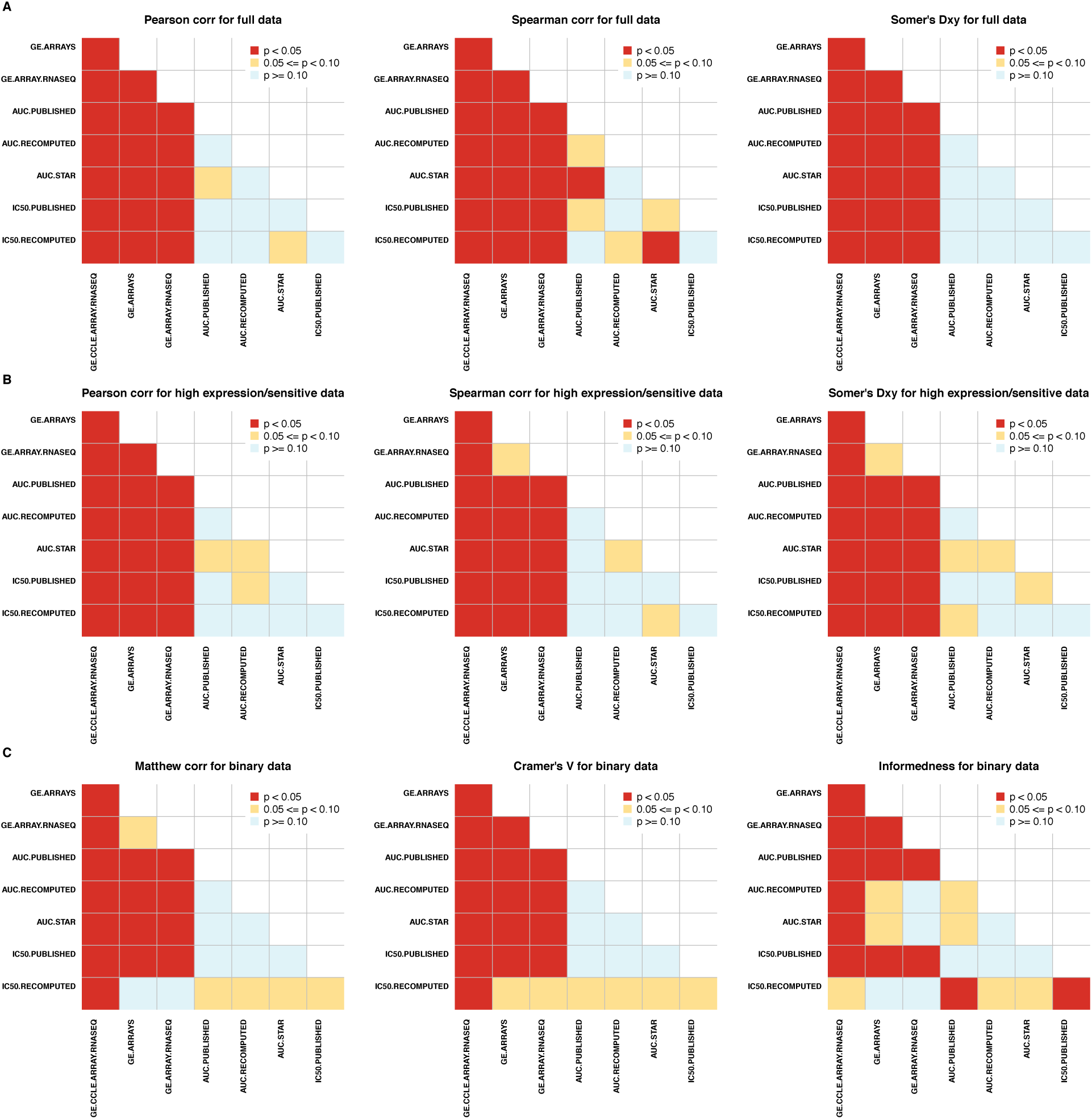
Statistical test for difference in consistency for gene expression and drug sensitivity data. Each cell in the matrix represents the p-value (coded by colour) for a given pairwise comparison for consistency values. For instance, consistency of gene expression data is statistically significantly higher than consistency of drug sensitivity data. GE.CCLE.ARRAY.RNASEQ: Consistency between gene expression data generated using Affymetrix HG-U133PLUS2 microarray and Illumina RNA-seq platforms within CCLE; GE.ARRAYS: Consistency between gene expression data generated using Affymetrix HG-U133A and HG-U133PLUS2 microarray platforms in GDSC and CCLE, respectively; GE.ARRAY.RNASEQ: Consistency between gene expression data generated using Affymetrix HG-U133PA microarray and Illumina RNA-seq platforms in GDSC and CCLE, respectively; AUC.PUBLISHED: Consistency of AUC values as published in GDSC and CCLE; AUC.PUBLISHED: Consistency of AUC values as published in GDSC and CCLE; AUC.RECOMPUTED: Consistency of AUC values in GDSC and CCLE as recomputed using our *PharmacoGx* tool; AUC.STAR: Consistency of AUC values in GDSC and CCLE as recomputed from the common concentration range using our *PharmacoGx* tool; IC50.PUBLISHED: Consistency of IC_50_ values as published in GDSC and CCLE; IC50.RECOMPUTED: Consistency of IC_50_ values in GDSC and CCLE as recomputed using our *PharmacoGx* tool.

**Supplementary Figure 15:**
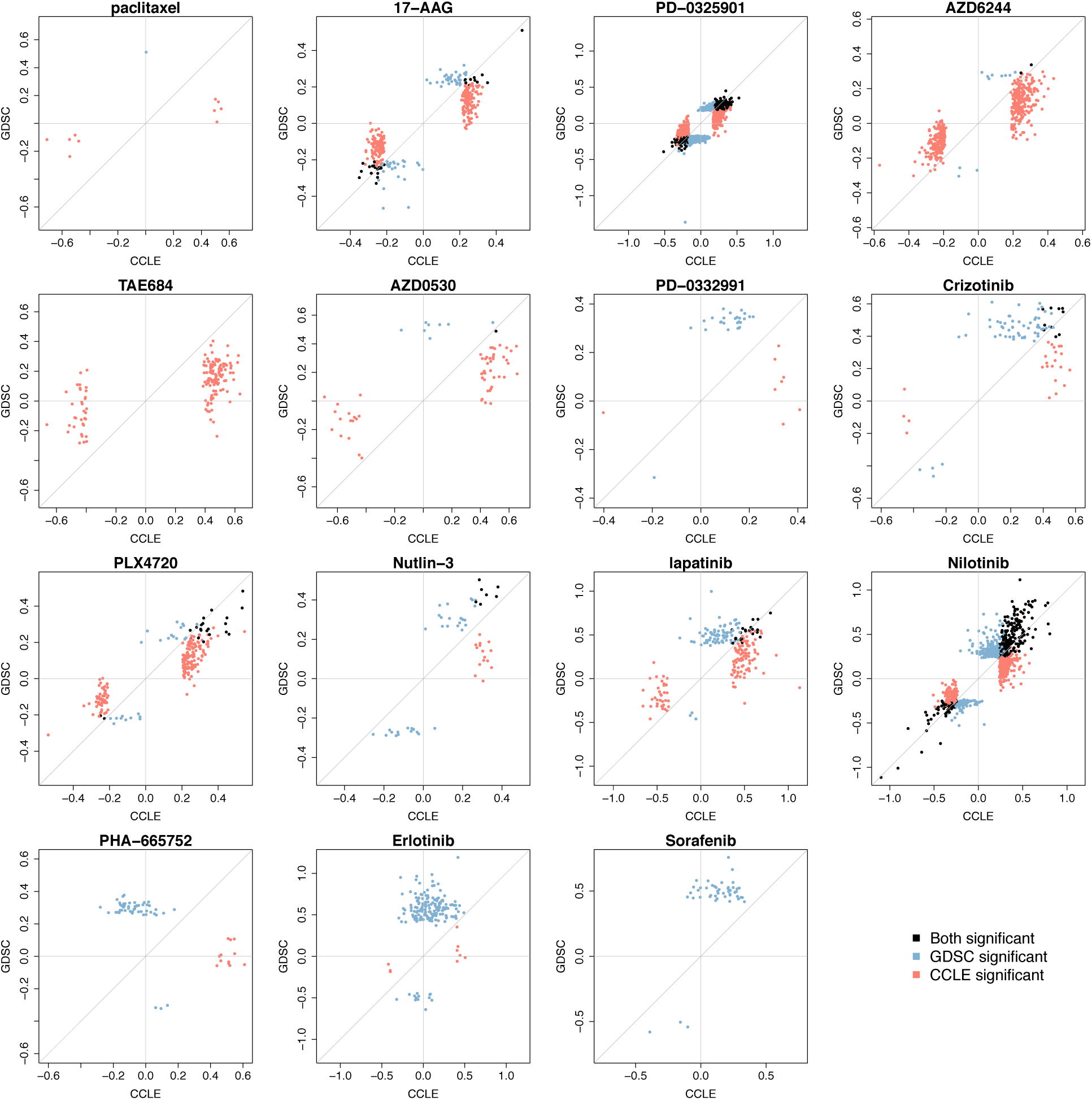
Scatterplot representing the effect size of the significant gene-drug associations (FDR < 5%) identified using continuous AUC and the common cell lines screened both in GDSC and CCLE. Gene-drug associations are identified using gene expression data and continuous published AUC as input and output of a linear model, respectively. In case of perfect consistency, all points would lie on the grey diagonal.

**Supplementary Figure 16:**
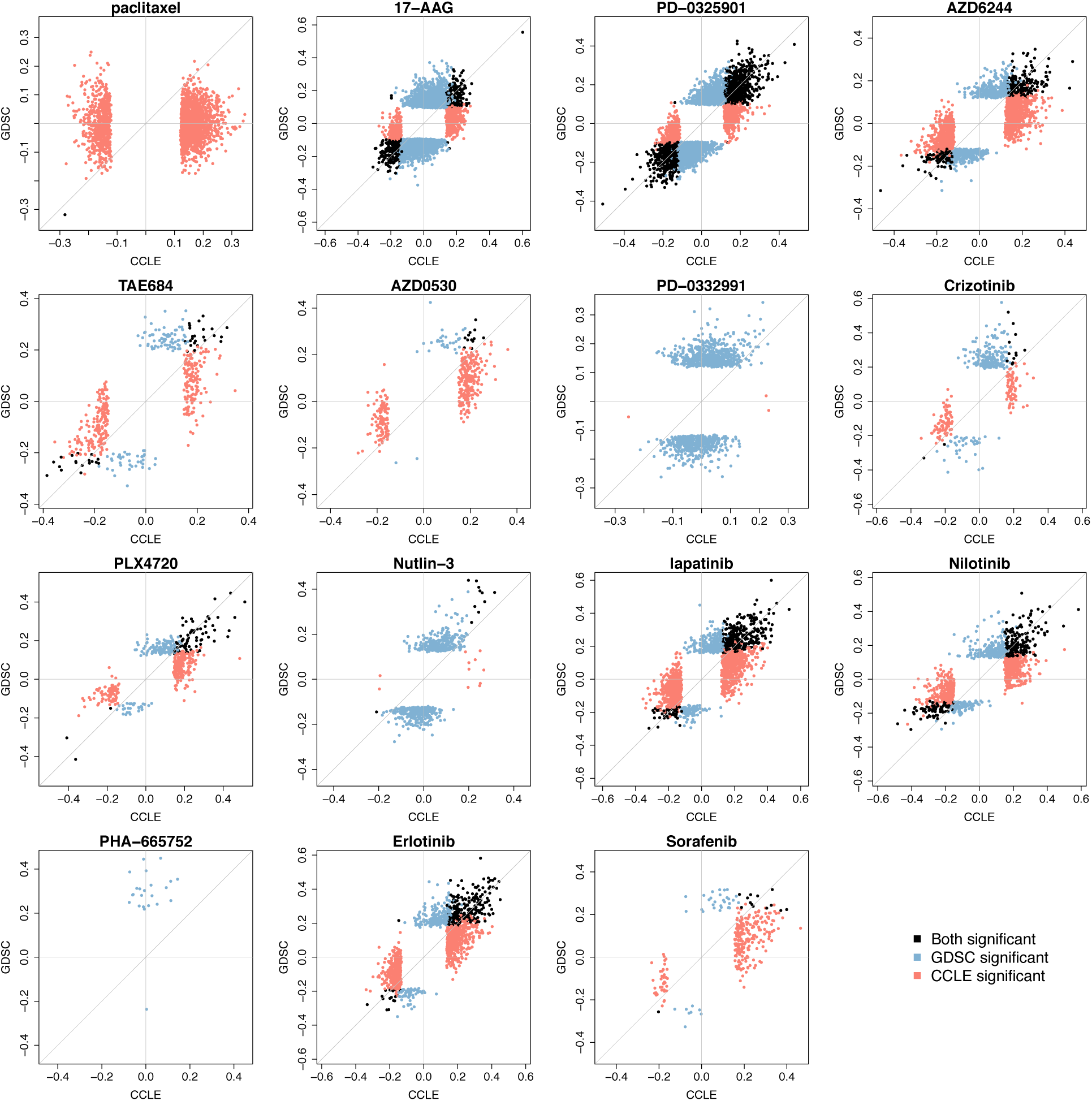
Scatterplot representing the effect size of the significant gene-drug associations (FDR < 5%) identified using continuous AUC and all cell lines screened in each study. Gene-drug associations are identified using gene expression data and continuous published AUC as input and output of a linear model, respectively. In case of perfect consistency, all points would lie on the grey diagonal.

**Supplementary Figure 17:**
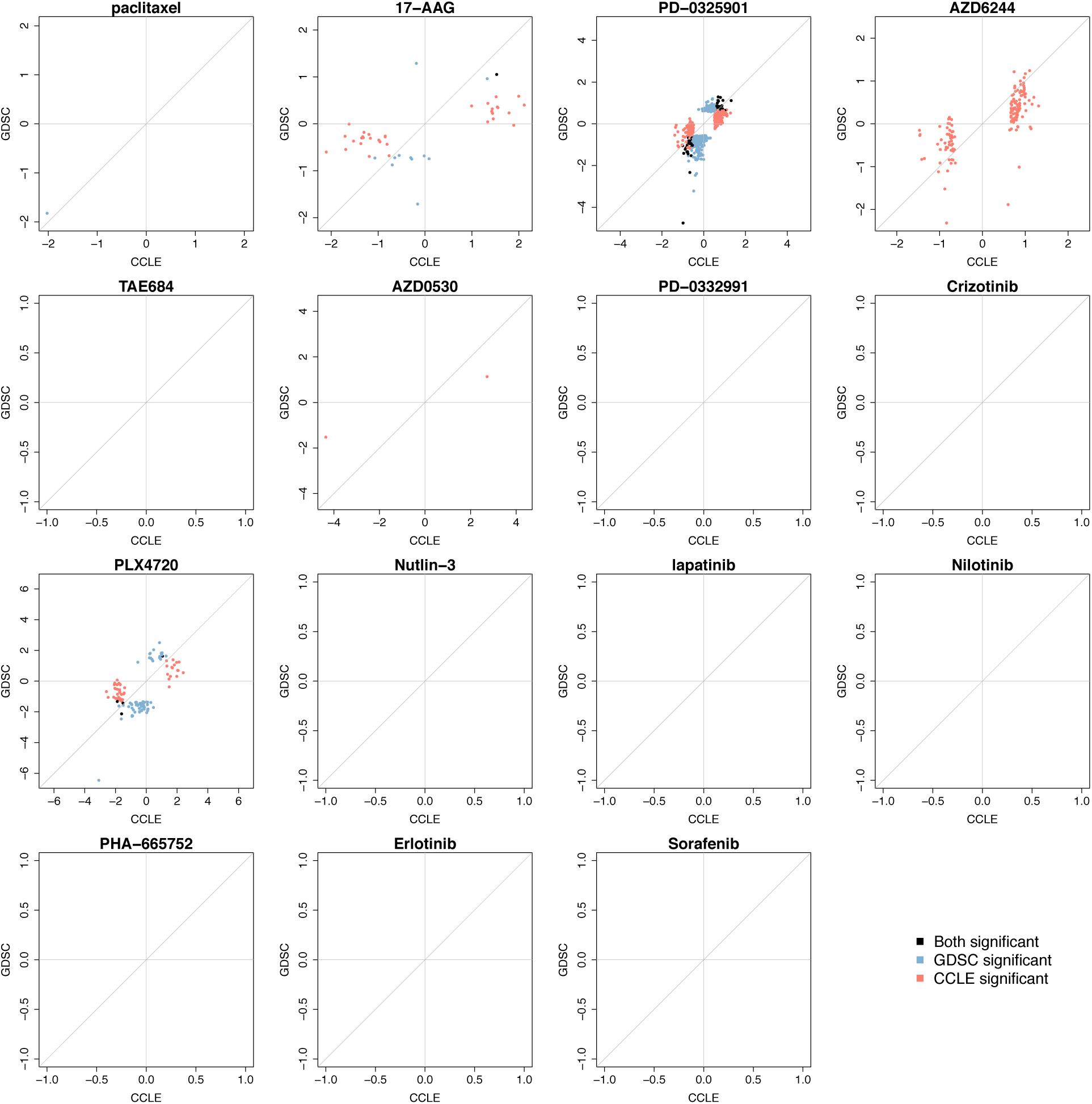
Scatterplot representing the effect size of the significant gene-drug associations (FDR < 5%) identified using discretized AUC and the common cell lines screened both in GDSC and CCLE. Gene-drug associations are identified using gene expression data and discretized published AUC as input and output of a linear model, respectively. Note that the small number of cell lines classified as “sensitive” did not allow for finding enough significant gene-drug associations for the majority of the drugs. This is due to the lack of convergence of the logistic regression model when 3 or less cell lines are in one category.

**Supplementary Figure 18:**
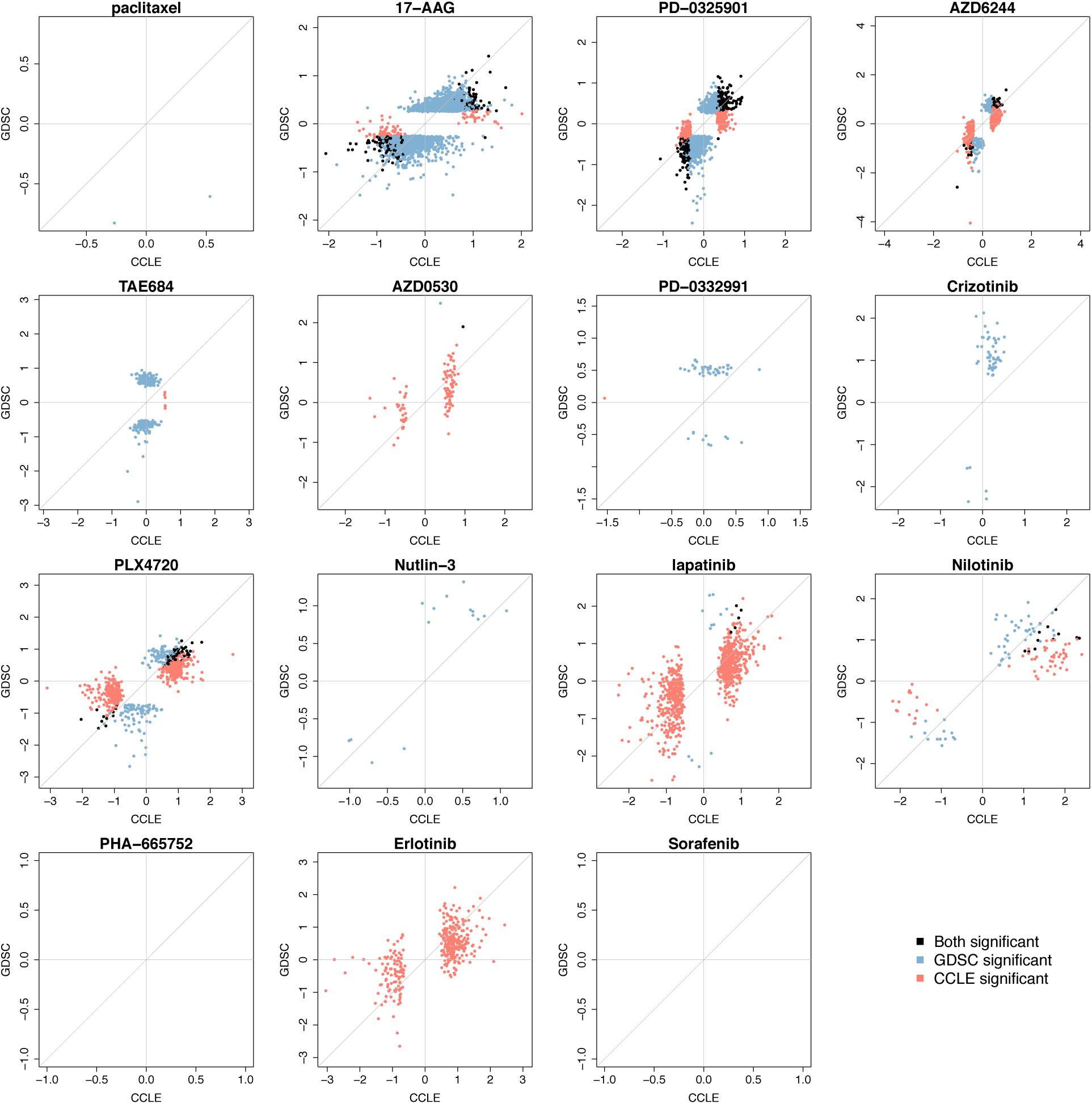
Scatterplot representing the effect size of the significant gene-drug associations (FDR < 5%) identified using discretized AUC and all cell lines screened in each study. Gene-drug associations are identified using gene expression data and discretized published AUC as input and output of a linear model, respectively. Note that the small number of cell lines classified as “sensitive” did not allow for finding enough significant gene-drug associations for PHA-665752 and sorafenib. This is due to the lack of convergence of the logistic regression model when 3 or less cell lines are in one category.

**Supplementary Figure 19:**
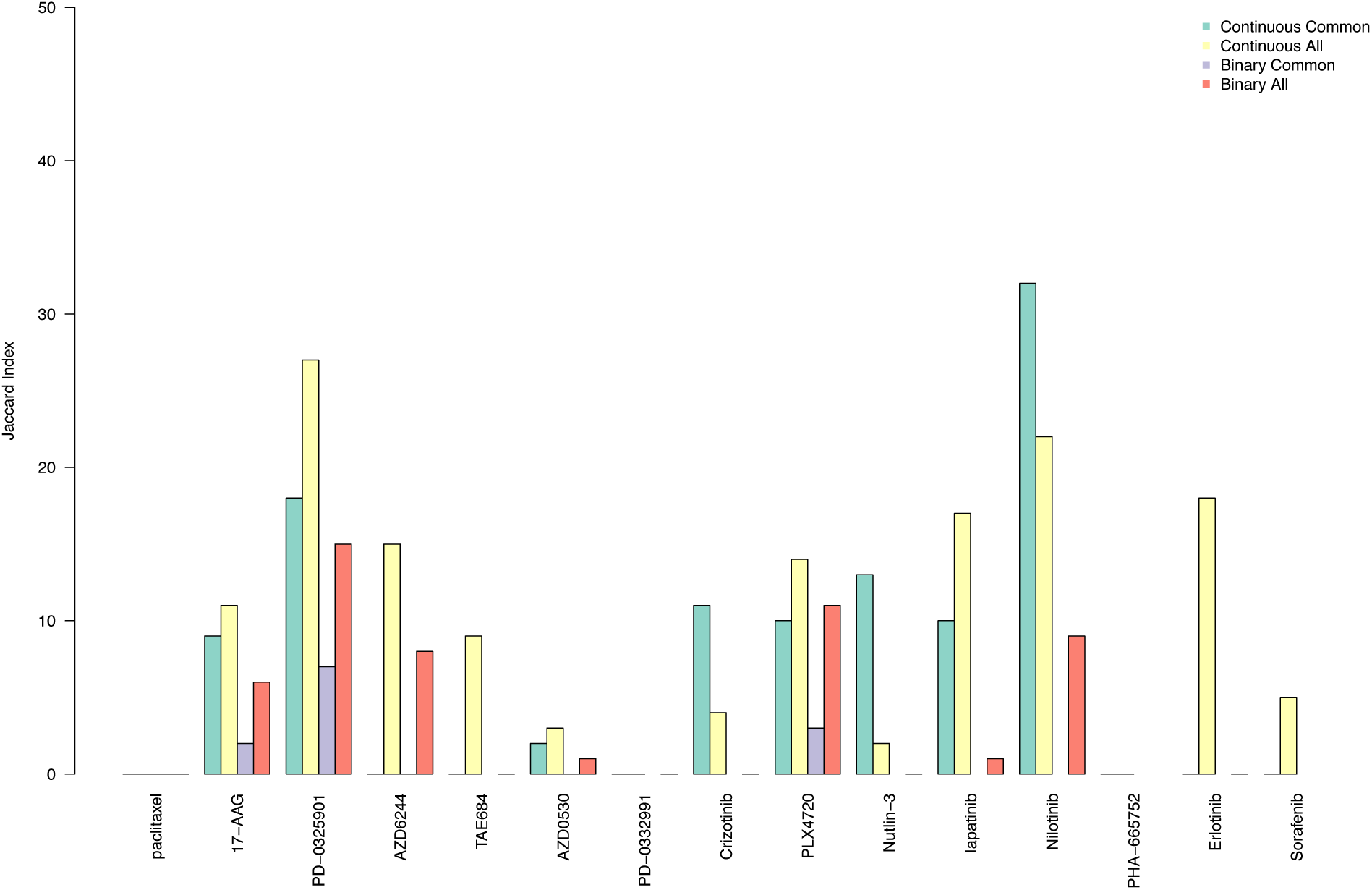
Barplot representing the overlap, as estimated by the Jaccard index, between expression-based gene-drug associations found in GDSC and CCLE. ’Continuous Common’ refers to the associations identified using continuous published AUC values on the common cell lines in GDSC and CCLE; ‘Continuous All’ refers to the associations identified using continuous published AUC values on the entire panel of cell lines screened in each study; ’Binary Common’ refers to the associations identified using the discretized (binary) published AUC values on the common cell lines in GDSC and CCLE; ’Binary All’ refers to the associations identified using the discretized (binary) published AUC values on the entire panels of cell lines screened in each study

**Supplementary Figure 20:**
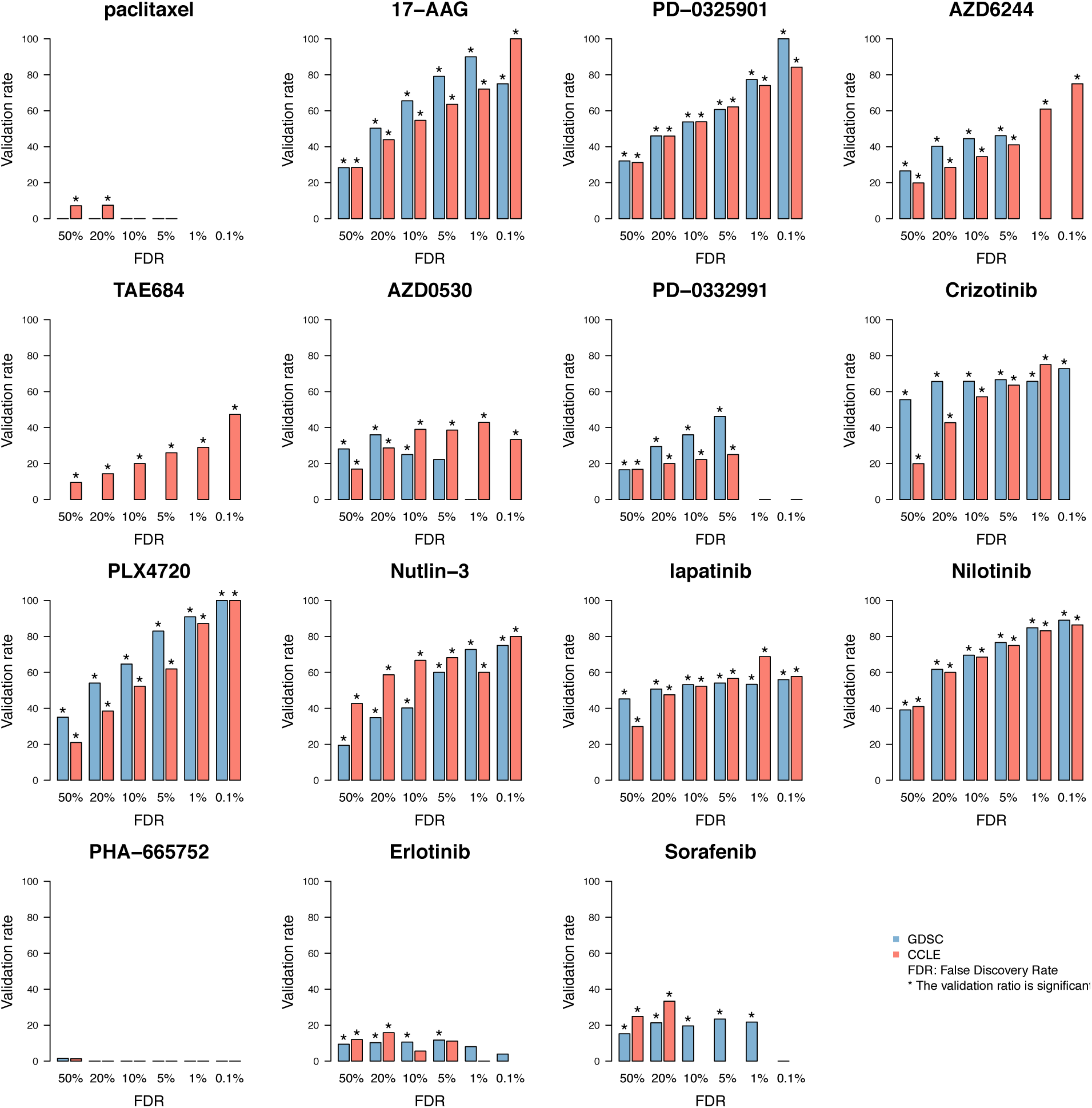
Proportion of validated biomarkers with decreasing FDR using common cell lines screened both in GDSC and CCLE. Gene-drug associations are identified using gene expression data and continuous published AUC as input and output of a linear mode, respectively. The symbol ’*’ represents the significance of the proportion of validated gene-drug associations, computed as the frequency of 1000 random subsets of markers of the same size having equal or greater validation rate compared to the observed rate.

**Supplementary Figure 21:**
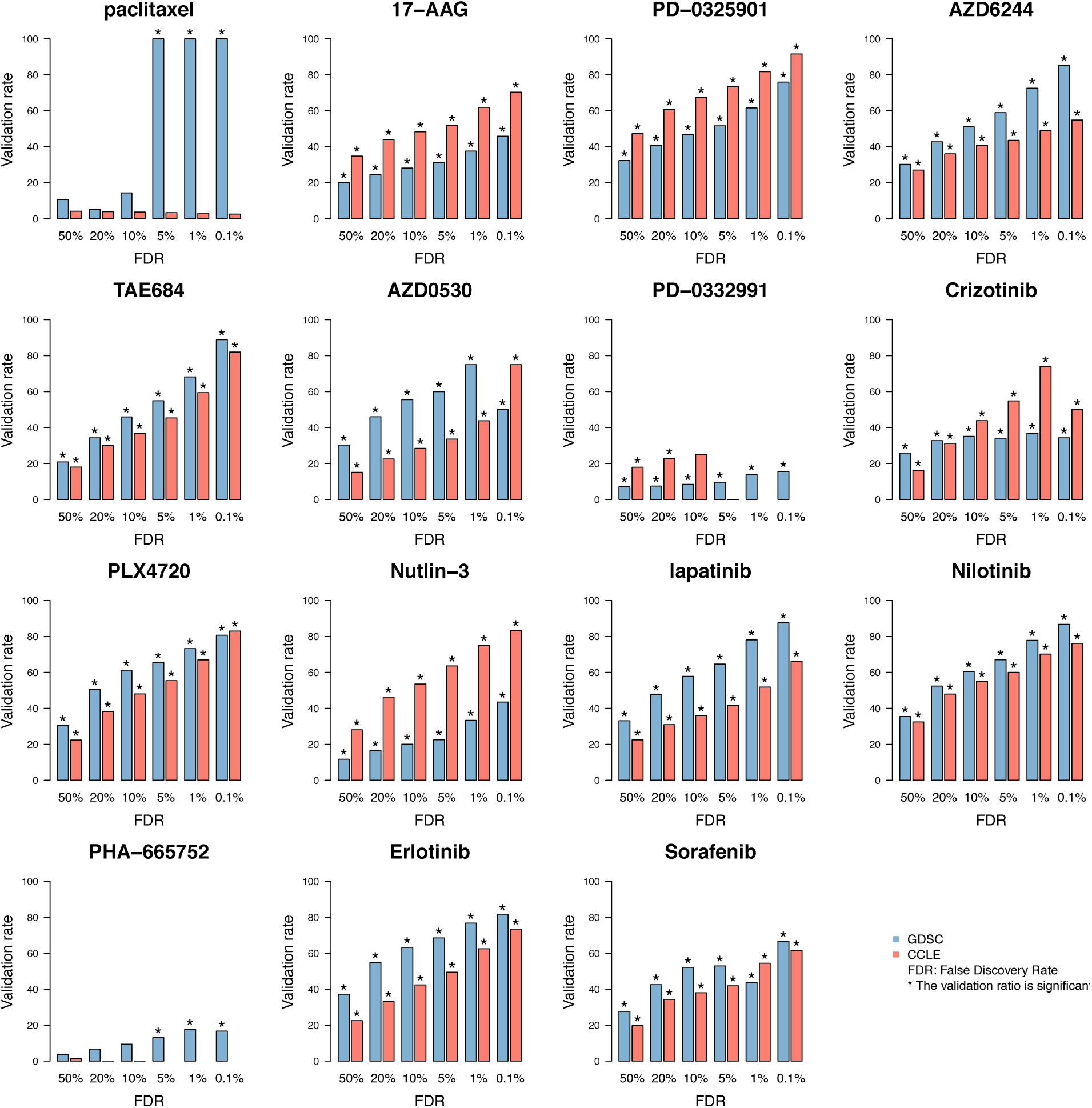
Proportion of validated biomarkers with decreasing FDR using all cell lines in each study. Gene-drug associations are identified using gene expression data and continuous published AUC as input and output of a linear mode, respectively. The symbol ’*’ represents the significance of the proportion of validated gene-drug associations, computed as the frequency of 1000 random subsets of markers of the same size having equal or greater validation rate compared to the observed rate.

## 5 Supplementary Files

**Supplementary File 1.** SNP fingerprints of all the cell lines profiled with SNP arrays in GDSC and CCLE.

**Supplementary File 2.** All the noisy curves identified in GDSC and CCLE.

**Supplementary File 3.** All drug dose-response curves in common between GDSC and CCLE.

**Supplementary File 4.** All drug dose-response curves for replicated experiments using AZD6482 in GDSC.

**Supplementary File 5.** Spreadsheets reporting the statistics (effect size and significance) for all expression-based gene-drug associations for each drug using the common cell lines screened both in GDSC and CCLE. Gene-drug associations were estimated using each gene expression as input and continuous published AUC as output in a linear regression model adjusted for tissue source.

**Supplementary File 6.** Spreadsheets reporting the statistics (effect size and significance) for all expression-based gene-drug associations for each drug using the entire panel of cell lines in GDSC and CCLE. Gene-drug associations were estimated using each gene expression as input and continuous published AUC as output in a linear regression model adjusted for tissue source.

**Supplementary File 7.** Spreadsheets reporting the statistics (effect size and significance) for all mutation-based gene-drug associations for each drug using the entire panel of cell lines in GDSC and CCLE. Gene-drug associations were estimated using each presence indicator of mutations as input and continuous published AUC as output in a linear regression model adjusted for tissue source.

## 6 Supplementary Methods

### PharmacoGx: Structure of the PharmacoSet class

@ 

~~~
annotation
~~~

:

$ 

~~~
name
~~~

: Acronym of the pharmacogenomic dataset.
$ 

~~~
dateCreated
~~~

: When the object was created.
$ 

~~~
sessionInfo
~~~

: Software environment used to create the object.
$ 

~~~
call
~~~

: Set of parameters used to create the object.

@ 

~~~
datasetType
~~~

: Either ‘sensitivity’, ‘perturbation’, or ‘both’

@ 

~~~
cell
~~~

: data frame annotating all cell lines investigated in the study.

@ 

~~~
drug
~~~

: data frame annotating all the drugs investigated in the study.

@ 

~~~
sensitivity
~~~

:

$ 

~~~
n
~~~

: Number of experiments for each cell line treated with a given drug
$ 

~~~
info
~~~

: Metadata for each pharmacological experiment.
$ 

~~~
raw
~~~

: All cell viability measurements at each drug concentration from the drug dose-response curves.
$ 

~~~
phenotype
~~~

: Drug sensitivity values summarizing each dose-response curve (IC_50_, AUC, etc.)

@ 

~~~
perturbation
~~~

:

$ 

~~~
n
~~~

: Number of experiments for each cell line perturbed by a given drug, for each molecular data type
$ 

~~~
info
~~~

: ’The metadata for the perturbation experiments is available for each molecular type by calling the appropriate info function’

@ 

~~~
molecularProfiles
~~~

: List of ExpressionSet objects containing the molecular profiles of the cell lines, such as mutations, gene expressions, or copy number variations.

### SNP fingerprinting

The full pipeline to generate and compare cell line SNP fingerprints is provided below.

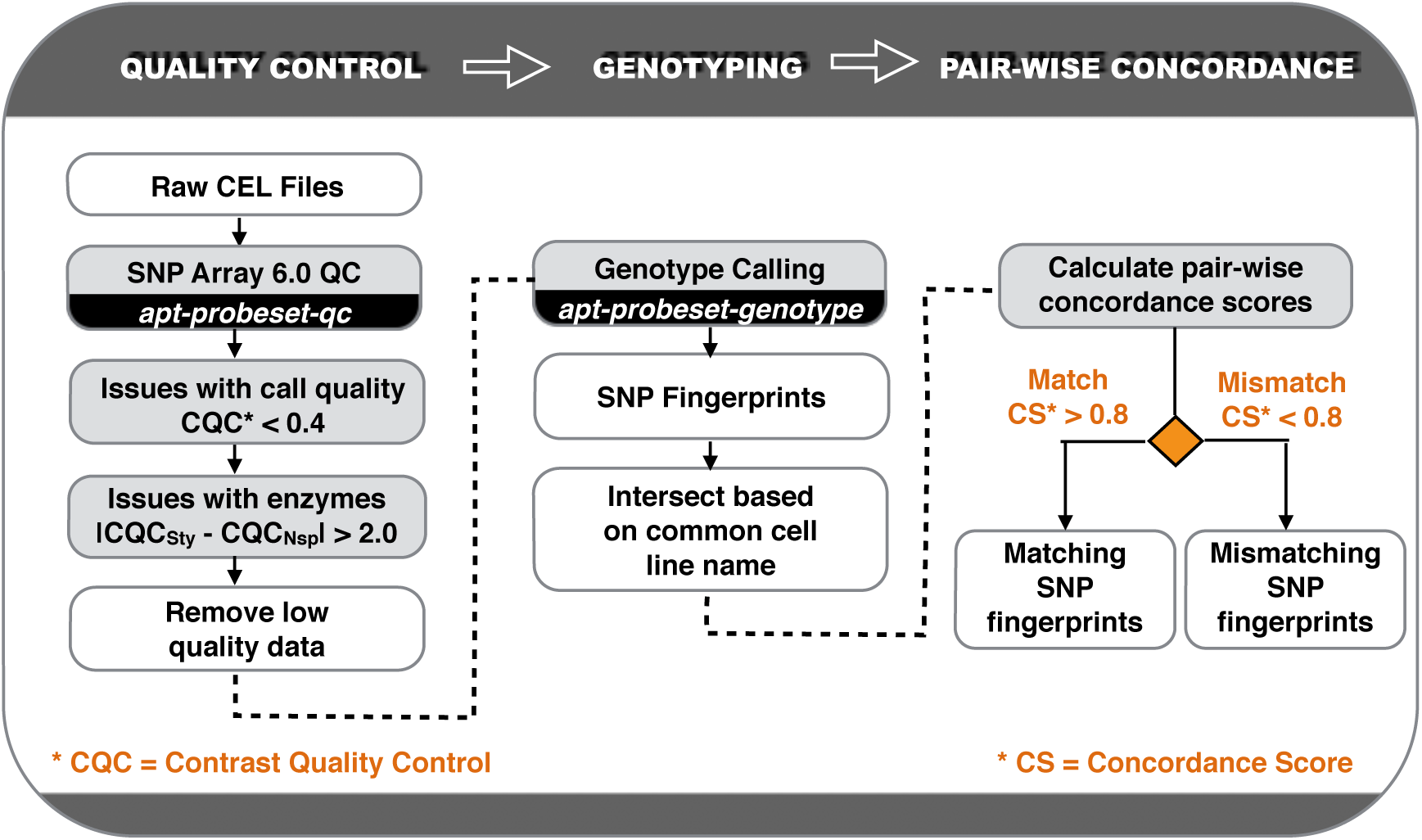

### Affymetrix Power Tools (APT)

Quality control metrics were performed using the apt-probeset-qc program from the Affymetrix Power Tools (Version 1.16.1) suite of tools. Poor quality data was identified using the following criteria outlined via the Affymetrix White Pages: (i) Contrast QC (CQC) sample values less than 0.4; (ii) the proportion of samples for a dataset that falls below 0.4 is greater than 10%; (iii) the mean CQC of all samples in a dataset is less than 1.7; and (iv) the absolute difference between the CQC of Nsp and Sty fragments is greater than 2 18,19. All raw CEL files that failed these metrics were removed from subsequent analysis. The apt-probeset-genotype program was used to call genotypes for the raw CEL files using the birdseed-v2 algorithm and default parameters 20. All remaining files were then intersected based on common cancer cell line names.

### Pair-wise Concordance of SNP Fingerprints

Pairwise concordance scores between all unique SNP fingerprints were calculated using the formula given by Hong et al. [4]:

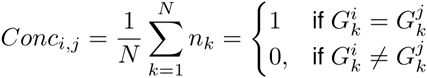

where *N* is the total number of SNPs being compared (909,623 SNPs) and *G* is the genotype for SNP *k* in sample *i* or sample *j*. A concordance score greater than 0.80 was used to indicate consistent genetic identity.

### Filtering of drug dose-response curves

Our quality control approach is based on the assumption that the observed difference Δ_*i,i+1*_ between the cell viability measures in the presence of the *i* +1^*st*^ and *i*^*th*^ highest drug concentration values tested should be less than a small positive threshold ∈ in some large fraction ÏĄ of the cases (1). Unfortunately, setting ∈ small enough to identify all noisy cases in this manner also causes many non-noisy cases to be misidentified as noisy. Consider, for instance, a non-noisy dose response curve that monotonically increases its viability from 99% to 100% over 6 successive drug concentrations, then sees that viability fall monotonically to 20% over the next two concentrations. Consequently, we also required the sum of the Δ_*i*+1,*i*_ to be less than *∈* (2) and the sum of the Δ_*i,j*_ ∀*i,j* to be less than 2*∈* (3).

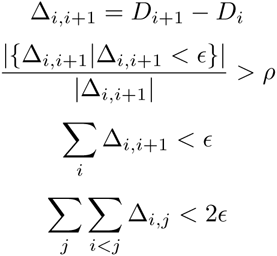

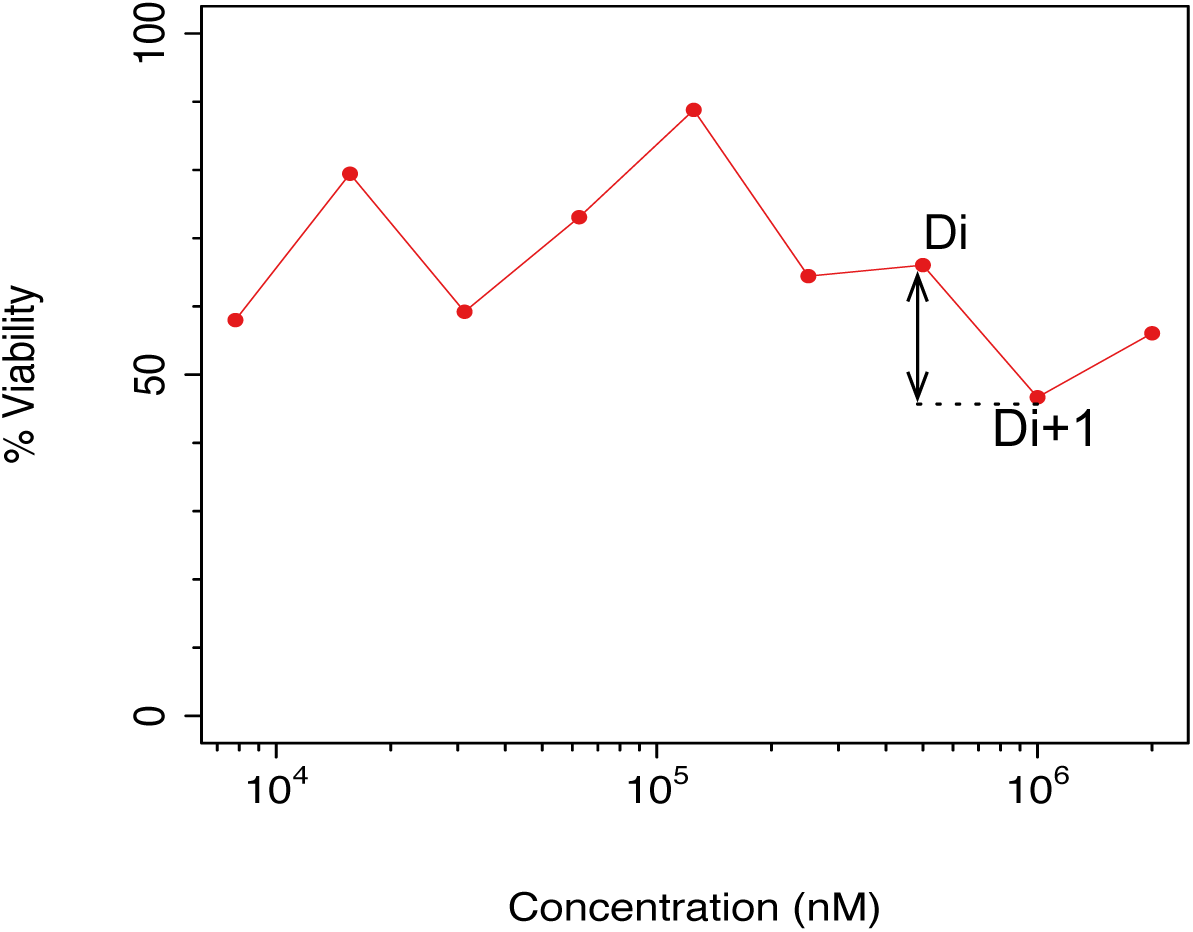

### Fitting of drug dose-response curves

All dose-response curves were fitted to the equation

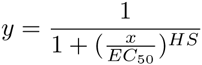

where *y* = 0 denotes death of all treated cells, *y* = *y*(0) = 1 denotes no effect of the drug dose, *EC*_50_ is the concentration at which viability is reduced to half of the viability observed in the presence of an arbitrarily large concentration of drug, and *HS* is a parameter describing the cooperativity of binding. *HS* < 1 denotes negative binding cooperativity, *HS* =1 denotes noncooperative binding, and *HS* > 1 denotes positive binding cooperativity.

The dose-response data in the GDSC and CCLE datasets, as well as the work of Fallahi et al., clearly demonstrate that most drugs are not able to kill all cancerous cells, even at extremely high concentration. We therefore posit a “fractional kill” scenario in which heterogeneous cell lines contain some cells that are resistant to a given drug as well as some that are sensitive to the drug, and only sensitive cells can be killed by the drug. To account for this, we add a third parameter *E* to the model, representing the fraction of resistant cells in the cell line. The dose-response equation now becomes

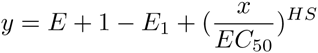

This is the basic mathematical structure that was posited to underlie the dose-response data observed in the study. Consequently, median cellular viability data from all datasets was fit by means of least-squares regression to equations of this type. To ensure robustness of the curve-fitting algorithms, bounds were placed on the values of each of these parameters. Drugs were assumed not to increase the fitness of malignant cells, so *E* was constrained to lie in the interval [0,1]. Drugs were also assumed to be effective in concentration regimes similar to those seen in extant drugs, so *EC*_50_ was assumed to lie in [1pM,1M]. Finally, we follow Fallahi et al. [2] in allowing HS to lie anywhere in [0,4].

Barretina et al. [1] fit dose-response data to one of three models. In most cases, their model of choice was identical to our own, with the addition of a maximum viability parameter *E*_0_. Their dose response equation then became

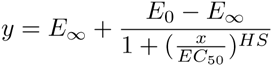

The inclusion of this parameter makes comparison of dose-response curves problematic. With its inclusion, the viability of the cell line in the absence of any drug becomes

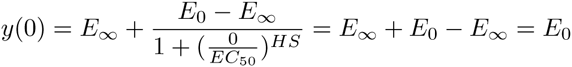

As a result, the viability measures of different drug-cell line combinations are normalized differently, and direct comparison of viability predictions from different dose-response curves is no longer appropriate. The *IC*_50_ values they reported, however, were simply the concentrations at which their fitted curves reached viability reduction of 50% of cellular viability. The end result was a reported *IC*_50_ value that assumed normalization of viability data to the negative control associated with a curve fitted assuming normalization of viability data to a reference level that was most consistent with the observed data. The *IC*_50_ values published in the paper’s supplementary information thus represented viability reduction by a fraction that varied from cell line to cell line.

In GDSC [3], the following five-parameter model was used:

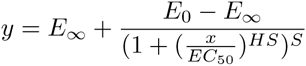

However, since the *E*_0_ parameter is fixed by controls, their curve can be represented as

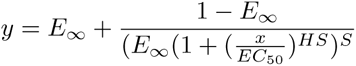

This parameter accounts for the presence of an antagonistic binding of the drug, and introduces asymmetry into the theoretical log dose-response curve. The extra parameter, known as the “Schild slope”, allows the dose-response curve to be non-monotonic.

While this parameter is well-founded biologically, we chose not to use it in our own dose-response curves. As only medians of technical replicates are available for CCLE, using a 4-parameter model would have increased our susceptibility to overfitting noise in the sparse dose-response curves. Furthermore, we only rarely observed the non-monotonicity that necessitates the inclusion of a Schild slope parameter in a very small fraction of dose-response curves. For these reasons, we ultimately chose to use our simpler 3-parameter model to compare the dose-response curves from the GDSC and CCLE datasets.

### Area between the drug dose-response curves (ABC)

The ABC is a function of two dose-response curves from CCLE and GDSC which come from the same drug-cell line combination. It is calculated by taking the unsigned area between the two curves over the intersection of the concentration range tested in the CCLE curve and that tested in the GDSC curve, and normalizing that area by the length of the intersection interval.

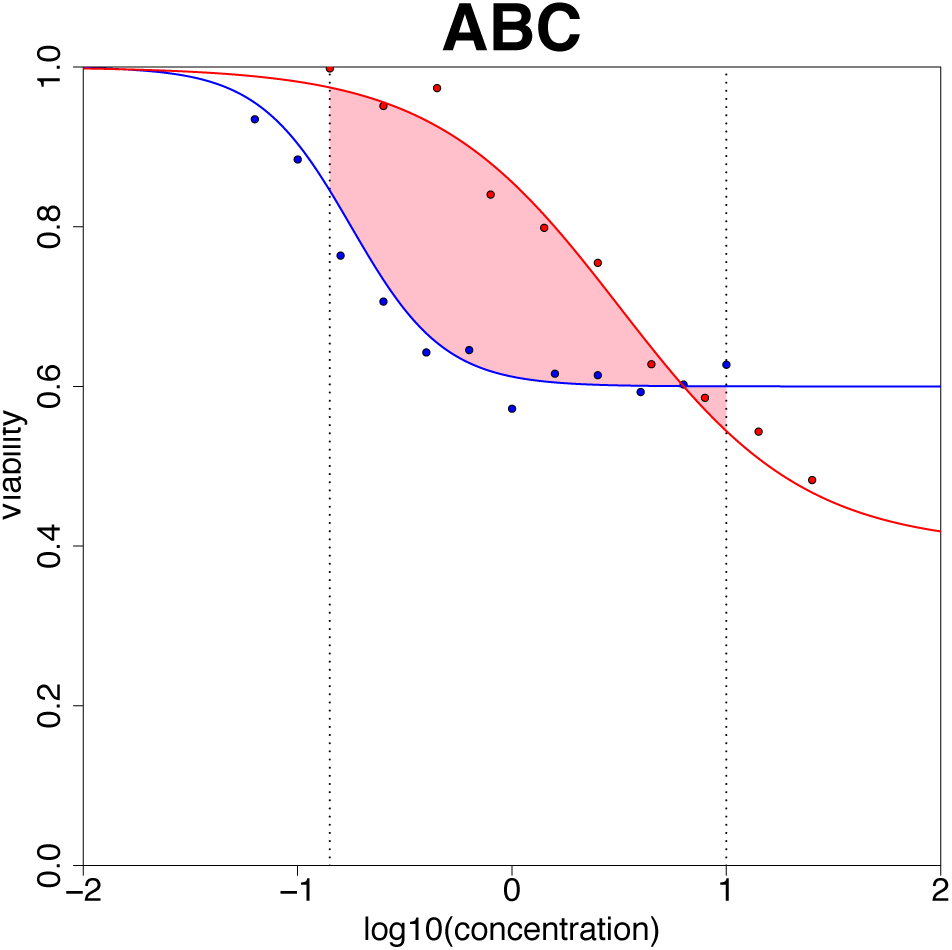

### Consistency *across* vs. *between* cell lines

We assessed the concordance of the gene expression, mutation and drug sensitivity of CGP and CCLE studies *across* and *between* cell lines, as illustrated in the figure below. When data are compared across cell lines, we assess whether, for a given gene expression or drug, the cell line data were concordant (a gene is expressed at a similar level or similar response to a drug is observed in the same set of cell lines for instance; panel **A**). When data are compared between cell lines, we assessed whether, for a given cell line, the genomic and pharmacological profiles were concordant in the two studies (a given cell line harbours similar gene expression patterns or pharmacological responses for instance; panel **B**).

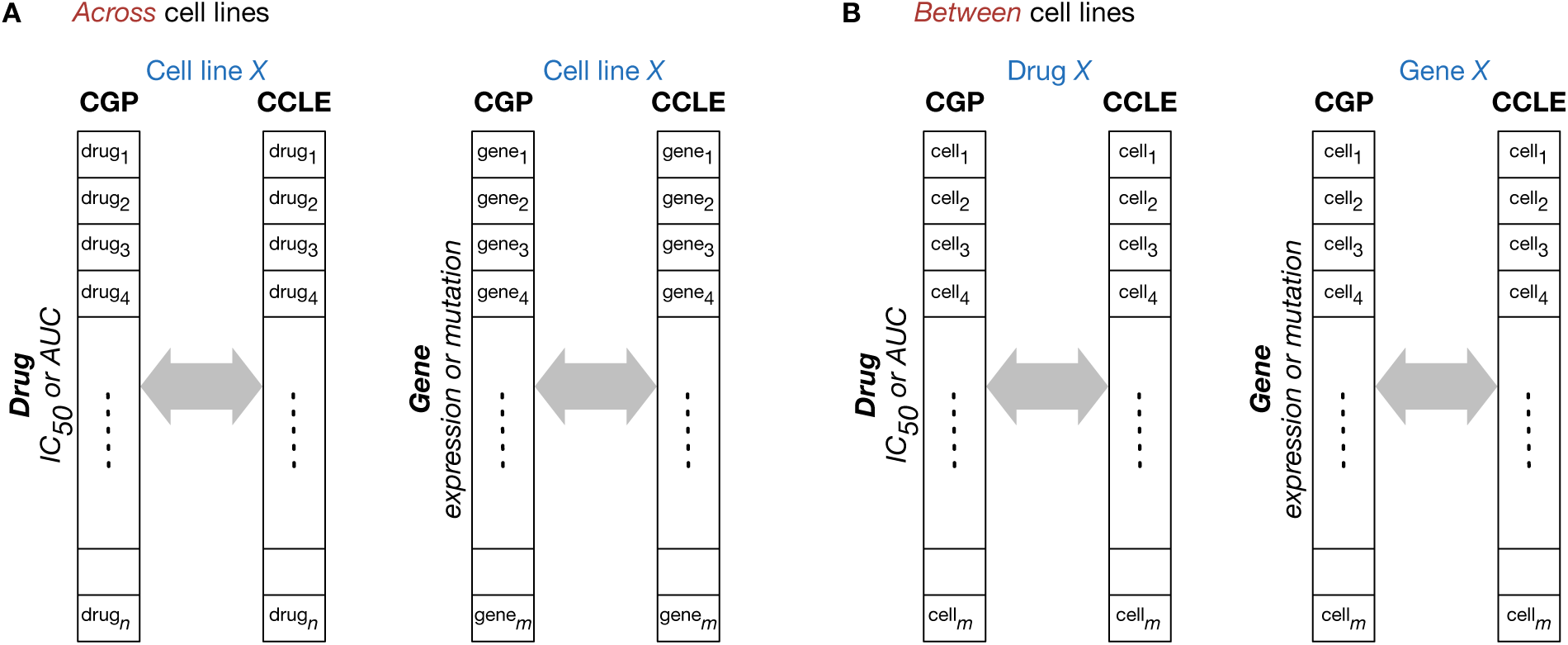

### 7 Acronyms

ABC: Area between the curves
AE: ArrayExpress by the European Bioinformatics Institute
AUC: Area under the dose response curve
AUC* or STAR: Area under the dose response curve calculated by considering only the common concentration range between
CCLE: The Cancer Cell Line Encyclopedia initiated by the Broad Institute of MIT and Harvard
CGHub: The Cancer Genomics Hub from the University of California Santa Cruz and the US National Cancer Institute
CGP: The Cancer Genome Project by the Wellcome Trust Sanger Institute
CMAP: Connectivity Map by the Broad Institute
COSMIC: Catalogue of Somatic Mutations in Cancer by the Wellcome Trust Sanger Institute
CRAMERV: Carmer’s V
DXY: Somers’ Dxy rank correlation
GDSC: The Cancer Genome Project initiated by the Wellcome Trust Sanger Institute
IC_50_: Concentration at which the drug inhibited 50% of the maximum cellular growth
INFORM: Informedness
MCC: Matthews correlation coefficient
PCC: Pearson product-moment correlation coefficient
QC: Quality control
RMA: Robust multi-array normalization
SCC: Spearman rank correlation coefficient
SNP: Single nucleotide polymorphism

